# Discovery of a primed endothelial progenitor that requires VEGF/ERK inhibition to complete vein differentiation

**DOI:** 10.1101/2025.10.11.681838

**Authors:** Lay Teng Ang, Sherry Li Zheng, Kevin J. Liu, Anastasiia Masaltseva, June Winters, Sawan K. Jha, Qingqing Yin, Crystal Qian, Xiaochen Xiong, Amir Dailamy, Ellie Xi, Juan C. Alcocer, Daniel W. Sorensen, Richard She, Karina Smolyar, Dorota Szumska, Svanhild Nornes, Renata M. Martin, Benjamin J. Lesch, Nicole K. Restrepo, Wenfei Sun, Jonathan S. Weissman, Heiko Lickert, Matthew H. Porteus, Mark A. Skylar-Scott, Christian Mosimann, Saulius Sumanas, Sarah De Val, Kristy Red-Horse, Kyle M. Loh

**Affiliations:** Institute for Stem Cell Biology & Regenerative Medicine, Stanford University, Stanford, CA, USA; Department of Urology, Stanford University, Stanford, CA, USA; Department of Developmental Biology, Stanford University, Stanford, CA, USA; Department of Bioengineering, Stanford University, Stanford, CA, USA; Department of Biology, Howard Hughes Medical Institute, Stanford University, Stanford, CA, USA; School of Biological Sciences, Nanyang Technological University, Singapore; Whitehead Institute for Biomedical Research, Department of Biology, Howard Hughes Medical Institute, Massachusetts Institute of Technology, Cambridge, MA, USA; Institute of Developmental and Regenerative Medicine, Department of Physiology, Anatomy and Genetics, University of Oxford, Oxford, United Kingdom; Department of Pediatrics, Stanford University, Stanford, CA, USA; Department of Pathology and Cell Biology, University of South Florida, Tampa, Florida, USA; Division of Endocrinology, Department of Medicine, Stanford University, Stanford, CA, USA; Institute of Diabetes and Regeneration Research, Helmholtz Center Munich, Neuherberg, Germany; German Center for Diabetes Research, Neuherberg, Germany; School of Medicine, Technical University of Munich, Munich, Germany; Basic Science and Engineering Initiative, Children’s Heart Center, Chan Zuckerberg Biohub, Stanford University, Stanford, CA, USA; Section of Developmental Biology, Department of Pediatrics, University of Colorado School of Medicine, Anschutz Medical Campus, Aurora, CO, USA; Ludwig Institute for Cancer Research Ltd, Nuffield Department of Medicine, University of Oxford, Oxford, United Kingdom

## Abstract

Extracellular signals and cell-fate trajectories during vein development remain elusive, despite trailblazing insights into artery development. Here we exploit human pluripotent stem cell differentiation and mouse embryology to present a model that answers longstanding questions: vein endothelial cell (EC) differentiation unfolds in two steps driven by opposing extracellular signals. First, VEGF differentiates mesoderm into “primed” ECs, newly-defined progenitors that co-express certain arterial (SOX17) and venous (APLNR) markers. Second, primed ECs execute vein differentiation upon VEGF/ERK inhibition; however, upon VEGF activation they can instead form artery ECs. The arteriovenous plasticity of primed ECs was supported by intersectional lineage tracing. Future venous genes including *NR2F2* harbor poised chromatin in primed ECs, but are only transcribed upon VEGF/ERK inhibition. SOXF transcription factors, including SOX17, confer primed ECs with vein differentiation competence. Collectively, this two-step vein differentiation model—entailing primed EC intermediates and VEGF/ERK inhibition to trigger vein differentiation—has implications for VEGF-modulating therapies.

## INTRODUCTION

Endothelial cells (ECs) comprise the inner lining of blood vessels, and there are multiple subtypes of ECs, including artery, vein, capillary, and lymphatic ECs; each executes specialized functions and expresses different genes^1-10^. For instance, artery ECs express NOTCH ligands that specify the smooth muscle cells that ensconce arteries^11^. By contrast, vein ECs can construct valves that prevent backward blood flow^12^. The prevailing model suggests that artery ECs express SOXF family transcription factors (SOX7, SOX17, and SOX18) that impart arterial identity^13-19^. Conversely, vein ECs express the transcription factor NR2F2 and transmembrane proteins APLNR/APJ and FLRT2^20-26^.

Pathbreaking work has discovered key precepts of artery EC development^3-10^: VEGF/ERK activation^27-29^, NOTCH activation^30-34^, and cell-cycle arrest^23,35,36^ all drive arterialization. Nevertheless, many mysteries continue to surround the development of vein ECs^6,8^. For instance, one might assume that arteries and veins should arise contemporaneously, as both are required to form a functional circulatory system. Nevertheless, during the development of the first intraembryonic blood vessels, arteries emerge earlier than veins, as shown by anatomy and molecular markers in mouse and zebrafish embryos^24,37-42^. Why veins should emerge after arteries remains a longstanding curiosity. This piques questions about the earliest steps of venous development and the exact identity of the progenitor cells that build veins.

The extracellular signals that specify venous identity also warrant further investigation^6,8^. The transcription factor NR2F2 is a master regulator of vein EC identity^22^, but the upstream signals that ignite NR2F2 expression remain largely unknown^6,8^. VEGF is a master regulator of EC identity^4,43-47^, but has paradoxical roles in venous development. On one hand, VEGF is *required* for vein EC formation, as ECs are absent in *Vegfr2^-/-^* mouse embryos^45^. On the other hand, VEGF also *represses* vein formation, as it instructs arterial identity in zebrafish embryos^27,28^. How can VEGF both promote and inhibit vein EC specification during vascular development?

Here we introduce a conceptual model of human vein EC differentiation that addresses certain mysteries surrounding vein development. We unveil two separable steps of vein differentiation, driven by opposing extracellular signals. First, VEGF *activation* differentiates lateral mesoderm into a newly discovered “primed” EC intermediate, which co-expresses “arterial” marker SOX17 and “venous” marker APLNR. Primed ECs harbor the competence to generate both vein and artery ECs *in vitro*, consistent with broad arteriovenous plasticity in the vascular system^10^. Intersectional genetic tracing supports the notion that *Sox17*+ *Aplnr*+ ECs form both vein and artery ECs *in vivo*. Within primed ECs, future venous and arterial genes are poised at the chromatin level, but subsequently VEGF/ERK *inhibition* installs fully-fledged venous identity, and converts these poised venous genes into an active state.

The discovery of primed ECs may explain several longstanding curiosities in vascular development, including why veins tend to emerge after arteries *in vivo*^24,37-42^. Our model also reconciles why VEGF/ERK signaling can paradoxically both promote^45^ and inhibit^27-29^ vein EC development: the same signal executes both functions, at different times. The timing of an extracellular signal is of paramount importance, as it is interpreted in a temporally dynamic way. Our discovery that VEGF/ERK inhibition is required for vein EC specification also has potential implications for VEGF pathway inhibitors that are clinically used to block tumor angiogenesis^46,47^.

## RESULTS

### Discovery of “primed” ECs that arise during human vascular differentiation *in vitro*

At 24 hour increments during hPSC differentiation into artery and vein ECs^48^, we mapped gene expression (through single-cell [scRNAseq] and bulk-population RNA-sequencing), chromatin accessibility (using OmniATACseq^49^), and cell-surface marker expression (through high-throughput screening of 332 cell-surface markers) (**Fig. 1A**, **Fig. S1A-E**, **Table S1-S2**). We created an interactive web browser to explore scRNAseq data of each differentiation stage (https://anglab.shinyapps.io/artery-vein-scrna-seq/).

**Figure 1:**
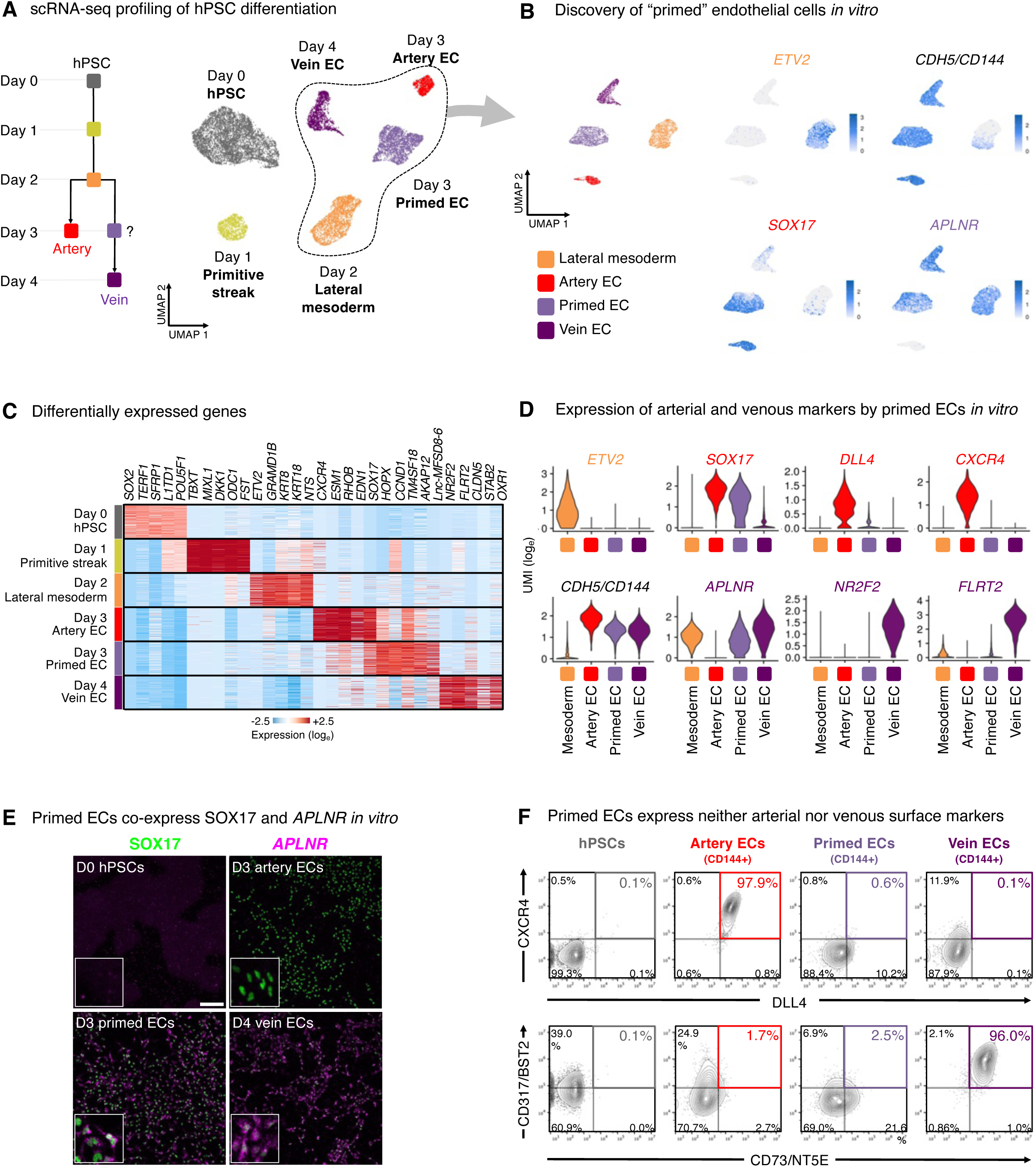
A roadmap for human arteriovenous differentiation reveals primed endothelial cells. A) scRNA-seq of day-0 H1 hPSCs, day-1 primitive streak, day-2 lateral mesoderm, CD144+ FACS-purified day-3 artery ECs, CD144+ FACS-purified day-3 primed ECs, and CD144+ FACS-purified day-4 vein ECs. B) Differentially expressed genes that distinguish each hPSC-derived cell-type, as detected by scRNAseq. scRNA-seq was performed on day-0 H1 hPSCs, day-1 primitive streak, day-2 lateral mesoderm, CD144+ FACS-purified day-3 artery ECs, CD144+ FACS-purified day-3 primed ECs, and CD144+ FACS-purified day-4 vein ECs. C) scRNA-seq of day-2 lateral mesoderm, CD144+ FACS-purified day-3 artery ECs, CD144+ FACS-purified day-3 primed ECs, and CD144+ FACS-purified day-4 vein ECs. D) scRNA-seq of day-2 lateral mesoderm, CD144+ FACS-purified day-3 artery ECs, CD144+ FACS-purified day-3 primed ECs, and CD144+ FACS-purified day-4 vein ECs. Gene expression is depicted in log_e_ unique molecular identifier (UMI) counts. E) Combined immunostaining for SOX17 protein and HCR3 *in situ* hybridization for *APLNR* mRNA in the indicated H1 hPSC-derived cell-types. Scale: 200 μm. F) Flow cytometry of H1 CRISPRi-expressing hPSCs, day-3 artery ECs, day-3 primed ECs, and day-4 vein ECs. Day 3-4 populations were pre-gated on the CD144+ EC subset before depicting marker expression.

Investigating stepwise molecular changes during venous differentiation in detail, we discovered an unexpected intermediate with unique gene expression, chromatin, and cell-surface marker signatures—which we term “primed” ECs—that preceded the emergence of vein ECs. Day-2 lateral mesoderm expressed the angioblast transcription factor *ETV2*^50^, but upon 24 hours of further differentiation, *ETV2* was silenced, whereupon EC surface markers (e.g., *CD144/VE-CADHERIN*) became expressed by day 3 (**Fig. 1B-D**, **Fig. S2A-B**). However, mature venous markers *NR2F2* and *FLRT2* were only expressed by day 4 of vein differentiation (**Fig. 1C-D, Fig. S2A-B**).

What is the identity of these incipient day-3 “primed” ECs emerging prior to expression of fully-fledged venous markers by day 4? Curiously, CD144+ primed ECs co-expressed the “arterial” marker *SOX17*^13-17^ and “venous” marker *APLNR*^20,21^ (**Fig. 1B,D**). By contrast, cells subjected to arterial differentiation solely expressed *SOX17*, but not *APLNR*, as expected^13-17,20-26^ (**Fig. 1B,D**). We confirmed that single primed ECs co-expressed SOX17 and *APLNR*, as shown by combined *in situ* hybridization and immunostaining (**Fig. 1E**) and scRNAseq (**Fig. S2C**). However, primed ECs did not express additional markers of arterial (*GJA4/CX37*, *UNC5B*, *DLL4, MECOM, HEY1, EFNB2, CXCR4*, or *IGFBP3*), venous (*NR2F2*, *FLRT2*, or *NRP2*), or angioblast (*ETV2*) identity^23,24^ (**Fig. 1D**, **Fig. S2B,D**). High-throughput profiling of cell-surface markers revealed that primed ECs did not express markers of artery ECs (CXCR4+ DLL4+) or vein ECs (CD73+ CD317+)^51-53^, suggesting that primed ECs can be classified as neither arterial nor venous (**Fig. S3A-F**, **Table S3**). Having discovered primed ECs, we subsequently explored the extracellular signals that drive entry into, and exit from, this intermediate state.

### Two steps of vein EC differentiation *in vitro*: VEGF/ERK activation, followed by inhibition

What extracellular signals specify venous identity? We tested different combinations of the developmental signals shown in **Fig. 2A**. This revealed that VEGF *activation* was required to differentiate day-2 lateral mesoderm into day-3 primed ECs, alongside inhibition of the artery-specifying TGFϕ3 and NOTCH pathways^30-34,48^ (**Fig. 2B**). During this first 24-hour interval, VEGF/ERK activation was crucial to generate ECs; in its absence, EC formation *in vitro* was significantly impaired (**Fig. 2B**, **Fig. S4A**). The requirement for VEGF to differentiate lateral mesoderm into primed ECs is consistent with how VEGF is required for EC emergence *in vivo*^27,45^.

**Figure 2:**
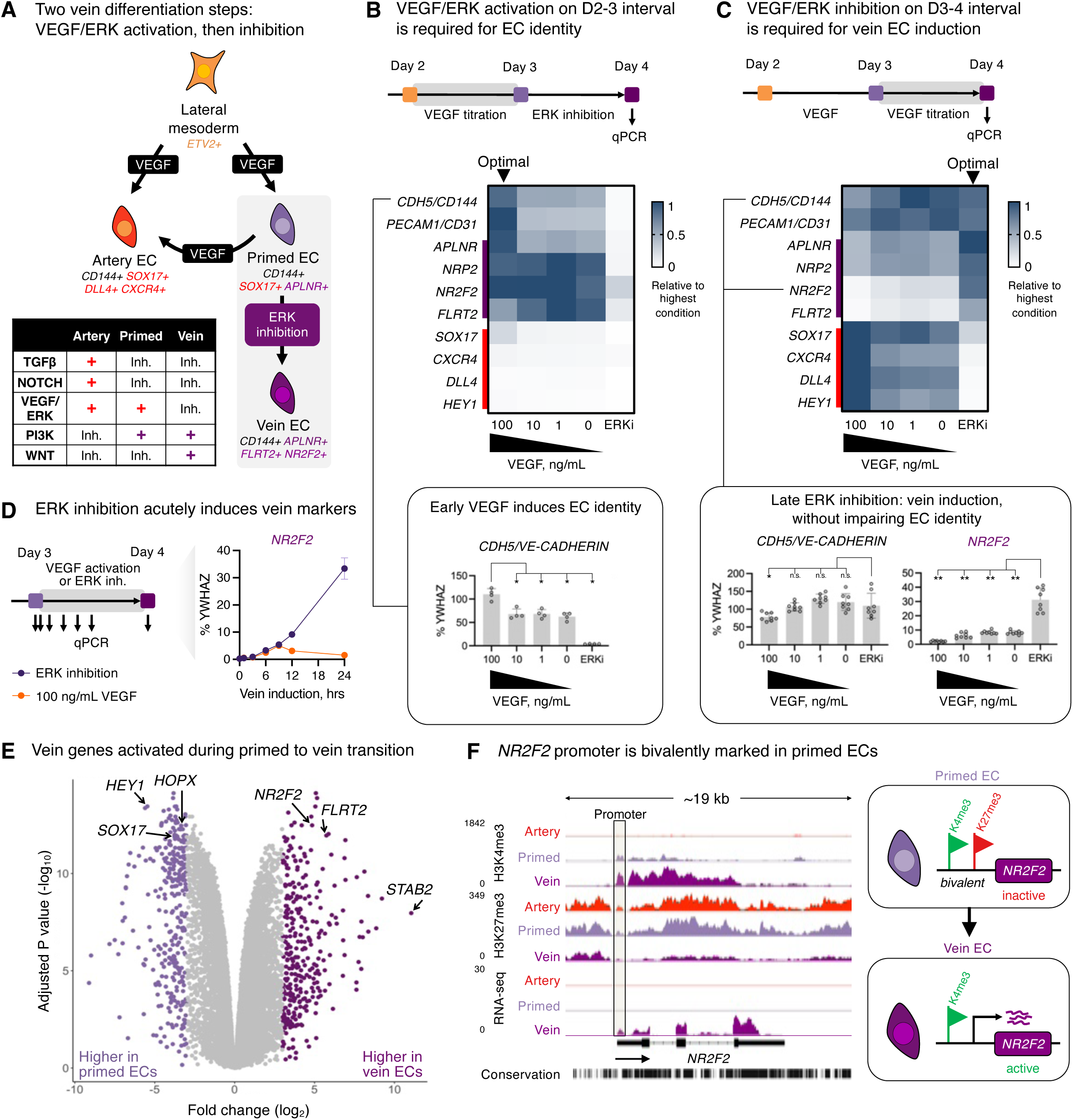
Two separable steps of vein differentiation driven by temporally dynamic VEGF/ERK activation, followed by inhibition. A) Summary of the present study. B) First, H1 hPSCs were differentiated into day-2 lateral mesoderm^48^. Then, lateral mesoderm was then treated with different doses of VEGF pathway modulators (VEGF [0-100 ng/mL] or ERK inhibitor [ERKi; PD0325901, 500 nM]) alongside other primed EC-inducing signals (TGFϕ3 inhibitor + NOTCH inhibitor + BMP inhibitor + WNT inhibitor + Vitamin C) for 24 hours. Subsequently, cells were subject to vein EC differentiation for 24 hours. qPCR was conducted on day 4 of hPSC differentiation. In heatmaps, expression is normalized to the sample with the highest expression in either panel B or C. This revealed that high VEGF for 24 hours is required during primed EC specification to subsequently generate vein ECs by day 4. Statistics: Wilcoxon rank sum test. Error bars: standard deviation (SD). *P<0.05. C) First, H1 hPSCs were differentiated into day-3 primed ECs. Then, primed ECs were then treated with different doses of VEGF pathway modulators (VEGF [0-100 ng/mL] or ERK inhibitor [ERKi; PD0325901, 500 nM]) alongside other vein EC-inducing signals (TGFϕ3 inhibitor + NOTCH inhibitor + WNT agonist + Vitamin C) for 24 hours. qPCR was conducted on day 4 of hPSC differentiation. In heatmaps, expression is normalized to the sample with the highest expression in either panel B or C. This revealed that ERK inhibition for 24 hours is required to generate day-4 vein ECs, and that after cells acquire endothelial identity, VEGF/ERK is dispensable for the continued expression of pan-EC markers. Statistics: Wilcoxon rank sum test. Error bars: SD. **P<0.01, *P<0.05, n.s.: not significant. D) First, H1 hPSCs were differentiated into day-3 primed ECs. Then, primed ECs were then treated with either VEGF (100 ng/mL) or ERK inhibitor (PD0325901, 500 nM) alongside other vein EC-inducing signals (TGFϕ3 inhibitor + NOTCH inhibitor + WNT agonist + Vitamin C) for 1-24 hours, followed by qPCR. This revealed that ERK inhibition for 12 hours significantly upregulated venous marker expression. E) Bulk-population RNA-seq of FACS-purified CD144+ day-3 primed ECs and day 4-vein ECs generated from H1 hPSCs. Differentially expressed genes are colored purple. F) CUT&RUN profiling of H3K4me3 and H3K27me3 and bulk RNA-seq H1 hPSC-derived day-3 artery ECs, day-3 primed ECs, and day-4 vein ECs.

While VEGF was essential to generate day-3 primed ECs, our combinatorial testing of developmental signals unexpectedly revealed that VEGF/ERK *inhibition* was necessary for their further progression into day-4 vein ECs (**Fig. 2C**, **Fig. S4B**). Upon VEGF/ERK inhibition, venous markers *NR2F2* and *FLRT2* sharply increased, and primed EC marker *SOX17* was silenced (**Fig. 2D-E**, **Fig. S2A**). 12 hours of VEGF/ERK inhibition sufficed to significantly activate *NR2F2* expression (**Fig. 2D**). By contrast, continued VEGF activation at this stage was arterializing, and repressed venous markers (**Fig. 2C-D**), as observed in zebrafish embryos^27,28^. The requirement for VEGF/ERK inhibition to confer venous identity was surprising, as VEGF is typically thought to be broadly required for all ECs^4,27,45^. Nevertheless, after cells have acquired endothelial identity, we found that VEGF was dispensable for cells to remain endothelial (**Fig. 2C**).

While primed ECs differentiated into vein ECs upon VEGF/ERK inhibition, they could instead adopt artery EC identity if challenged with artery-inducing signals, such as VEGF activation and PI3K inhibition^48,54^ (**Fig. S4C**). The competence of primed ECs to form both vein and artery ECs *in vitro* reflects known arteriovenous plasticity within the vasculature *in vivo*^10^.

Given that primed ECs harbor vein differentiation competence, we hypothesized that venous genes might be poised for future activation at the chromatin level. To test this hypothesis, we mapped the genome-wide distribution of H3K4me1, H3K4me3, and H3K27me3 using CUT&RUN^55^. In primed ECs, future venous genes such as *NR2F2*, *NT5E*, and *MAF* were not yet expressed, but nevertheless their promoter regions were bivalently marked by activation-associated H3K4me3 and repression-associated H3K27me3 (**Fig. 2F**, **Fig. S4D**). Such promoter bivalency can denote genes poised for future activation in anticipation of future developmental decisions^56^. VEGF/ERK inhibition transitioned the promoter elements of these venous genes from bivalency (H3K4me3+ H3K27me3+) to full activation (H3K4me3+ H3K27me3-), which was accompanied by transcription (**Fig. 2F**, **Fig. S4D**). Likewise, the promoter elements of future arterial genes were bivalently marked in primed ECs, foreshadowing arterial differentiation competence (**Fig. S4E**).

We thus propose two sequential steps of vein differentiation driven by temporally dynamic VEGF *activation* to specify primed ECs, followed by VEGF/ERK *inhibition* to induce vein ECs (**Fig. 2A**). This temporally-dynamic signaling switch reconciles why VEGF/ERK signaling has been reported to both promote^45^ and inhibit^27-29^ vein formation *in vivo*: the same signal can do both, at different times.

### Comparison of differentiation methods suggests that VEGF/ERK inhibition is integral to venous identity

Despite widespread success in differentiating hPSCs towards artery ECs, it has generally proven more challenging to generate vein ECs^6^. We hypothesized that difficulties in generating vein ECs might be attributable to the temporally-dynamic effects of VEGF: current differentiation methods typically entail prolonged VEGF activation^51-53,57-59^, which might inadvertently deter venous differentiation. To this end, we compared 8 hPSC differentiation methods by collating published scRNAseq datasets and generating new ones^48,58-62^ (**Fig. 3A**, **Fig. S5A-B**). What is the diversity and arteriovenous character of cells emerging from each protocol?

**Figure 3:**
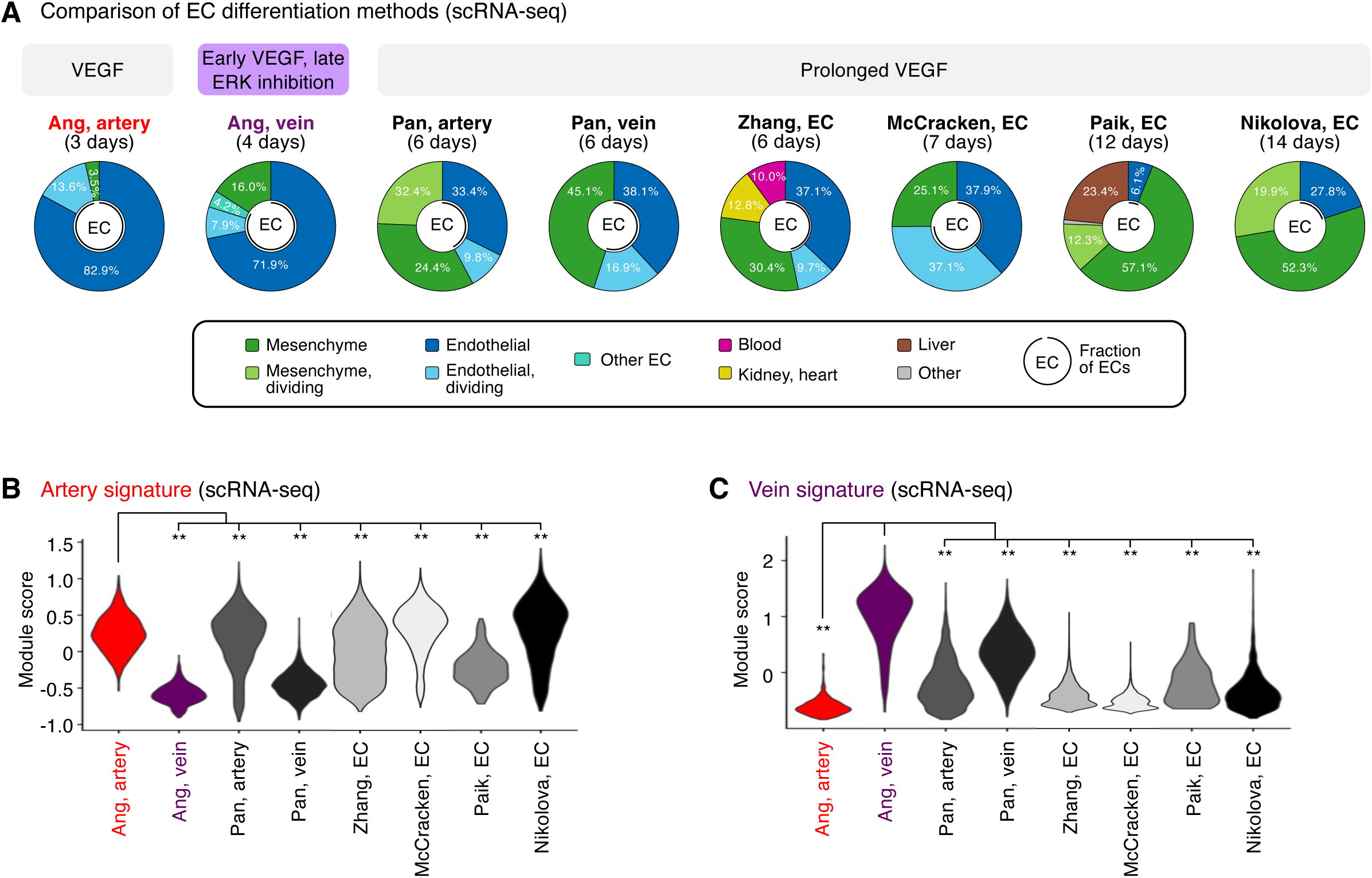
Comparison of differentiation methods suggests that temporally dynamic VEGF modulation is crucial to impart venous identity. A) scRNA-seq of hPSCs differentiated into ECs using various protocols, with the number of days of differentiation indicated. scRNA-seq datasets were subclustered at the same resolution to identify population heterogeneity. Clusters were annotated by marker expression. The proportion of ECs is indicated. scRNA-seq data were obtained from this study, Ang et al., 2022^48^, Pan et al., 2024^58^, McCracken et al., 2019^62^, Paik et al., 2018^60^, and Nikolova et al., 2025^59^. B) scRNA-seq of differentiated hPSC populations described in Fig. 3A, and ECs were computationally isolated. An expression module score^93^ of arterial markers defined *in vivo* by Hou et al., 2022^24^ is shown. Statistics: Wilcoxon rank sum test. **P<0.01. C) scRNA-seq of differentiated hPSC populations described in Fig. 3A, and ECs were computationally isolated. An expression module score^93^ of venous markers defined *in vivo* by Hou et al., 2022^24^ is shown. Statistics: Wilcoxon rank sum test. **P<0.01.

All 8 differentiation methods generated ECs and mesenchymal cells in varying proportions. The two differentiation protocols employed in this study^48^ yielded the highest EC percentages (84.0-96.5%), with the remaining non-ECs comprising mesenchymal cells (**Fig. 3A**, **Fig. S5C**, **Fig. S6**). Certain differentiation protocols^60,61^ yielded a wider spectrum of non-ECs, including blood, kidney, heart, and liver (**Fig. 3A**, **Fig. S5C**, **Fig. S6**). To minimize the emergence of non-ECs, we developed improved protocols to generate artery and vein ECs at higher purity and scale, yielding up to 654 million artery or vein ECs in a single batch (**Fig. S7A-F**).

Next, we quantified the arteriovenous character of ECs emerging from each *in vitro* differentiation protocol, using *in vivo*-defined arteriovenous marker signatures^24^. Multiple protocols generated artery ECs—perhaps reflecting the pervasive *in vitro* use of VEGF (**Fig. S5B**), which is arterializing *in vivo*^27,28^—although the arterial marker score differed between protocols (**Fig. 3B**, **Fig. S6**). The artery EC differentiation protocol employed in this study^48^ imparted the strongest arterial identity (**Fig. 3B**; P<0.0001), potentially attributable to the explicit inhibition of vein-inducing signal PI3K^29,63^ to sharpen arterial identity. Only two differentiation protocols^48,58^ predominantly generated vein ECs, consistent with how it has proven more challenging to instate venous identity *in vitro*^6^. The protocol employed in this study^48^—which conferred the strongest venous identity (**Fig. 3C**; P<0.0001)—entailed VEGF-induced generation of primed ECs, followed by VEGF/ERK inhibition to specify vein ECs. Other methods relying on sustained VEGF activation^58^ less effectively induced venous character (**Fig. 3C**, **Fig. S6**). After generating vein ECs, we developed enhanced methods to expand them for at least 6 days *in vitro* while preserving venous identity (**Fig. S7G-L**).

In summary, this comparison of differentiation protocols suggests that (1) transition through primed EC intermediates and (2) temporally dynamic VEGF/ERK modulation are critical to maximize vein EC generation *in vitro*. Prolonged VEGF activation is the cornerstone of extant EC differentiation protocols (**Fig. S5B**), but temporally dynamic VEGF/ERK activation, followed by inhibition, may prove decisive in imparting venous identity *in vitro*.

### Sox17+ Aplnr+ primed ECs exist in early embryos and contribute to veins

Having discovered *SOX17*+ *APLNR*+ primed ECs *in vitro*, we investigated their existence and fate *in vivo*. Sox17 marks arteries *in vivo*^13-17^, but others suggest that it is more broadly expressed in the vasculature^64,65^. Whole mount immunostaining revealed that Sox17 marked most, if not all, Erg+ ECs in the E8.5 mouse embryo, including those in the dorsal aorta and the vitelline vein (**Fig. 4A-B**, **Fig. S8A**). In E9.5 mouse embryos, Sox17 was expressed by both the dorsal aorta and cardinal vein (**Fig. 4C**). Indeed, permanent labeling of *Sox17*+ cells and all of their progeny with *Sox17*-*Cre* resulted in both artery and vein ECs being lineage traced (**Fig. S8B-C**), consistent with past observations^66,67^. Despite the prominent role of Sox17 in artery development^13-17^, these lineage tracing and immunostaining results demonstrate that early vein ECs also transiently express Sox17, although it is turned off later in the mature cardinal vein by E10.5^16,17^. Sox17 is thus expressed by most ECs in early vascular development, and these cells construct most of the nascent vasculature.

**Figure 4:**
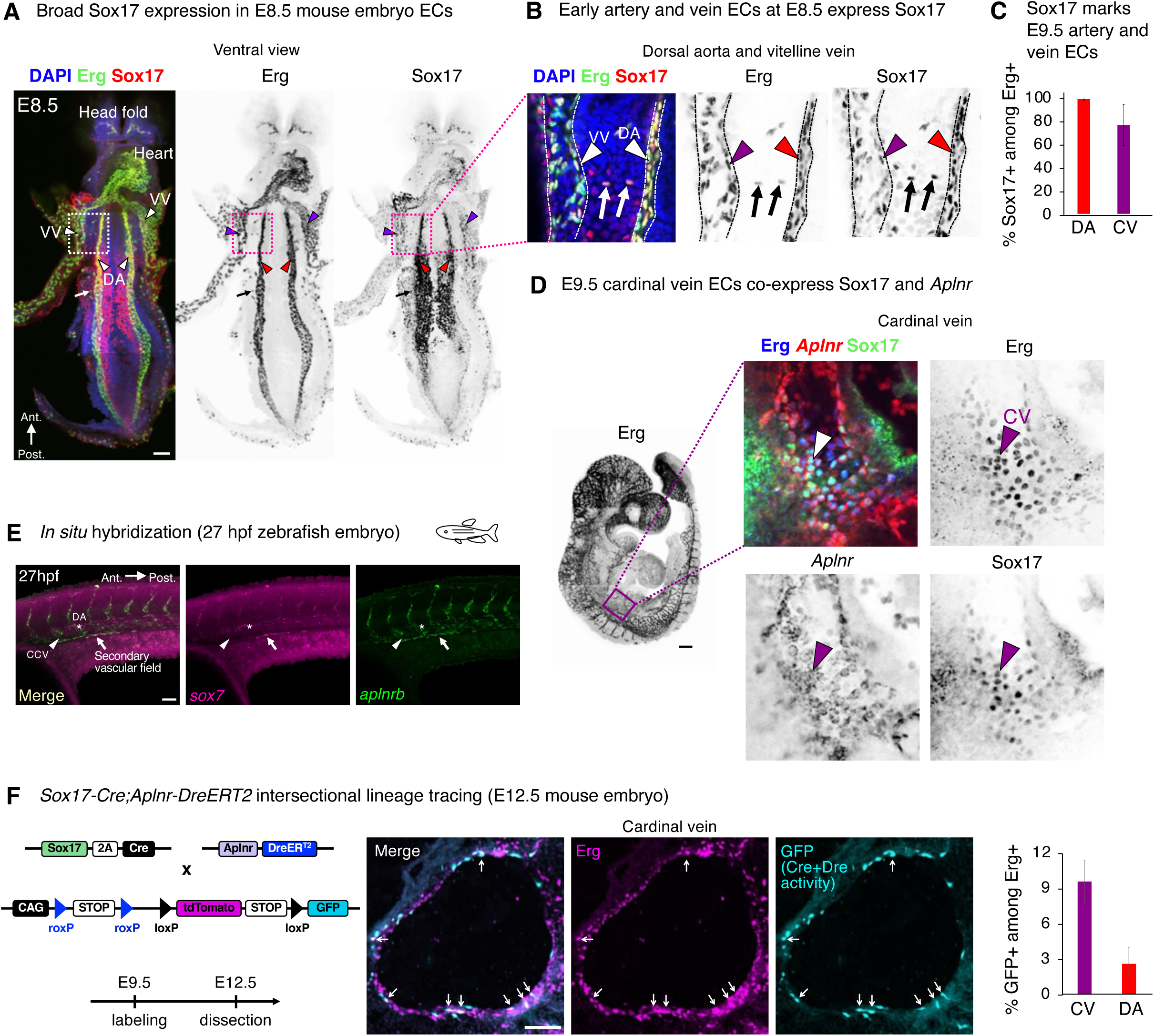
*Sox17*+ *Aplnr*+ endothelial cells exist in the early embryo, and contribute to both veins and arteries. A) Whole-mount Erg and Sox17 immunostaining of E8.5 mouse embryo. Ant: anterior. Post: posterior. DA: dorsal aorta (arrowhead). VV: vitelline vein (arrowhead). Arrow: single migrating Erg+ Sox17+ ECs. Scale: 100 μm. B) Whole-mount Erg and Sox17 immunostaining of E8.5 mouse embryo. C) Percentage of Erg+ ECs in the E9.5 dorsal aorta (DA) or cardinal vein (CV) that express Sox17. Error bars: SD. D) Left: whole-mount Erg immunostaining of E9.5 mouse embryo. Right: E9.5 mouse embryo sectioned and immunostained for Sox17 and Erg proteins, alongside *in situ* staining for *Aplnr* mRNA. Arrow: cardinal vein. Scale: 200 μm. E) *sox7* and *aplnrb* staining of 27-hour post fertilization zebrafish embryos by HCR3 *in situ* hybridization. Lateral view of the mid-trunk region. Scale: 50 μm. F) 4OHT was administered *in utero* to E9.5 *Sox17-Cre*; *Aplnr-DreER*; *RLTG* mouse embryos^66,70,71^, which were then isolated at E12.5, cryosectioned, and immunostained for Erg and GFP proteins. Arrows: GFP+ vein ECs. Scale: 100 μm. Error bars: SD.

Sox17 and *Aplnr* were co-expressed in at least two vascular beds of the E9.5-E10 mouse embryo: the vascular plexus and early cardinal vein. First, the vascular plexus represents a plastic progenitor of both arteries and veins^10^, and we found that the head and limb plexus expressed both Sox17 and *Aplnr* (**Fig. S8D**). Second, the common cardinal vein—the earliest intraembryonic vein^37^—also co-expressed Sox17 and *Aplnr*. This was independently confirmed by two different means to detect *Aplnr* expression: *in situ* hybridization (**Fig. 4D**) and Aplnr-CreER activity^20^ (**Fig. S8D-E**). While the early cardinal vein anatomically appears as a vein, from a molecular perspective it transiently expresses Sox17 and *Aplnr*; it thus mirrors primed ECs *in vitro*, prior to downregulating “arterial” marker Sox17^16,17^. Suggestive of evolutionary conservation of this primed state, we found that *sox7* (a zebrafish *sox17* homolog^68^) was co-expressed alongside *aplnrb* in the zebrafish secondary vascular field (**Fig. 4E**), a plastic progenitor field capable of generating both arteries and veins^69^.

Finally, we employed intersectional lineage tracing to test whether *Sox17*+ *Aplnr*+ ECs contribute to both artery and vein ECs. *Sox17-Cre*^66^ and *Aplnr-DreER*^70^ driver mice were crossed with an intersectional reporter mouse^71^. In this approach, *GFP* expression denotes both Cre- and Dre-mediated recombination^71^, reflecting previous expression of *Sox17* and *Aplnr*. Tamoxifen injection at E8.5 led to GFP labeling of the E12.5 common cardinal vein, supporting the hypothesis that primed ECs form vein ECs *in vivo*; artery ECs were also labeled at lower frequency (**Fig. 4H**). Taken together, *Sox17*+ *Aplnr*+ ECs contribute to both veins and arteries in mouse embryos, and cells with molecular features of primed ECs may be evolutionarily conserved in zebrafish embryos.

### SOXF transcription factors are required for both artery and vein differentiation

Our discovery that SOX17 marks primed ECs *in vivo* and *in vitro* was surprising, as the prevailing view is that the SOXF transcription factor (TF) family, including SOX7, SOX17, and SOX18, imparts arterial identity^13-19^. Primed ECs express all three *SOXF* TFs (**Fig. 5A**). Beyond artery identity, are SOXF TFs also required for human vein EC differentiation? To answer this question, we first engineered hPSCs to express an enhanced CRISPR interference (CRISPRi) effector (dCas9-ZIM3 KRAB)^72^. hPSCs were transduced with sgRNAs targeting *SOX7*, *SOX17,* and/or *SOX18*, and then subject to EC differentiation (**Fig. 5B**). Individual CRISPRi knockdown of either *SOX7*, *SOX17*, or *SOX18* modestly affected arteriovenous identity (**Fig. S9A-B**), consistent with genetic redundancy between SOXF TFs^7,68,73^. We thus simultaneously knocked down all three *SOXF* TFs using CRISPRi with high efficiency (**Fig. S9C-D**). As expected^13-19^, triple *SOXF* knockdown abrogated artery differentiation, with near-complete loss of arterial markers *CXCR4* and *DLL4* (**Fig. 5C-D**, **Fig. S9E-F**, **Table S5**).

**Figure 5:**
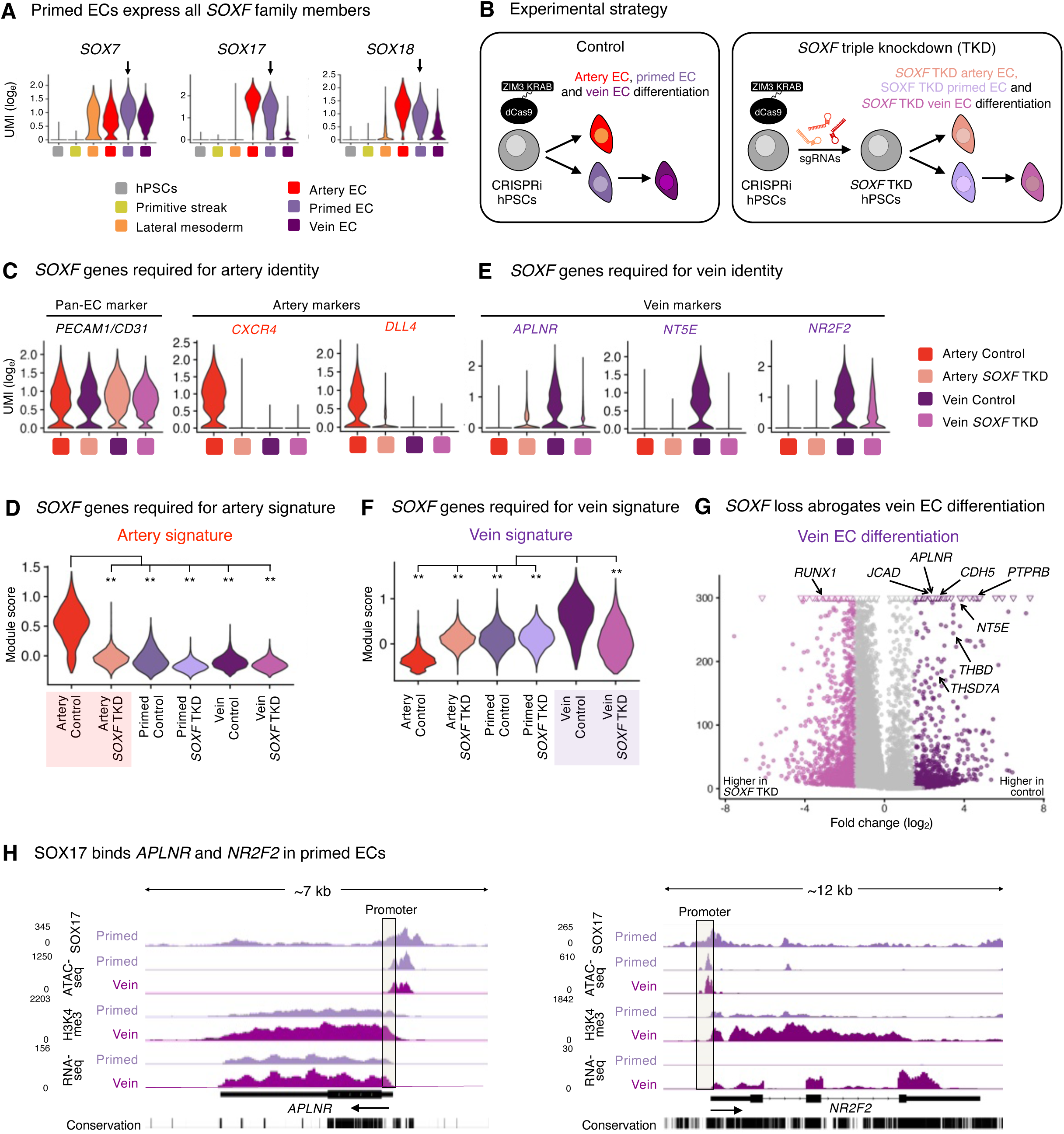
SOXF transcription factors are required for human vein EC specification *in vitro*. A) scRNA-seq of day-2 lateral mesoderm, CD144+ FACS-purified day-3 artery ECs, CD144+ FACS-purified day-3 primed ECs, and CD144+ FACS-purified day-4 vein ECs generated from H1 hPSCs. Arrows indicate that primed ECs express *SOX7*, *SOX17*, and *SOX18*. B) H1 CRISPRi-expressing hPSCs were transduced with sgRNAs targeting *SOX7*, *SOX17*, and *SOX18* (*SOXF* triple knockdown [TKD]), and then subsequently differentiated into artery and vein ECs. C) scRNA-seq of day-3 artery ECs and day-4 vein ECs generated from H1 control vs. *SOXF* TKD hPSCs. Mesenchymal cells were computationally excluded. D) scRNA-seq of day-3 artery ECs, day-3 primed ECs, and day-4 vein ECs generated from H1 control vs. *SOXF* TKD hPSCs. Mesenchymal cells were computationally excluded. A transcriptional module score^93^ computed from a panel of *in vivo*-defined arterial marker genes *in vivo*^24^ is shown. Statistics: Wilcoxon rank sum test. **P<0.01. E) scRNA-seq of day-3 artery ECs and day-4 vein ECs generated from H1 control vs. *SOXF* TKD hPSCs. Mesenchymal cells were computationally excluded. F) scRNA-seq of day-3 artery ECs, day-3 primed ECs, and day-4 vein ECs generated from H1 control vs. *SOXF* TKD hPSCs. Mesenchymal cells were computationally excluded. A transcriptional module score^93^ computed from a panel of *in vivo*-defined venous marker genes *in vivo*^24^ is shown. Statistics: Wilcoxon rank sum test. **P<0.01. G) scRNA-seq of day-3 artery ECs and day-4 vein ECs generated from H1 control vs. *SOXF* TKD hPSCs. Mesenchymal cells were computationally excluded. Differentially expressed genes between control vs. *SOXF* TKD ECs are colored. H) OmniATAC-seq, CUT&RUN, and bulk RNA-seq of H1 hPSC-derived day-3 primed ECs and day-4 vein ECs.

Strikingly, triple *SOXF* knockdown also strongly impaired vein differentiation: venous markers *NR2F2*^22^, *APLNR*^20,21^ and *NT5E/CD73*^51-53^ were reduced (**Fig. 5E**, **Fig. S9E**, **Table S6**). Beyond individual markers, triple *SOXF* knockdown significantly impaired the venous transcriptional signature more broadly, as computed from known venous marker panels^24^ (**Fig. 5F-G**), and downregulated gene sets associated with blood vessel development, migration, and junction formation (**Fig. S9G**). Upon *SOXF* loss, cells retained pan-EC marker *PECAM1* (**Fig. 5C**, **Fig. S9C**), but selectively downregulated *VE-CADHERIN* (**Fig. S9E,H**), thus mirroring zebrafish results^74^.

To investigate why SOXF TFs are required for venous differentiation, we mapped the genome-wide binding of SOX17 in primed ECs using CUT&RUN^55^. SOX17 directly bound the *APLNR* and *NR2F2* promoter elements in primed ECs, prior to venous differentiation and predating *NR2F2* transcription (**Fig. 5H**, **Fig. S9H**). Taken together, SOXF TFs directly bind to, and are required to activate, venous genes during human vein EC differentiation. This builds on the roles of SOXF TFs in mouse and zebrafish vein development^74-76^. We show that while SOXF is required for the specification of both artery and vein ECs, it regulates different genes in each: it is necessary for arterial gene expression in artery ECs, but is required for venous gene expression in vein ECs.

### Accessible chromatin landscapes link VEGF/ERK inhibition to venous identity

We reasoned that TF motifs enriched in either artery- or vein-specific genomic regulatory elements might illuminate regulators of arteriovenous identity in an unbiased way. To this end, we mapped accessible chromatin in FACS-purified hPSC-derived artery and vein ECs using OmniATACseq^49^ (**Fig. 6A**, **Table S7**). AP1 and ETS TFs are transcriptional effectors of ERK signaling^77^, and remarkably their motifs comprised the top 15 TF motifs enriched in artery-specific regulatory elements, and were correspondingly depleted in vein-specific regulatory elements (P<10^-1080^ and P<10^-958^, respectively; **Fig. 6A**, **Table S8-S9**). At the mRNA level, AP1 (*JUNB*) and ETS (*ETV1*, *ETV4*, and *ETV5*) family TFs were upregulated in artery, relative to vein, ECs (**Fig. 6B**). As a positive control, motifs of known arterial (e.g., SOXF) and venous (e.g., NR2F, MAF) TF families were also reciprocally enriched in artery- or vein-specific regulatory elements (**Fig. 6A-B**, **Table S8-S9**). In summary, the accessible chromatin landscape of artery ECs is dominated by ERK transcriptional effectors, whereas vein EC accessible chromatin is depleted of such ERK effectors. The accessible chromatin landscapes of artery and vein ECs reveal how extracellular signals shape chromatin, and link VEGF/ERK inhibition to venous identity.

**Figure 6:**
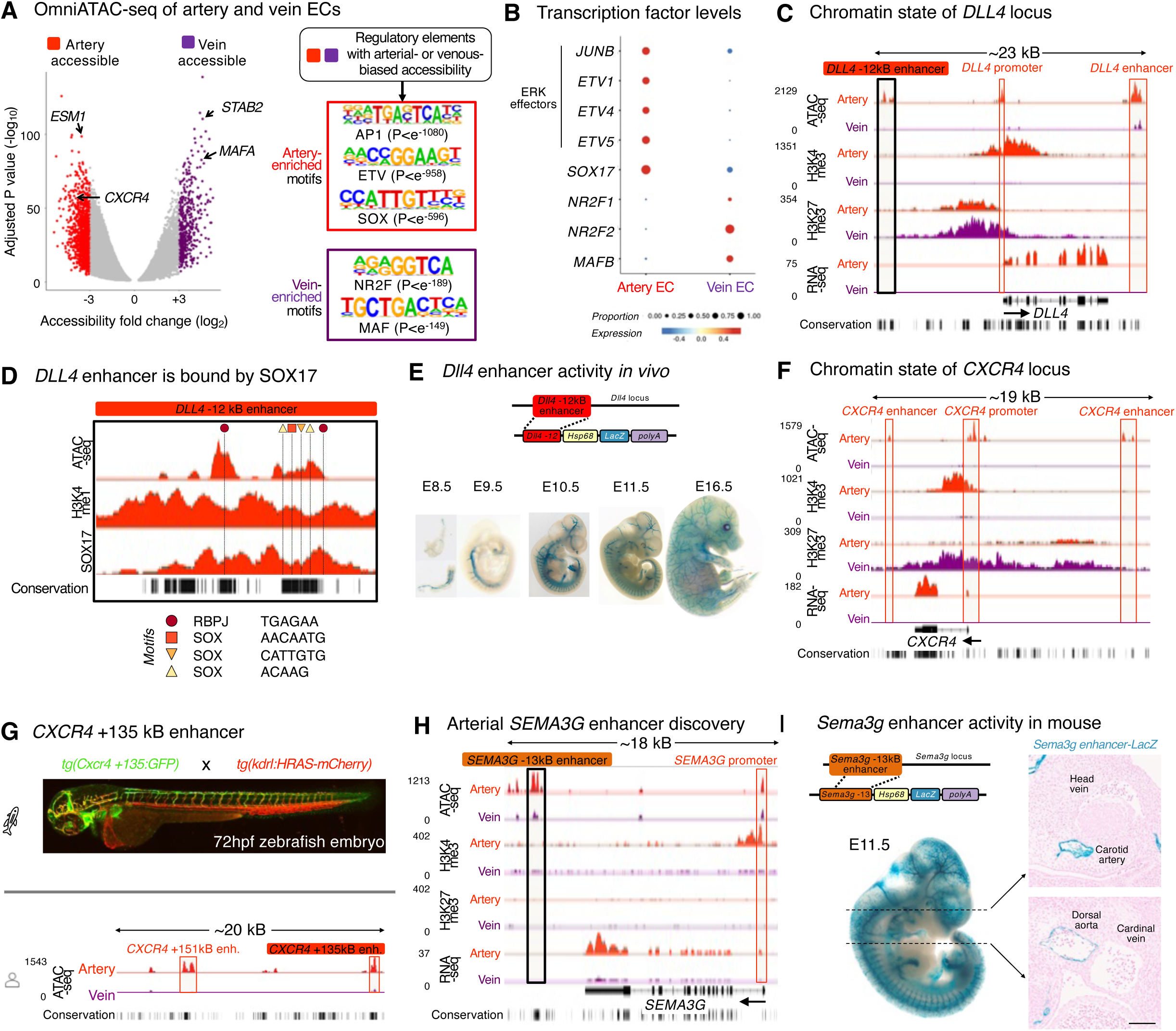
Chromatin analysis links VEGF/ERK inhibition to vein EC specification. A) OmniATAC-seq of H1 hPSC-derived day-3 artery ECs and day-4 vein ECs. Left: genetic loci preferentially accessible in either artery or vein ECs are colored. Right: transcription factor motifs enriched in artery- or vein-accessible chromatin elements. B) Expression of AP1, ETV, SOX, NR2F, and MAF family transcription factors at the mRNA level, as shown by scRNAseq of FACS-purified CD144+ DLL4+ CD73lo/- day-3 artery ECs and CD144+ DLL4- CD73hi day-4 vein ECs generated from H1 hPSCs. C) OmniATAC-seq, CUT&RUN, and bulk RNA-seq of H1 hPSC-derived day-3 artery ECs and day-4 vein ECs. D) OmniATAC-seq and CUT&RUN analysis of the *DLL4 -*12kb enhancer^78^ in H1 hPSC-derived day-3 artery ECs, with transcription factor motifs labeled. E) LacZ staining of E8.5-E16.5 mouse embryos bearing a *Dll4* -12 kB enhancer transgene driving *LacZ* expression^78^. F) OmniATAC-seq, CUT&RUN, and bulk RNA-seq of H1 hPSC-derived day-3 artery ECs and day-4 vein ECs. G) *Top*: image of a 3 day-post-fertilization (dpf) zebrafish bearing a *Cxcr4* +135 kB enhancer driving *GFP* expression, together with a *kdrl:HRAS-mCherry* transgene to label ECs^79^. *Bottom*: ATAC-seq of H1 hPSC-derived day-3 artery ECs and day-4 vein ECs, highlighting the *CXCR4* +135 kB enhancer whose ortholog was tested in the zebrafish transgenic assay^79^. H) OmniATAC-seq, CUT&RUN, and bulk RNA-seq of H1 hPSC-derived day-3 artery ECs and day-4 vein ECs. I) LacZ staining of E11.5 mouse embryo bearing a *Sema3g* -13 kB enhancer transgene driving *LacZ* expression (VISTA Enhancer Browser ID hs2179)^82,83^. Scale: 100 μm.

Chromatin profiling illuminated a wealth of regulatory elements with arterial- or venous-specific accessibility (**Fig. 6A**). In addition to known arterial enhancer elements in the *DLL4* and *CXCR4* loci^78,79^ (**Fig. 6C-G**), our chromatin data revealed a novel, arterial-specific enhancer 13 kB upstream of *SEMA3G* gene, which encodes an arterially expressed guidance cue required for vascular pathfinding^80,81^ (**Fig. 6H**). Transgenic reporter assays delineated that this novel *Sema3g* enhancer was active in arteries, but not veins, within mouse^82,83^ and zebrafish embryos, attesting to its arterial specificity (**Fig. 6I, Fig. S10A**). Taken together, hPSC-derived artery and vein ECs are distinguished by a wealth of arteriovenous regulatory elements at the chromatin level. Our genomic resource provides a means to identify additional such elements (**Table S7**, **Fig. S10B-C**), as exemplified by the arterial *SEMA3G* enhancer.

## DISCUSSION

Curiously, despite incisive insights into mechanisms of artery EC development^3-10,23,27-36^, longstanding mysteries have surrounded vein EC development^6,8^. What are the early embryonic progenitors that build veins? Why do veins often emerge after arteries *in vivo*^24,37-42^? What are the upstream extracellular signals that specify venous identity? Here we discover primed ECs that co-express the “arterial” marker *Sox17* and “venous” marker *Aplnr*, and can form vein ECs *in vivo* and *in vitro*. Consistent with widespread arteriovenous plasticity within the vasculature^10^, primed ECs are not lineage-committed, as they also competent to generate artery ECs *in vivo* and *in vitro*. Collectively, the concept of primed ECs unifies several observations concerning endothelial development, including why the first intraembryonic veins generally emerge after arteries and why the earliest veins transiently “mis-express” certain arterial markers^24,37-42^. We further discover that VEGF/ERK inhibition is crucial for primed ECs to differentiate into vein ECs.

### VEGF/ERK activation, followed by inhibition, drives vein EC differentiation

We present evidence for a two-step model for vein development *in vitro*: VEGF/ERK *activation* is initially required to endow endothelial identity during the conversion of lateral mesoderm into primed ECs, but 24 hours later, VEGF/ERK *inhibition* is subsequently required for vein EC specification. This reconciles the paradox of why VEGF/ERK both promotes^45^ and represses^27-29^ vein ECs *in vivo*: the same signal executes both roles, but at different times. The idea that VEGF/ERK inhibition is required for vein specification may be surprising, as VEGF activation is often construed as being broadly necessary for EC identity^4,27,45^. However, timing is paramount: while VEGF induces ECs, we suggest that once a cell has acquired EC identity, it now re-interprets VEGF/ERK inhibition to signify that it should adopt vein fate. Indeed, in other developmental venues, the same extracellular signal can specify different cellular identities at distinct times^84^. We propose that VEGF is a double-edged sword in venous differentiation: it induces ECs but subsequently represses venous identity. The temporally dynamic effects of VEGF likely contribute to past challenges in generating vein ECs *in vitro*. Indeed, VEGF is ubiquitous in EC differentiation and culture media, but our findings suggest that it may inadvertently suppress venous identity in certain contexts.

Our results may also pertain to the resistance of tumor vasculature against VEGF inhibitor therapies designed to block tumor angiogenesis^46,47^. If VEGF/ERK inhibition instructs ECs to adopt venous identity in certain contexts, this may represent a possible mechanism contributing to VEGF inhibitor resistance.

Why do major intraembryonic arteries precede veins *in vivo*? The answer may lie in how VEGF induces both endothelial and arterial identity^27-29,45^. Perhaps cells can contemporaneously acquire endothelial and arterial identity^37^ under the command of VEGF, but vein development must be parsed into two distinct steps wherein endothelial identity is first instated, followed by VEGF/ERK inhibition to specify veins. At the chromatin level, future venous genes—such as *NR2F2*—are in a poised state in primed ECs, but are rapidly upregulated upon VEGF/ERK inhibition. This fills a vital missing link in our knowledge of the upstream extracellular signals that ignite *NR2F2* expression^6,8^.

In other developmental contexts, the rich detail afforded by scRNAseq suggests that differentiation paths are continua^85,86^, piquing the philosophical question of where one cell state (e.g., primed) ends and the other (e.g., vein) begins. Nevertheless, primed ECs represent an intermediate step in arteriovenous differentiation, as they differ from vein ECs in multiple ways. First, they are functionally induced by opposing signals: VEGF/ERK activation vs. inhibition, respectively. Second, primed ECs do not express later-stage venous markers such as *NR2F2*, but harbor bivalent chromatin at future venous genes, implying developmental poising. Third, primed ECs lack venous cell-surface markers CD73 and CD317.

### SOXF transcription factors are required for vein and artery EC specification

While the early cardinal vein anatomically appears as a vein, we find that it molecularly co-expresses Sox17 and *Aplnr*. The early cardinal vein may thus transiently resemble primed ECs, prior to downregulating Sox17 at later stages^16,17^. Indeed, most if not all ECs temporarily express Sox17 in the mouse embryo, as shown by immunostaining and *Sox17-Cre* lineage tracing. Consistent with how the early vasculature broadly expresses Sox17 *in vivo*, simultaneous CRISPRi knockdown of all *SOXF* family members (*SOX7*, *SOX17*, and *SOX18*) compromised both human artery and vein EC differentiation *in vitro*.

Our findings support longstanding observations that SOXF transcription factors are required for arterial specification^13-19^, but additionally suggest important roles in venous identity. Further evidence supports our conclusions. First, both arteries and veins are morphologically lost in the *Sox7*-/- *Sox17*-/- *Sox18*-/- mouse retinal vasculature^76^. Second, double *sox7* and *sox18* knockdown abrogates vein development in zebrafish^74,75^. Third, SOXF motifs reside in not just arterial enhancers, but also in pan-endothelial enhancers, implying that SOXF has broader roles in endothelial identity beyond arterialization^79^. Taken together, SOXF transcription factors are expressed in primed ECs, and are crucial for human vein EC differentiation.

While SOXF is necessary for the specification of artery and vein ECs, it activates different genes in each. Our CRISPRi studies showed that SOXF upregulates arterial genes in artery ECs, whereas it drives expression of certain venous genes (e.g., *NR2F2* and *APLNR*) in vein ECs. Indeed, TFs execute cell-type-specific roles^87^. Could differing SOXF levels contribute to these outcomes? Artery ECs express higher levels of SOX17 than primed ECs (**Fig. 1D**). Perhaps higher vs. lower SOXF levels respectively bias ECs to arterial vs. venous identities, but wholesale SOXF depletion—as we achieved by CRISPRi—compromises both. SOXF TFs may also interact with differing cofactors in either artery vs. vein ECs to direct distinct transcriptional programs.

Taken together, we propose a two-step model for vein differentiation, wherein VEGF activation initially differentiates mesoderm into primed ECs, and subsequently, VEGF/ERK inhibition specifies vein ECs. This provides a new perspective on vein development and has practical ramifications, as hPSC differentiation into primed ECs represents a critical step to generate vein ECs.

## LIMITATIONS OF THE STUDY

First, VEGF/ERK is crucial for incipient EC specification, but how does it subsequently become repressed to permit vein EC specification *in vivo*? When and where VEGF ligands are expressed is the most parsimonious explanation for this temporally dynamic VEGF/ERK signaling switch. Initially, VEGF differentiates lateral mesoderm into ECs *in vivo*^27,45,88^. Subsequently, future cardinal vein ECs become separated from the VEGF source by several hundred microns, as shown in *Xenopus*^89^, likely explaining why VEGF/ERK signaling becomes attenuated to drive vein specification.

Second, what are the molecular mechanisms through which VEGF/ERK inhibition endows venous identity? In other cell-types, ERK2 transcriptionally pauses RNA polymerase II on future developmental genes^90^. It will be interesting to test if RNA polymerase II elongation is unleashed upon VEGF/ERK inhibition in primed ECs, driving *NR2F2* upregulation.

Third, our intersectional lineage tracing revealed that *Sox17*+ *Aplnr*+ ECs contribute to vein ECs, but not every vein EC was labeled. This likely reflecting incomplete recombination efficiency inherent to intersectional lineage tracing, or alternative sources of vein ECs. An open question remains whether *all* vein ECs emerge via primed EC intermediates, or whether alternative differentiation paths can also culminate in venous identity. This might be addressable in the future through unbiased CRISPR barcoding^91,92^.

## ACKNOWLEDGEMENTS

We thank Valerie Park, Liying Ou, Aaron McCarty, Catherine Carswell-Crumpton, the Stanford Institute for Stem Cell Biology & Regenerative Medicine, and the Stanford Stem Cell FACS Core Facility for infrastructure support. We thank Sophie Astrof for sharing *Aplnr-DreER* mice, and Ryan Corces, Arjun Rajan, Zhainib Amir-Ugokwe, Siva Vijayakumar, Len Pennacchio, Axel Visel, Stacey Lee, Avishek Ganguly, Yan Yang, Yimiao Qu, Alicia Wong, and Julie Chen for assistance. Genomics analyses were supported by the Stanford Diabetes Genomics Analysis Core Facility (NIH P30DK116074). This work was supported by NIH Director’s Early Independence Award DP5OD024558 (K.M.L.), NIH R01HL128503 (K.R.-H.), NIH R01HL153005 (S.S.), NIH R01HL168097 (C.M.), Breakthrough T1D Northern California Center of Excellence (K.M.L.), Additional Ventures Expansion Award (K.M.L., L.T.A.), Siebel Stem Cell Institute (L.T.A.), Ludwig Cancer Research (K.M.L., S.D.V.), Stanford Maternal and Child Health Research Institute Chambers Family Foundation Innovation Grant (K.M.L., L.T.A., K.R.-H.), Advanced Research Projects Agency for Health (ARPA-H) under award number AY1AX000002 (M.A.S.-S., K.M.L.), Helmholtz Society (H.L.), German Research Foundation (H.L.), European Research Council (H.L.), British Heart Foundation FS/SBSRF/22/31037 and RE/18/3/34214 (S.D.V.), Amaranth Foundation (K.M.L.), and Anonymous and Gilbert families (K.M.L.). S.L.Z. was supported by NSF and Stanford Graduate Fellowships. A.M. was supported by the Stanford Graduate Fellowship. R.M.M. was supported by NIH T32GM119995. C.M. is the The Helen and Arthur E. Johnson Chair For The Cardiac Research Director. L.T.A. is an Additional Ventures Catalyst to Independence Fellow and a Bladder Cancer Advocacy Network Career Development Awardee. K.R.-H. is a Howard Hughes Medical Institute Investigator. K.M.L. is a Human Frontier Science Program Young Investigator (RGY0069/2019), Packard Foundation Fellow, Pew Scholar, Baxter Foundation Faculty Scholar, and The Anthony DiGenova Endowed Faculty Scholar. Content is solely the responsibility of the authors and does not necessarily represent the official views of the NIH, ARPA-H, or other funding organizations.

## AUTHOR CONTRIBUTIONS

L.T.A., S.L.Z., K.J.L., A.M., Q.Y., C.Q., X.X., and K.M.L. differentiated and characterized hPSCs. S.L.Z., J.W., S.K.J., J.A., D.W.S., and K.R.-H. analyzed mouse embryos. L.T.A., S.L.Z., K.J.L., A.M., X.X., A.D., E.X., W.S., C.M., S.D.V., and K.M.L. performed genomics analyses. R.S., K.S., R.M.M., B.J.L., J.S.W., H.L., M.A.S.-S., and M.H.P. developed and provided reagents and advice. N.K.R. and S.S. stained zebrafish embryos. D.S., S.N., and S.D.V. analyzed enhancer transgenic embryos. L.T.A., K.R.-H., and K.M.L. supervised the study, with input from S.L.Z., K.J.L., and A.M.

## DECLARATION OF INTERESTS

Stanford University has filed patent applications related to endothelial differentiation. S.L.Z. and R.M.M. are presently employed in the biotechnology industry, but contributed to this work while affiliated with Stanford University.

## SUPPLEMENTAL FIGURE LEGENDS

**Figure S1:**
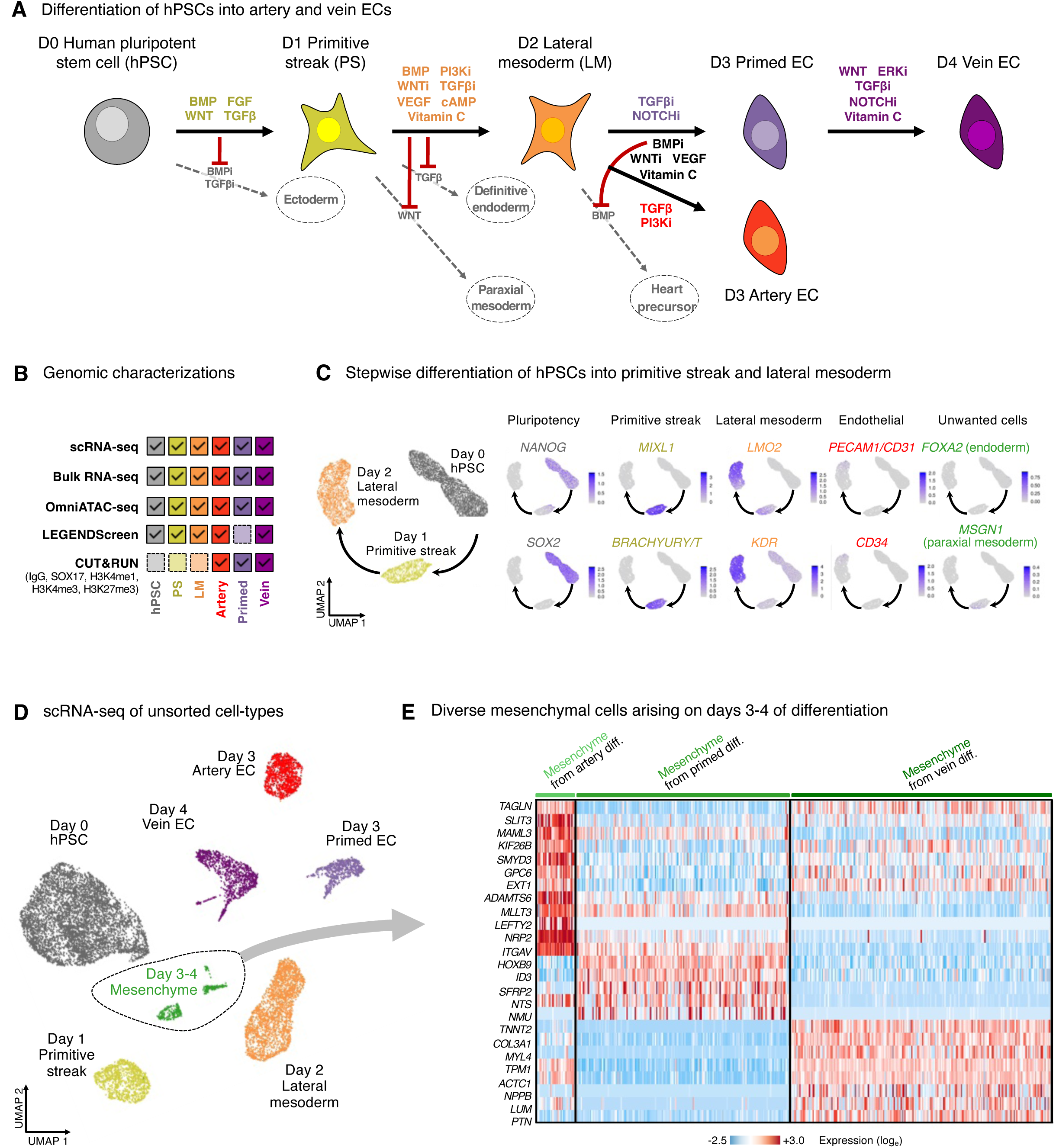
Cellular diversity during early steps of hPSC differentiation. A) Summary of hPSC differentiation approach^48^. i: inhibition. B) Summary of gene expression, chromatin state, and cell-surface marker profiling conducted as part of this study. Dotted lines indicate that profiling was not conducted on a given cell-type. C) Gene expression on days 0, 1, and 2 of H1 hPSC differentiation, as detected by scRNAseq. D) scRNAseq of day-0 H1 hPSCs that were differentiated into day-1 primitive streak, day-2 lateral mesoderm, day-3 artery ECs, day-3 primed ECs, and day-4 vein ECs. None of cell populations shown here were FACS purified and consequently, mesenchymal cells were detected alongside artery, primed, and vein ECs on days 3-4 of differentiation. The same day-0 hPSC, day-1 primitive streak, and day-2 primitive streak scRNAseq datasets were also used in Fig. 1A; however, FACS-purified downstream EC populations are shown in Fig. 1A. E) scRNAseq of H1 hPSC-derived mesenchymal cells that arose alongside day-3 artery ECs, day-3 primed ECs, and day-4 vein ECs. Differentially expressed genes that distinguish these three different types of mesenchyme are shown. Diff: differentiation.

**Figure S2:**
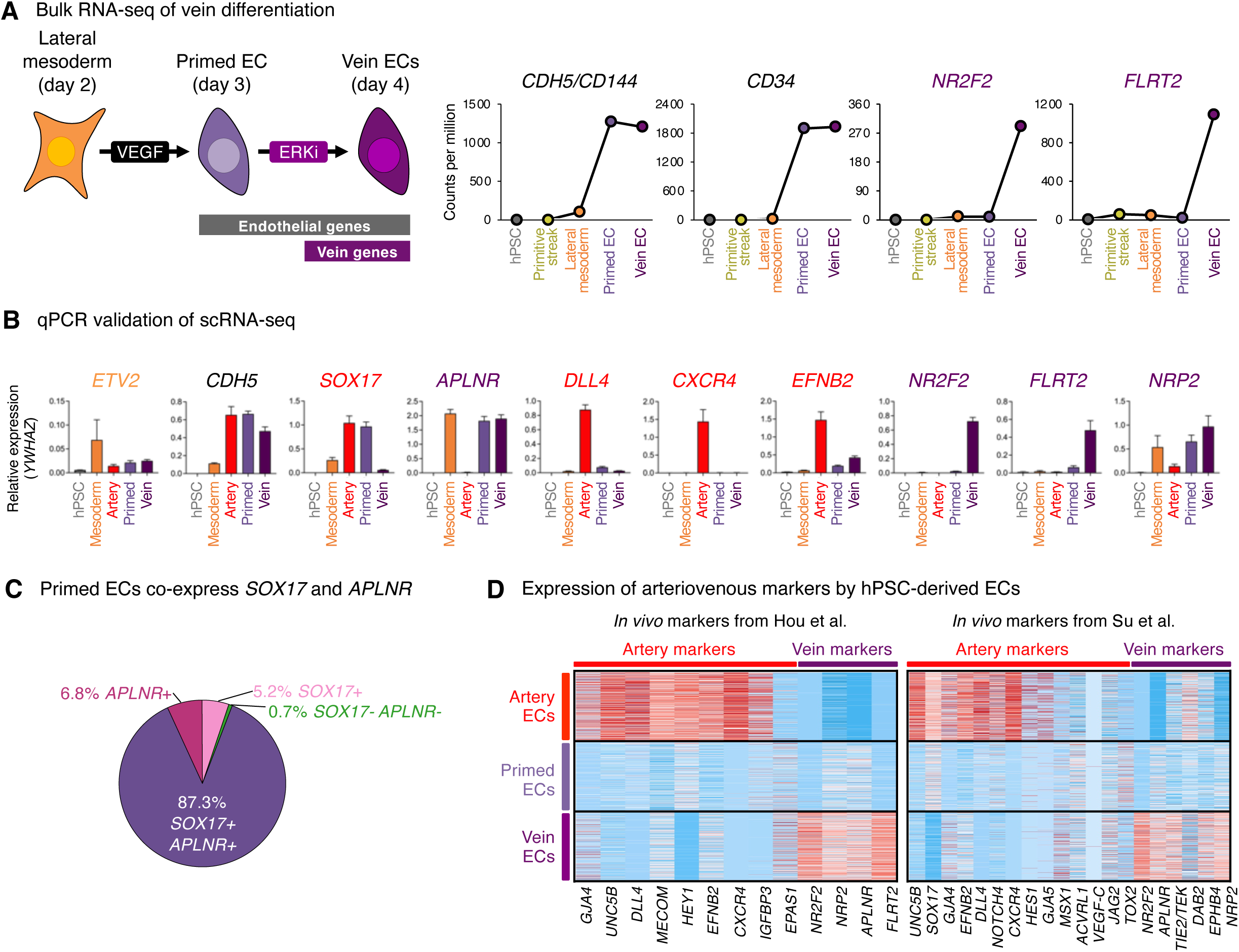
Characterization of primed endothelial cells *in vitro*. A) Bulk RNA-seq of the indicated H1 hPSC-derived cell-types. FACS was used to purify CD144+ primed and vein ECs for RNA-seq. i: inhibition. B) qPCR of the indicated H1 hPSC-derived cell-types. qPCR data normalized to reference gene *YWHAZ* (i.e., *YWHAZ* levels = 1.0). Error bars: standard deviation (SD). C) scRNAseq of H1 hPSC-derived day-3 primed EC populations, showing the percentage of cells that expressed *APLNR* and/or *SOX17* mRNAs. D) scRNAseq of FACS-purified CD144+ DLL4+ CD73lo/- day-3 artery ECs, CD144+ day-3 primed ECs, and CD144+ DLL4- CD73hi day-4 vein ECs generated from H1 hPSCs. In each of these *in vitro* cell-types, expression of *in vivo* arterial and venous markers reported by Hou et al., 2022^24^ and Su et al., 2018^23^ is shown.

**Figure S3:**
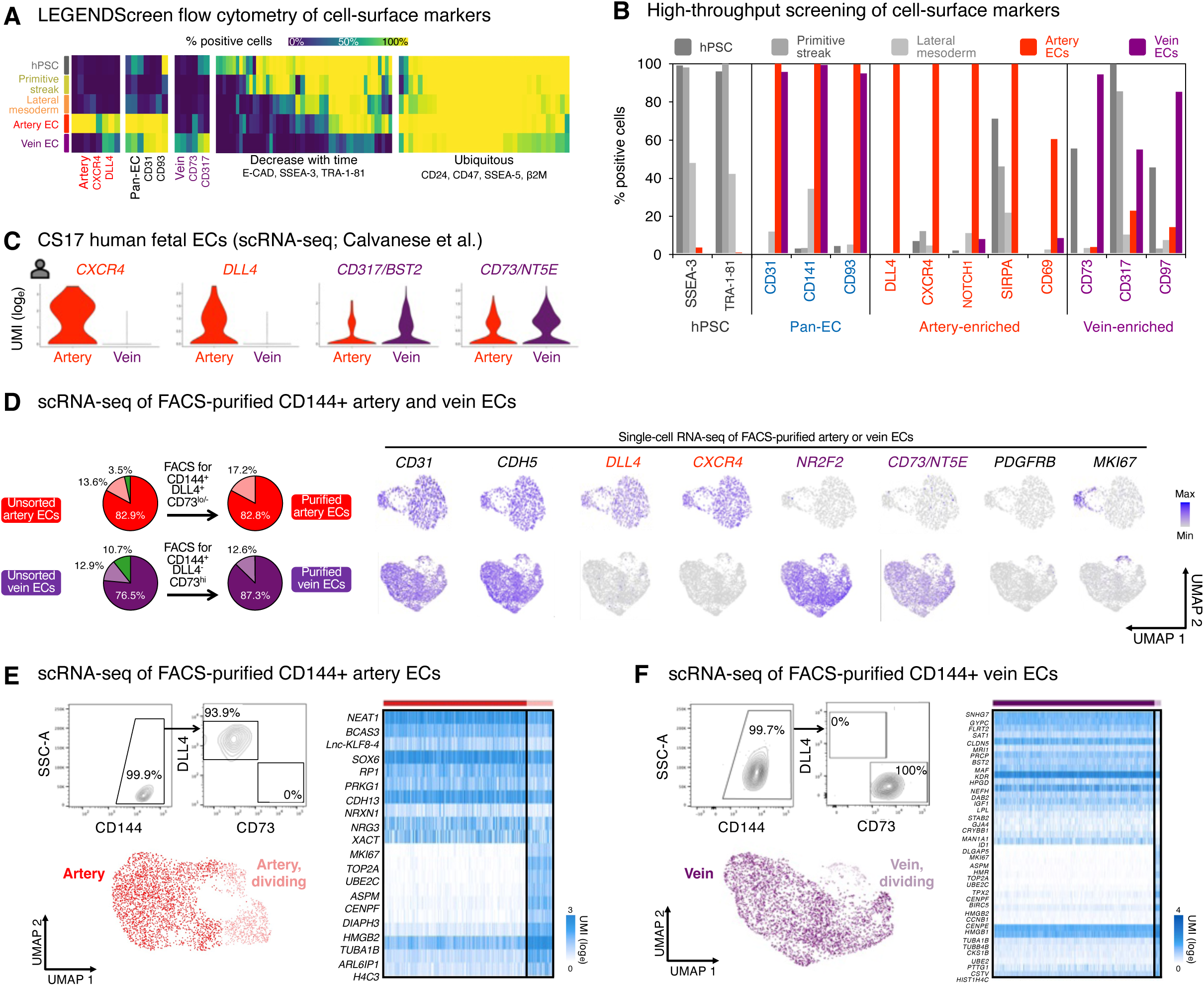
Cell-surface markers of human arteriovenous identity. A) LEGENDScreen high-throughput flow cytometry of cell-surface markers in day-0 hPSCs, day-1 primitive streak, day-2 lateral mesoderm, day-3 artery ECs, and day-4 vein ECs. Day-3 artery and day-4 vein EC populations were pre-gated on the CD144+ EC subset before depicting marker expression. B) LEGENDScreen high-throughput flow cytometry of cell-surface markers in day-0 hPSCs, day-1 primitive streak, day-2 lateral mesoderm, day-3 artery ECs, and day-4 vein ECs. Day-3 artery and day-4 vein EC populations were pre-gated on the CD144+ EC subset before depicting marker expression. C) scRNAseq of human Carnegie Stage 17 (CS17) fetal ECs. scRNAseq data were obtained from Calvanese et al., 2022^94^. D) Left: Population heterogeneity of H1 hPSC-derived day-3 artery ECs and day-4 vein ECs, before and after FACS purification based on the indicated cell-surface marker combinations. Proportions of cell clusters in scRNAseq data are shown. Right: scRNAseq of FACS-purified CD144+ DLL4+ CD73lo/- day-3 artery ECs and CD144+ DLL4- CD73hi day-4 vein ECs generated from H1 hPSCs. E) scRNAseq of FACS-purified CD144+ DLL4+ CD73lo/- day-3 artery ECs generated from H1 hPSCs. Left: FACS isolation strategy. Right: subclustering was performed to assess any potential population heterogeneity, and differentially expressed genes that distinguished cell subsets are shown. F) scRNAseq of FACS-purified CD144+ DLL4- CD73hi day-4 vein ECs generated from H1 hPSCs. Left: FACS isolation strategy. Right: subclustering was performed to assess any potential population heterogeneity, and differentially expressed genes that distinguished cell subsets are shown.

**Figure S4:**
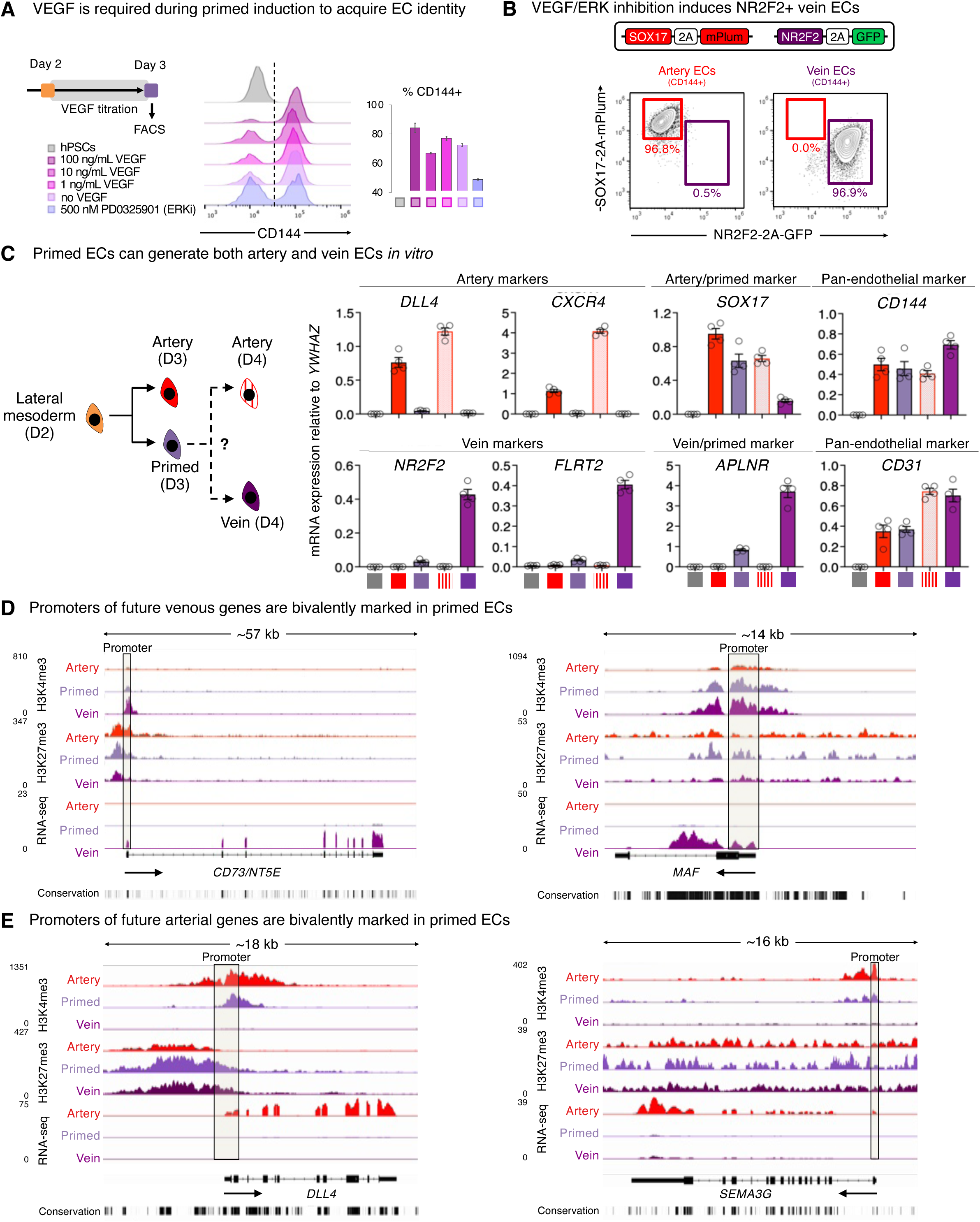
Generation and characterization of primed endothelial cells *in vitro*. A) First, H1 hPSCs were differentiated into day-2 lateral mesoderm^48^. Then, lateral mesoderm was then treated with different doses of VEGF pathway modulators (VEGF [0-100 ng/mL] or ERK inhibitor [ERKi; PD0325901, 500 nM]) alongside other primed EC-inducing signals (TGFϕ3 inhibitor + NOTCH inhibitor + BMP inhibitor + WNT inhibitor + Vitamin C) for 24 hours. Flow cytometry was conducted on day 3 of hPSC differentiation. This revealed that high VEGF for 24 hours is required during primed EC specification to efficiently generate ECs by day 3. B) Flow cytometry of H1 *SOX17-2A-mPlum; NR2F2-2A-GFP* hPSCs differentiated into day-3 artery ECs or day-5 vein ECs. Day-4 vein ECs were maintained in the same vein induction media until day 5. Day 3-5 populations were pre-gated on the CD144+ EC subset before depicting marker expression. C) First, H1 hPSCs were differentiated into day-3 primed ECs. Then, primed ECs were then treated with different doses of VEGF pathway modulators (VEGF [0-100 ng/mL] or ERK inhibitor [ERKi; PD0325901, 500 nM]) alongside other vein EC-inducing signals (TGFϕ3 inhibitor + NOTCH inhibitor + WNT agonist + Vitamin C) for 1-24 hours. qPCR was conducted on day 4 of hPSC differentiation, and expression is normalized to the sample with the highest expression. This revealed that ERK inhibition for 12 hours significantly upregulated venous marker expression. D) CUT&RUN profiling of H3K4me3 and H3K27me3 and bulk RNA-seq H1 hPSC-derived day-3 artery ECs, day-3 primed ECs, and day-4 vein ECs. E) CUT&RUN profiling of H3K4me3 and H3K27me3 and bulk RNA-seq H1 hPSC-derived day-3 artery ECs, day-3 primed ECs, and day-4 vein ECs.

**Figure S5:**
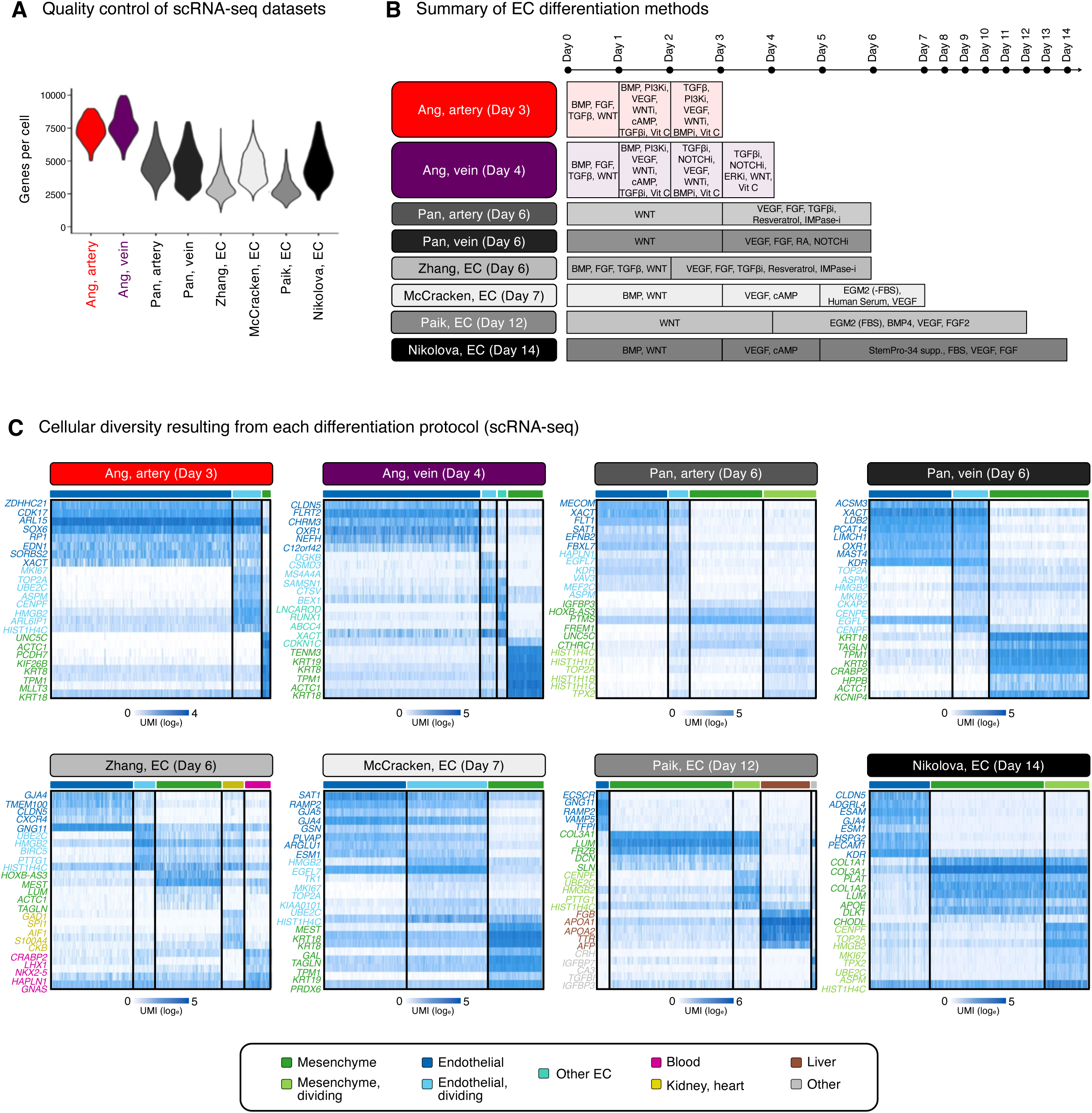
Comparison of methods to differentiate hPSCs into ECs. A) scRNA-seq of hPSCs differentiated into ECs using various protocols, depicting the number of genes detected per single cell as a quality control metric. B) Summary of protocols to differentiate hPSCs into ECs^48,58-62^ used to generate the scRNA-seq datasets described in Fig. 3A. C) scRNA-seq of differentiated hPSC populations described in Fig. 3A. Subclustering was performed to assess population heterogeneity, and differentially expressed genes that distinguished cell subsets are shown. Clusters are annotated and colored as described in Fig. 3A.

**Figure S6:**
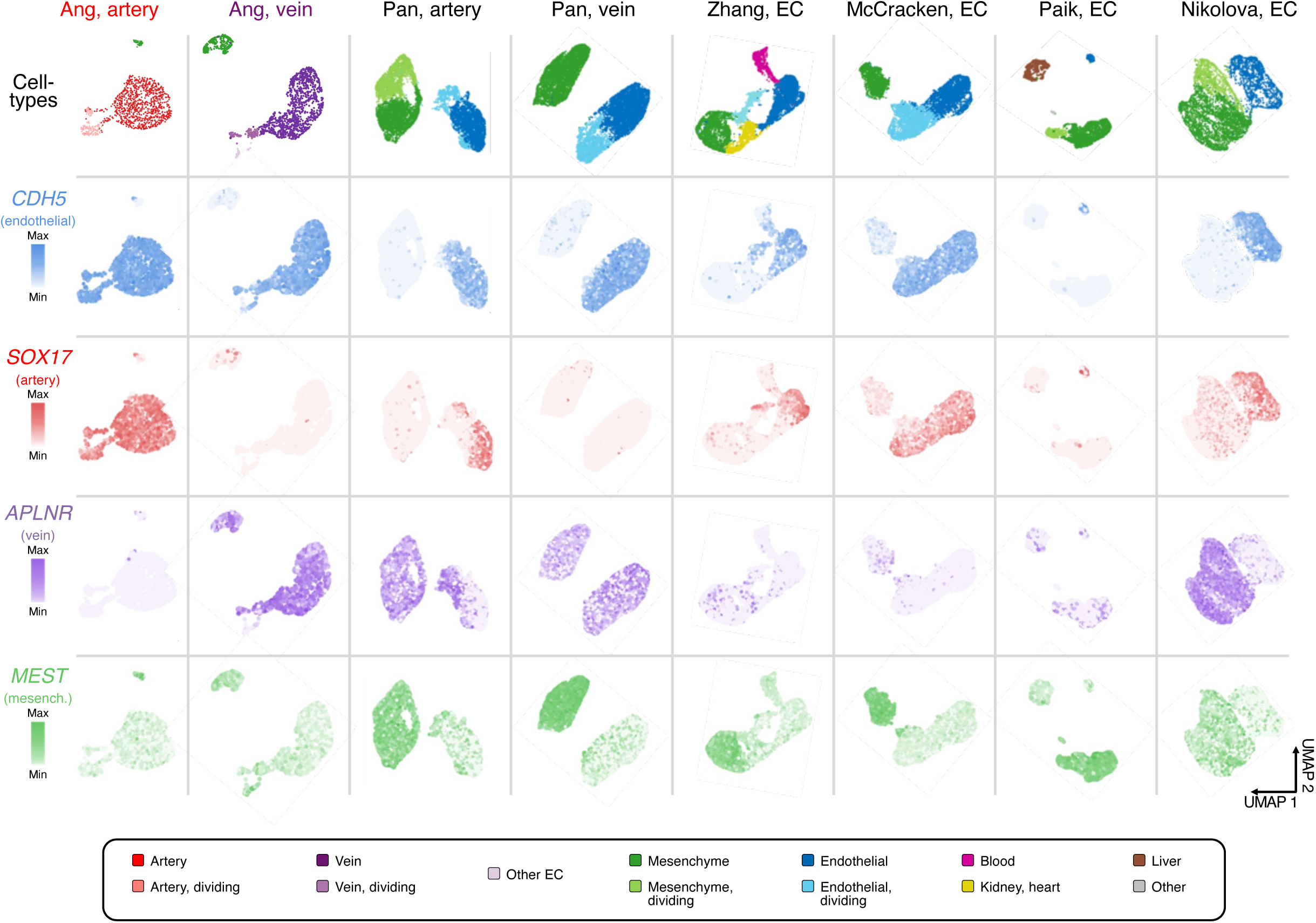
Comparison of methods to differentiate hPSCs into ECs. A) scRNA-seq of differentiated hPSC populations described in Fig. 3A, depicting expression of pan-endothelial, arterial, venous, and mesenchymal marker genes. Clusters are annotated and colored as described in Fig. 3A.

**Figure S7:**
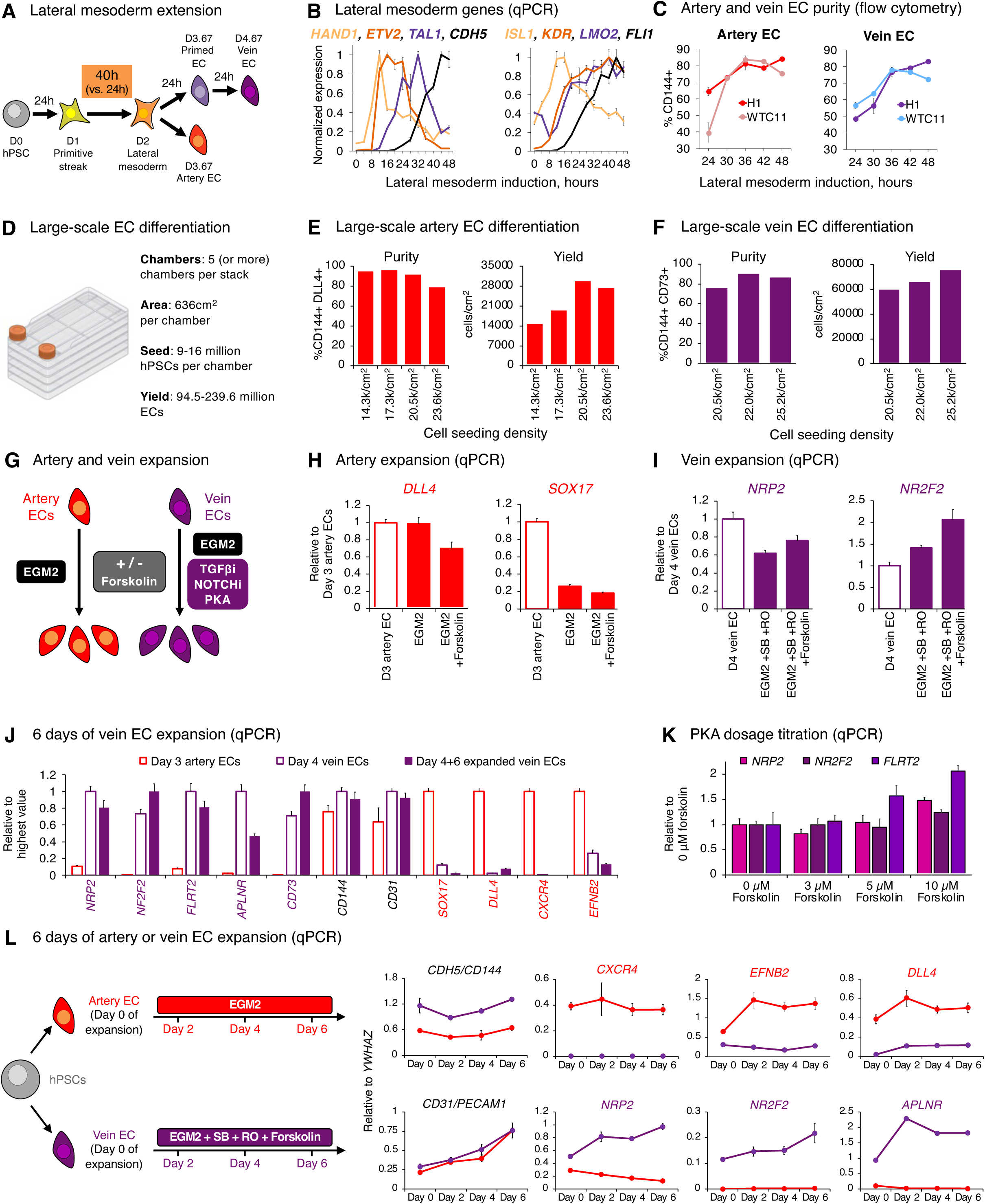
Improved generation and expansion of artery and vein ECs, while preserving arteriovenous identity. A) Schematic of differentiation protocol, which includes prolongation of lateral mesoderm induction from 24 hours (V1 protocol) to 40 hours (V2 protocol). B) qPCR of lateral mesoderm and vascular marker expression in H1 hPSC-derived day-1 primitive streak cells challenged with lateral mesoderm induction media for up to 48 hours. 0 hours corresponds to day-1 primitive streak cells, prior to addition of lateral mesoderm induction media. Cells were lysed every 4 hours and qPCR was performed. For each gene, expression is normalized to the sample with the highest value, which was set to “1.0”. Error bars: SEM. C) Effect of lateral mesoderm induction duration on the subsequent production of CD144+ artery and vein ECs from the H1 and WTC11 hPSC lines, as measured by flow cytometry. Duration of lateral mesoderm induction was varied from 24 hours (V1 protocol) to 48 hours, with all other parameters of the differentiation protocol unchanged. Error bars: SEM. D) Schematic of CellSTACK system for large-scale differentiation of hPSCs into ECs. E) Yield and percentage of CD144+ DLL4+ day-3.67 artery ECs generated in the large-scale CellSTACK differentiation system, depending on initial hPSC seeding density. Yield and differentiation purity were respectively measured by flow cytometry and cell counting. F) Yield and percentage of CD73+ DLL4+ day-4.67 vein ECs generated in the large-scale CellSTACK differentiation system, depending on initial hPSC seeding density. Yield and differentiation purity were respectively measured by flow cytometry and cell counting. G) Strategy for expanding hPSC-derived artery and vein ECs. Artery ECs are cultured in EGM2 medium^48^, and vein ECs are cultured in EGM2 medium + TGFβ inhibitor + NOTCH inhibitor + PKA activator. i: inhibitor. H) qPCR of H1 hPSC-derived artery ECs that were thawed in EGM2 medium^48^ and cultured for 6 days. ROCK inhibitor (Thiazovivin, 2 μM) was added for the first day of thawing to improve cell survival, and was subsequently removed in later days. Forskolin (3 μM) was added as indicated. For each gene, expression is normalized to the sample with the highest value, which was set to “1.0”. Error bars: SEM. I) qPCR of H1 hPSC-derived vein ECs that were thawed in EGM2 medium + SB505124 (2 μM, TGFβ inhibitor) + RO4929097 (1 μM, NOTCH inhibitor)^48^ and cultured for 6 days. ROCK inhibitor (Thiazovivin, 2 μM) was added for the first day of thawing to improve cell survival, and was subsequently removed in later days. Forskolin (3 μM) was added as indicated. For each gene, expression is normalized to the sample with the highest value, which was set to “1.0”. Error bars: SEM. J) qPCR of H1 hPSC-derived vein ECs that were thawed in EGM2 medium + SB505124 (2 μM, TGFβ inhibitor) + RO4929097 (1 μM, NOTCH inhibitor) + Forskolin (10 μM, adenylate cyclase agonist) and cultured for 6 days. ROCK inhibitor (Thiazovivin, 2 μM) was added for the first day of thawing to improve cell survival, and was subsequently removed in later days. As a control, H1 hPSC-derived artery ECs prior to expansion were also analyzed. qPCR data are normalized to the sample with the highest expression, which was set to “1.0”. Error bars: SEM. K) qPCR of H1 hPSC-derived vein ECs that were thawed in EGM2 medium + SB505124 (2 μM, TGFβ inhibitor) + RO4929097 (1 μM, NOTCH inhibitor), in the presence or absence of Forskolin (3-10 μM, adenylate cyclase agonist) and cultured for 6 days. qPCR data are normalized to the sample lacking forskolin, which was set to “1.0”. Error bars: SEM. L) qPCR of pan-endothelial, arterial, and venous marker expression over the course of 6-day expansion of SUN004.2 *CAG-mScarlet* hPSC-derived artery and vein ECs. Artery ECs were expanded in EGM2 medium, whereas vein ECs were expanded in EGM2 medium + SB505124 (2 μM, TGFβ inhibitor) + RO4929097 (1 μM, NOTCH inhibitor) + Forskolin (10 μM, adenylate cyclase agonist). qPCR data normalized to reference gene *YWHAZ* (i.e., *YWHAZ* levels = 1.0). Error bars: SEM.

**Figure S8:**
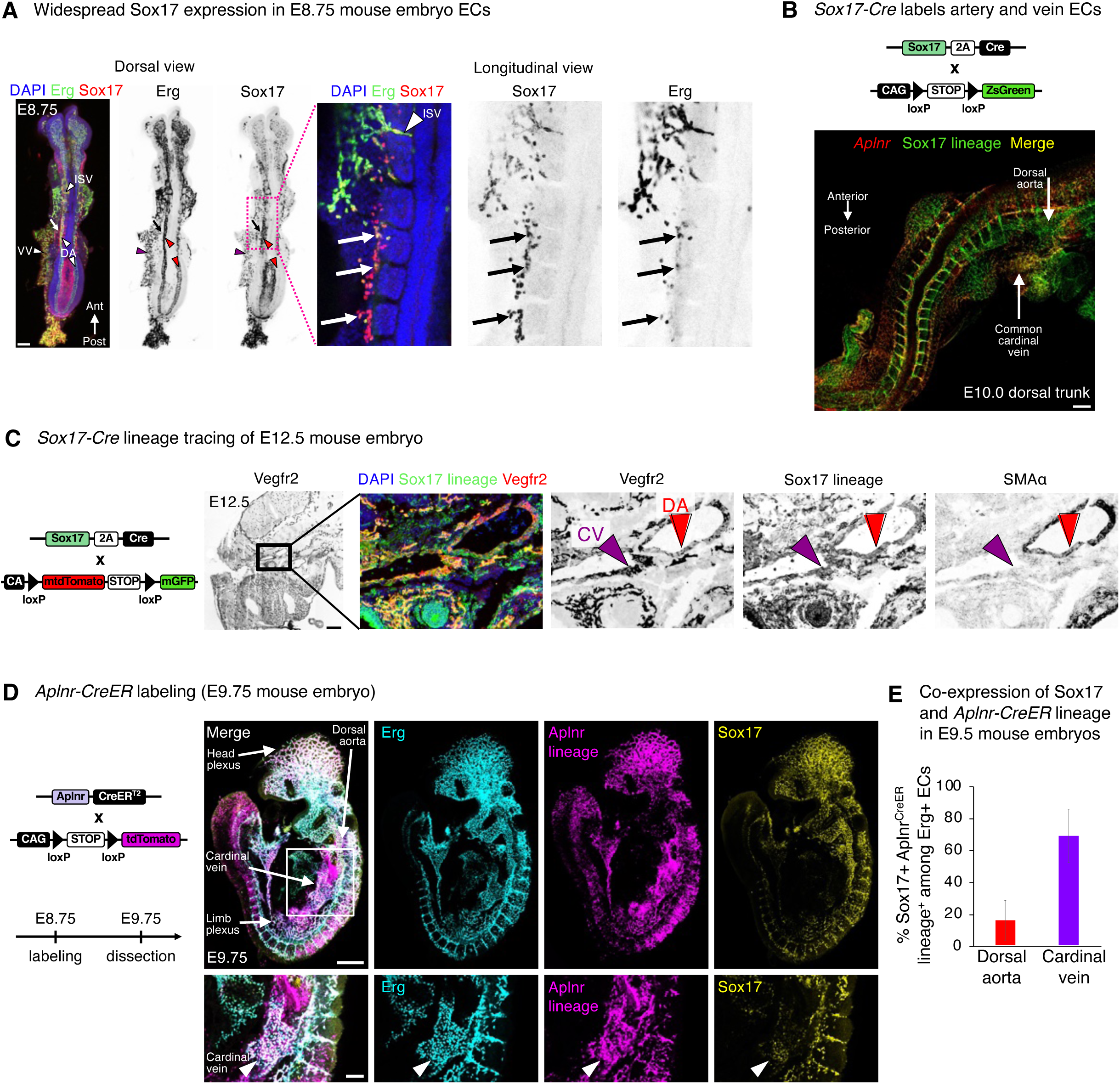
Existence and fate of *Sox17*+ *Aplnr*+ endothelial cells in the mouse embryo. A) Whole-mount Erg and Sox17 immunostaining of E8.75 mouse embryo. DA: dorsal aorta (arrowhead). VV: vitelline vein (arrowhead). ISV: intersomitic vessel (arrowhead). Arrows: solitary Sox17+ Erg+ ECs. Ant: anterior. Post: posterior. Scale: 100 μm. B) Whole-mount *Aplnr* mRNA staining of E10 *Sox17^Cre^; R26^zsGreen^* mouse embryos^66,95^ by HCR3 *in situ* hybridization. Scale: 100 μm. C) E12.5 *Sox17^Cre^; R26^mTmG^* mouse embryos^66,96^ were sectioned and stained for Vegfr2, SMAα, and GFP proteins. Scale: 200 μm. D) 4-hydroxytamoxifen (4OHT) was administered *in utero* to E8.75 *Aplnr-CreER*; *R26-tdTomato* mouse embryos^20,95^, which were then isolated at E9.75 and immunostained for Sox17 and Erg proteins. Bottom row arrows: cardinal vein. Scale: 200 μm (top row), 50 μm (bottom row). E) 4-hydroxytamoxifen (4OHT) was administered *in utero* to E8.75 *Aplnr-CreER*; *R26-tdTomato* mouse embryos^20,95^, which were then isolated at E9.75 and immunostained for Sox17 and Erg proteins. Quantification of Erg+ ECs in the dorsal aorta and cardinal vein that co-expressed tdTomato (indicative of *Aplnr-CreER* activity) and Sox17. Error bars: SD. **P<0.01.

**Figure S9:**
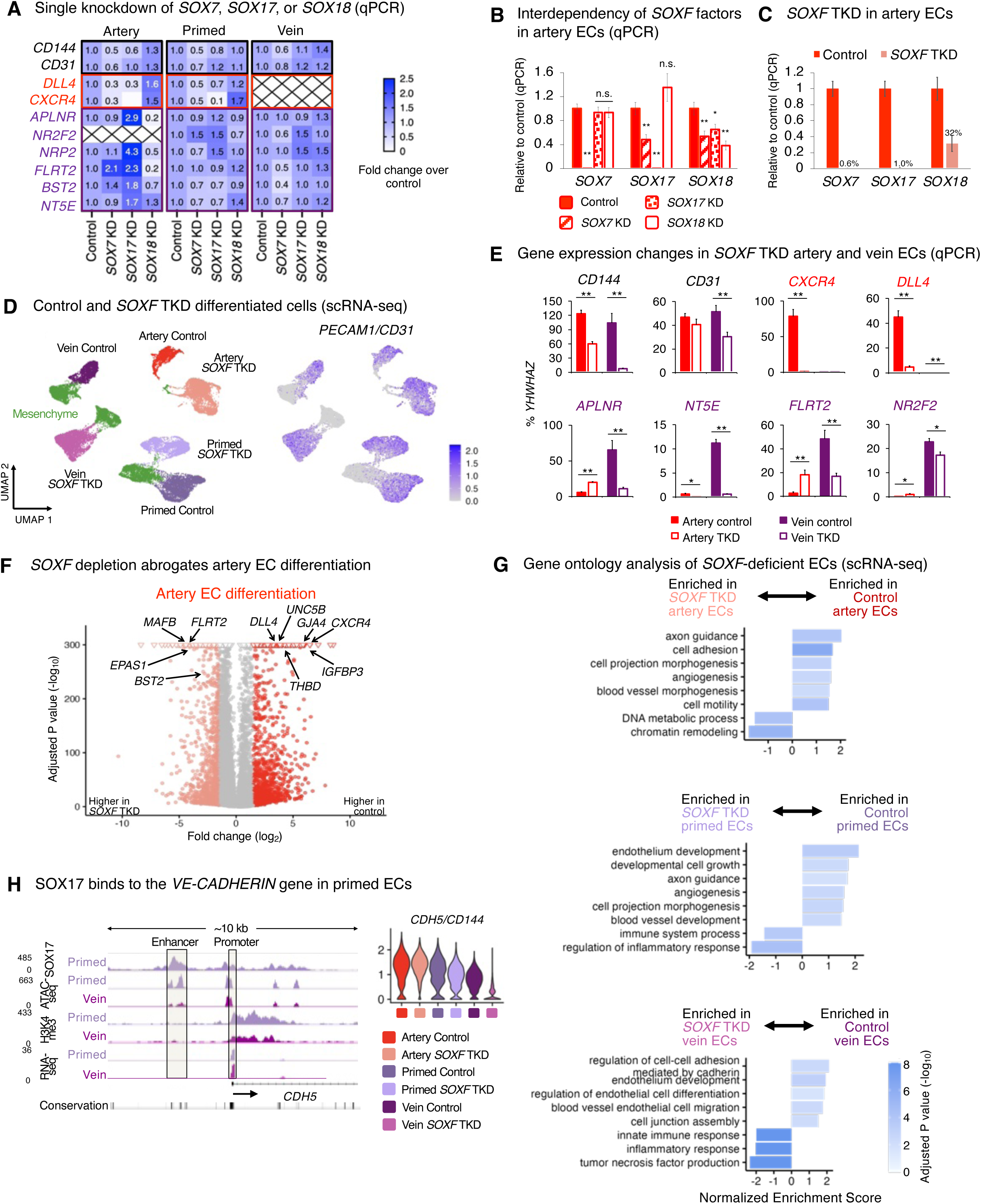
Triple CRISPRi knockdown of *SOXF* transcription factors impairs both human artery and vein EC differentiation *in vitro*. A) qPCR of day-3 artery ECs derived from control and single *SOX7*-, *SOX17*-, or *SOX18*-knockdown H1 CRISPRi hPSC lines. Gene expression normalized to levels in control hPSC-derived artery ECs, which was set as “1.0”. “X” indicates that qPCR data were not shown, because gene expression was under 2% of *YWHAZ* in control samples. Statistics: unpaired t-test. **P<0.01. B) qPCR of day-3 artery ECs derived from control and single *SOX7*-, *SOX17*-, or *SOX18*-knockdown H1 CRISPRi hPSC lines. Gene expression normalized to levels in control hPSC-derived artery ECs, which was set as “1.0”. Statistics: unpaired t-test. Error bars: SEM. n.s.: not significant. *P<0.05. **P<0.01. C) qPCR of day-3 artery ECs generated from H1 control vs. *SOXF* TKD hPSCs. Gene expression normalized to control artery ECs. Percentages indicate remaining gene expression in *SOXF* TKD artery ECs, relative to controls. Statistics: unpaired t-test. Error bars: standard error of the mean (SEM). D) *Left*: scRNA-seq of day-3 artery ECs, day-3 primed ECs and day-4 vein ECs generated from either H1 control or *SOXF* TKD CRISPRi hPSCs. *Right*: *PECAM1*+ ECs and *PECAM1*- mesenchymal cells were detected. The same scRNAseq datasets are shown here as in Fig. 4C, except that mesenchymal cells were computationally excluded in Fig. 4C. E) qPCR of day-3 artery ECs and day-4 vein ECs generated from H1 control vs. *SOXF* TKD CRISPRi hPSCs. Gene expression normalized to reference gene *YWHAZ* (i.e., *YWHAZ* = 100%). Statistics: unpaired t-test. Error bars: SEM. *P<0.05. **P<0.01. F) Differentially expressed genes between day-3 primed ECs generated from H1 control vs. *SOXF* TKD CRISPRi hPSCs are colored. G) Gene Set Enrichment analysis (GSEA)^97^ of scRNA-seq data from H1 CRISPRi control vs. *SOXF* TKD hPSCs differentiated into day-3 artery ECs, day-3 primed ECs, and day-4 vein ECs. Color represents the log_10_-transformed adjusted P-value. H) *Left*: OmniATAC-seq, CUT&RUN, and bulk RNA-seq of H1 hPSC-derived day-3 primed ECs and day-4 vein ECs. *Right*: scRNA-seq of day-3 artery ECs, day-3 primed ECs, and day-4 vein ECs generated from H1 control vs. *SOXF* TKD hPSCs. *SOXF* genes are required for *CDH5* (*VE-CADHERIN*) expression in primed ECs.

**Figure S10:**
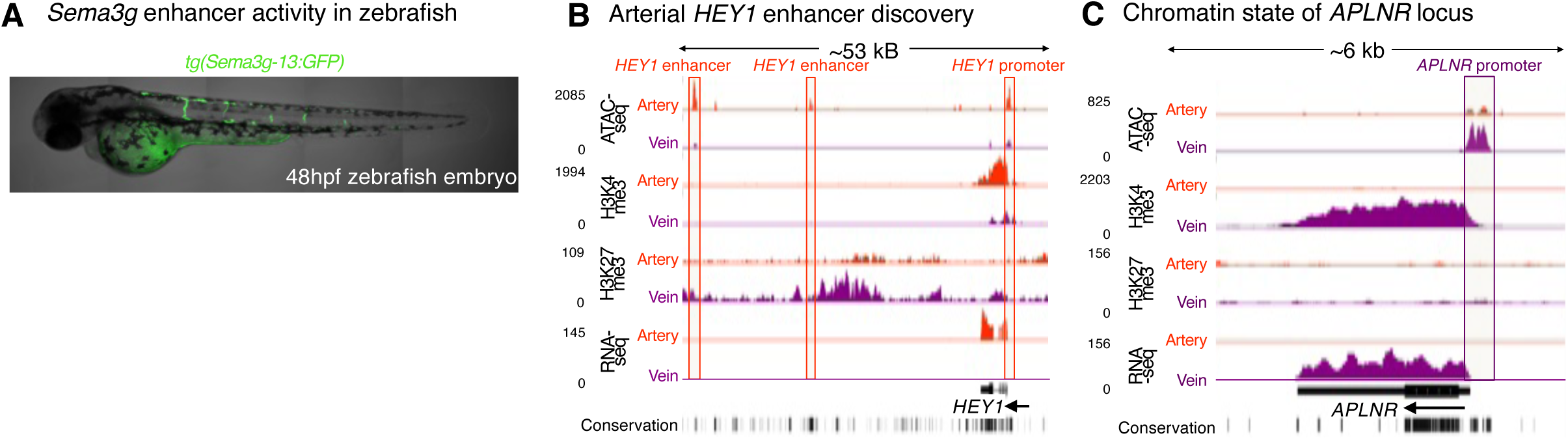
Chromatin hallmarks of human arteriovenous identity. A) Image of a 2 day-post-fertilization (dpf) zebrafish bearing a *Sema3g* -13 kB enhancer driving *GFP* expression. B) OmniATAC-seq, CUT&RUN, and bulk RNA-seq of H1 hPSC-derived day-3 artery ECs and day-4 vein ECs. C) OmniATAC-seq, CUT&RUN, and bulk RNA-seq of H1 hPSC-derived day-3 artery ECs and day-4 vein ECs.

## STAR METHODS

### Key Resources Table

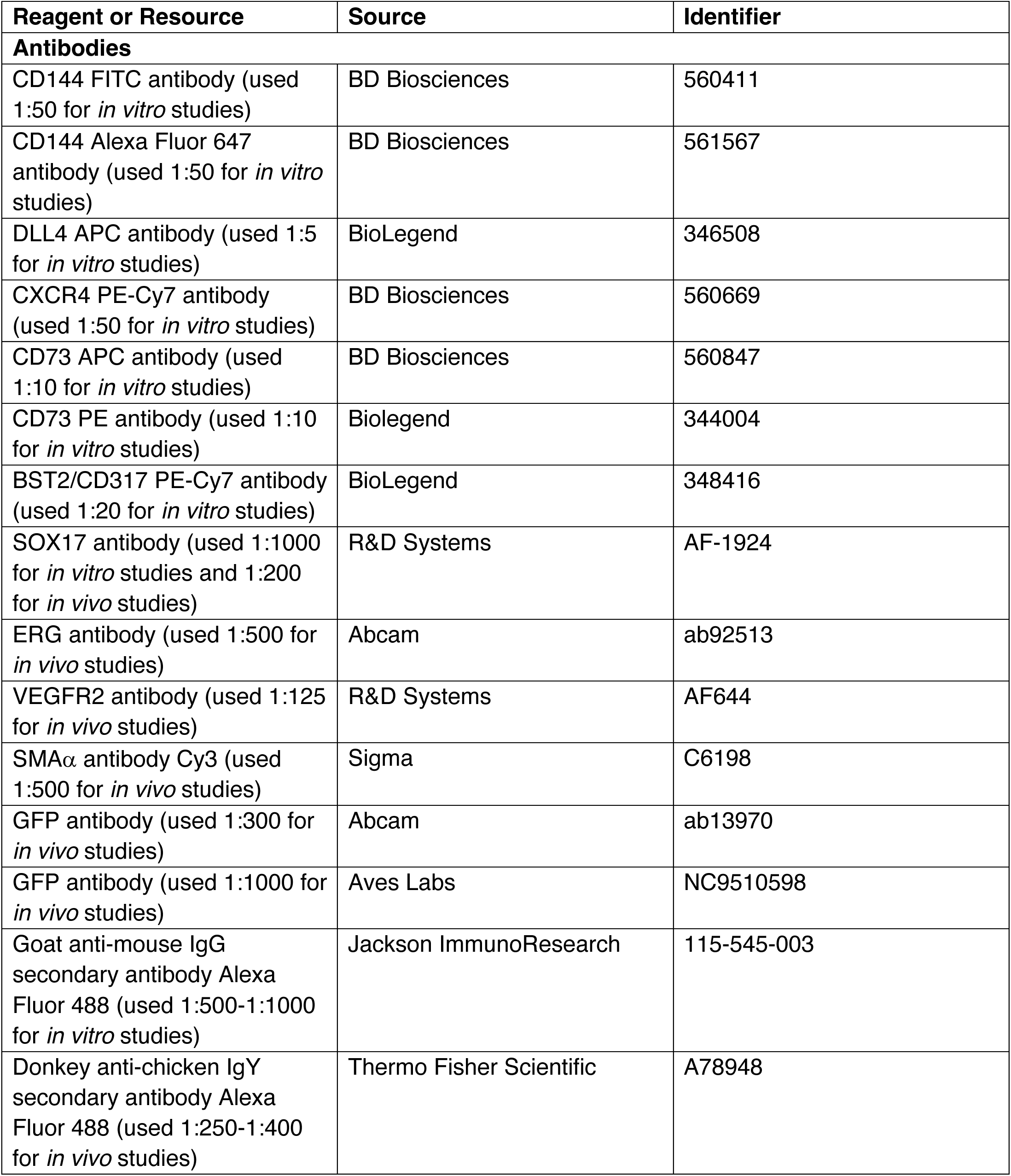
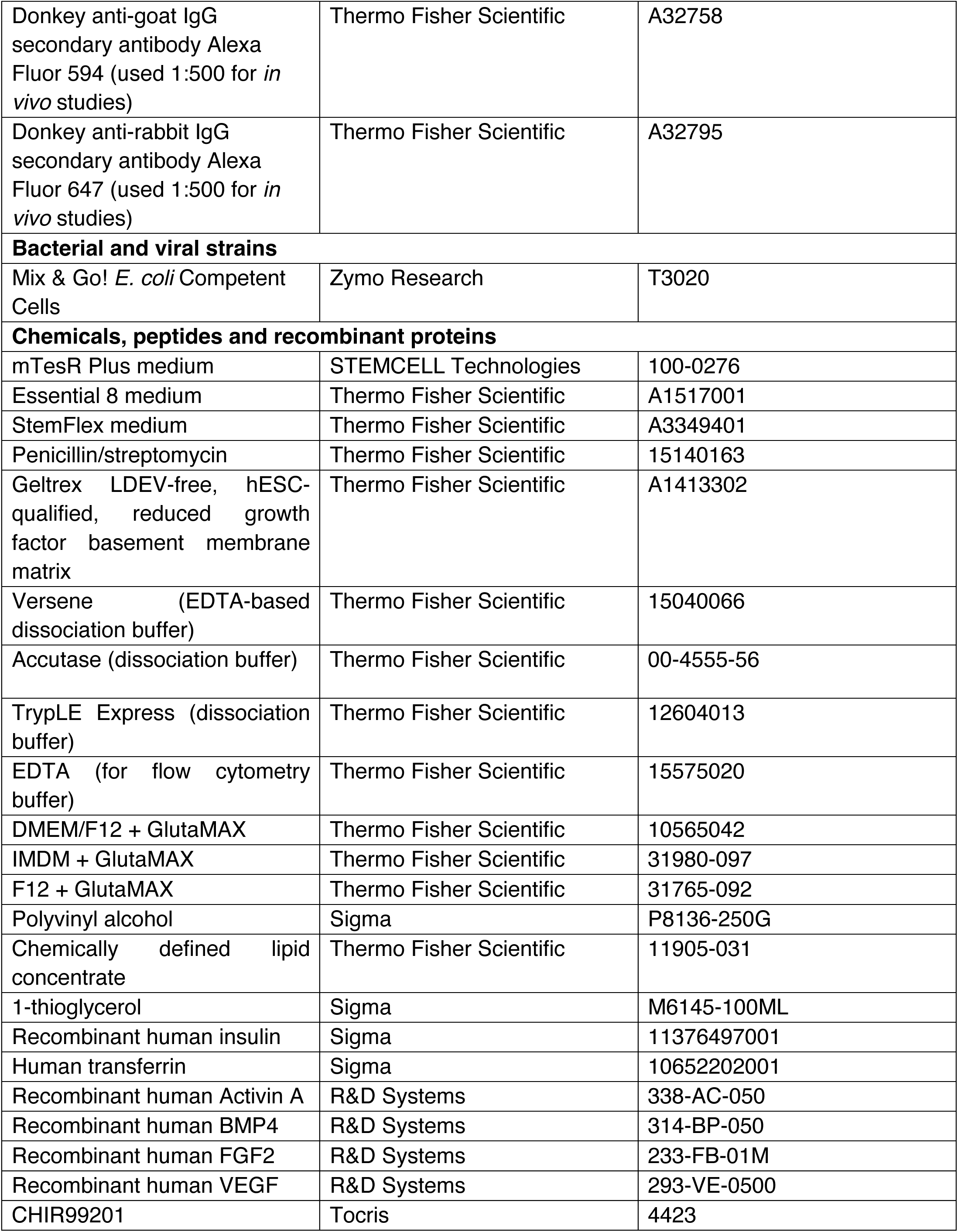
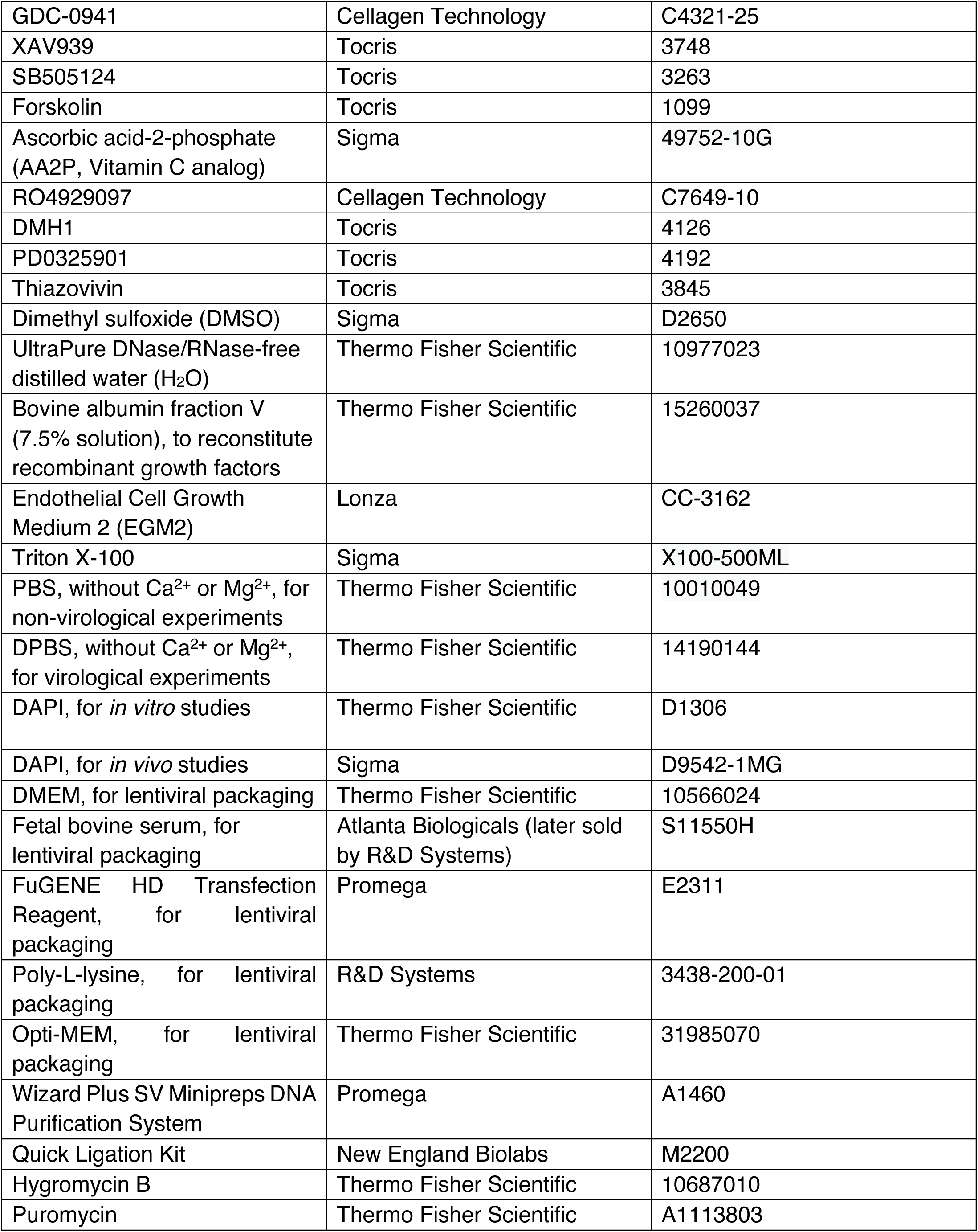
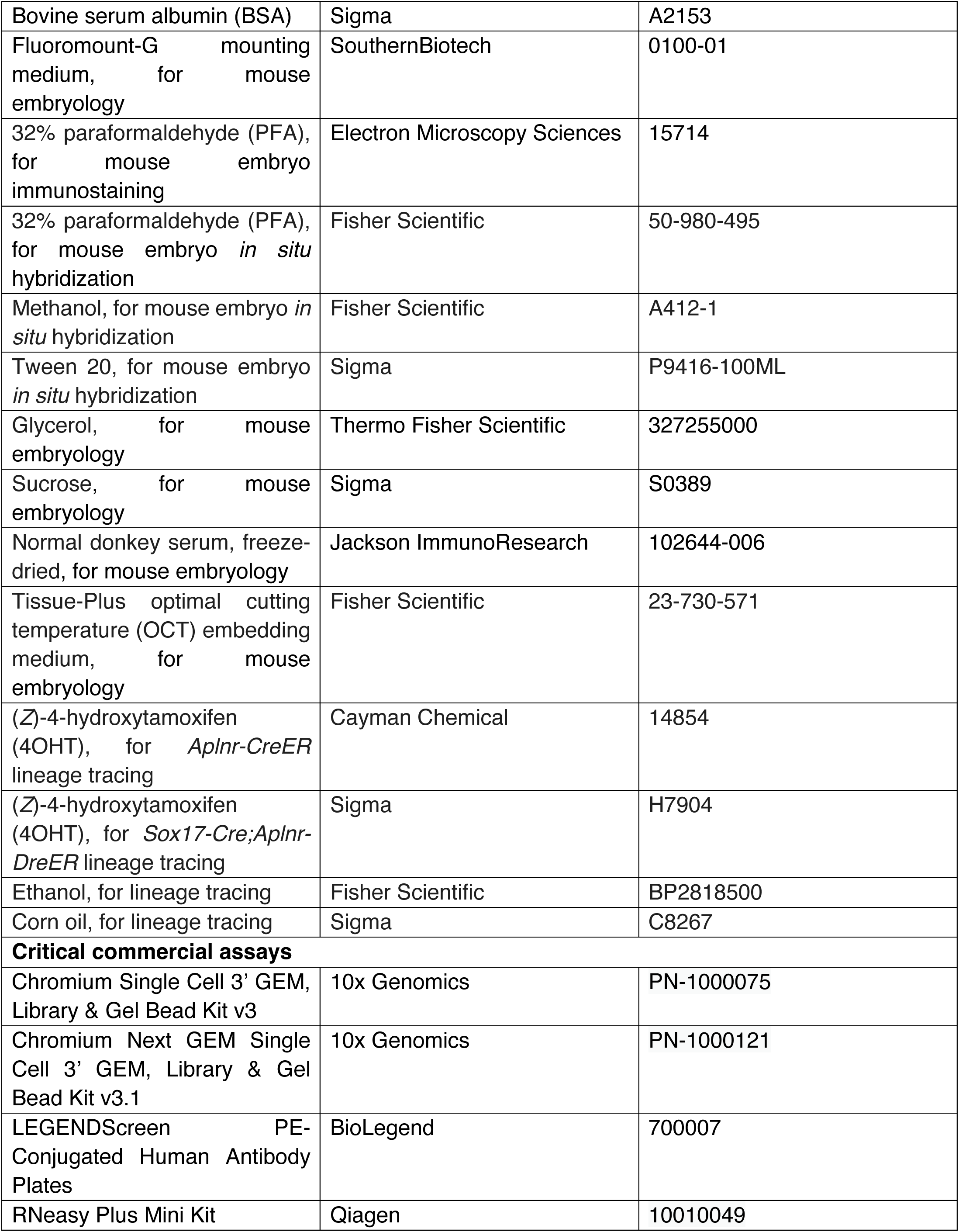
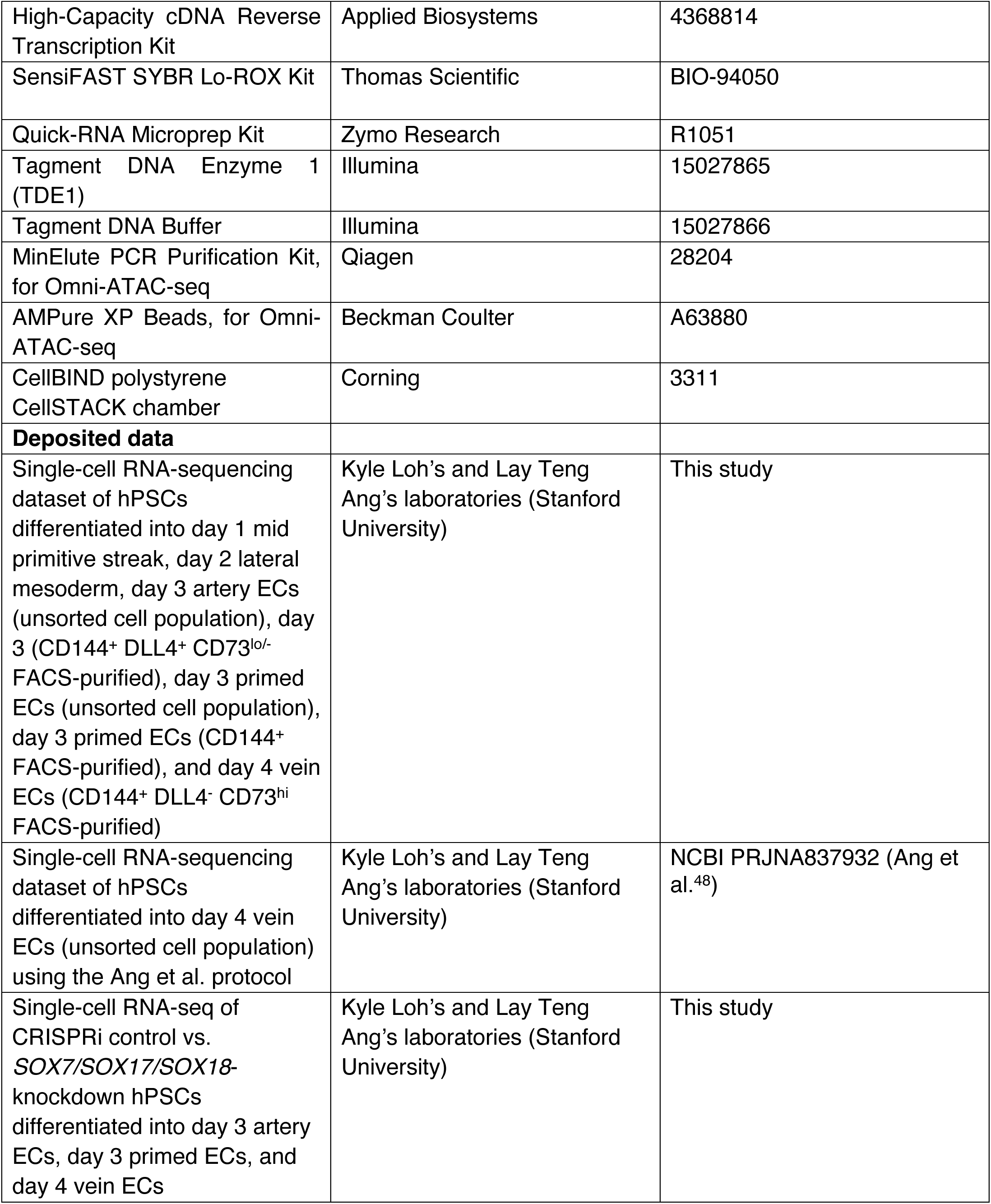
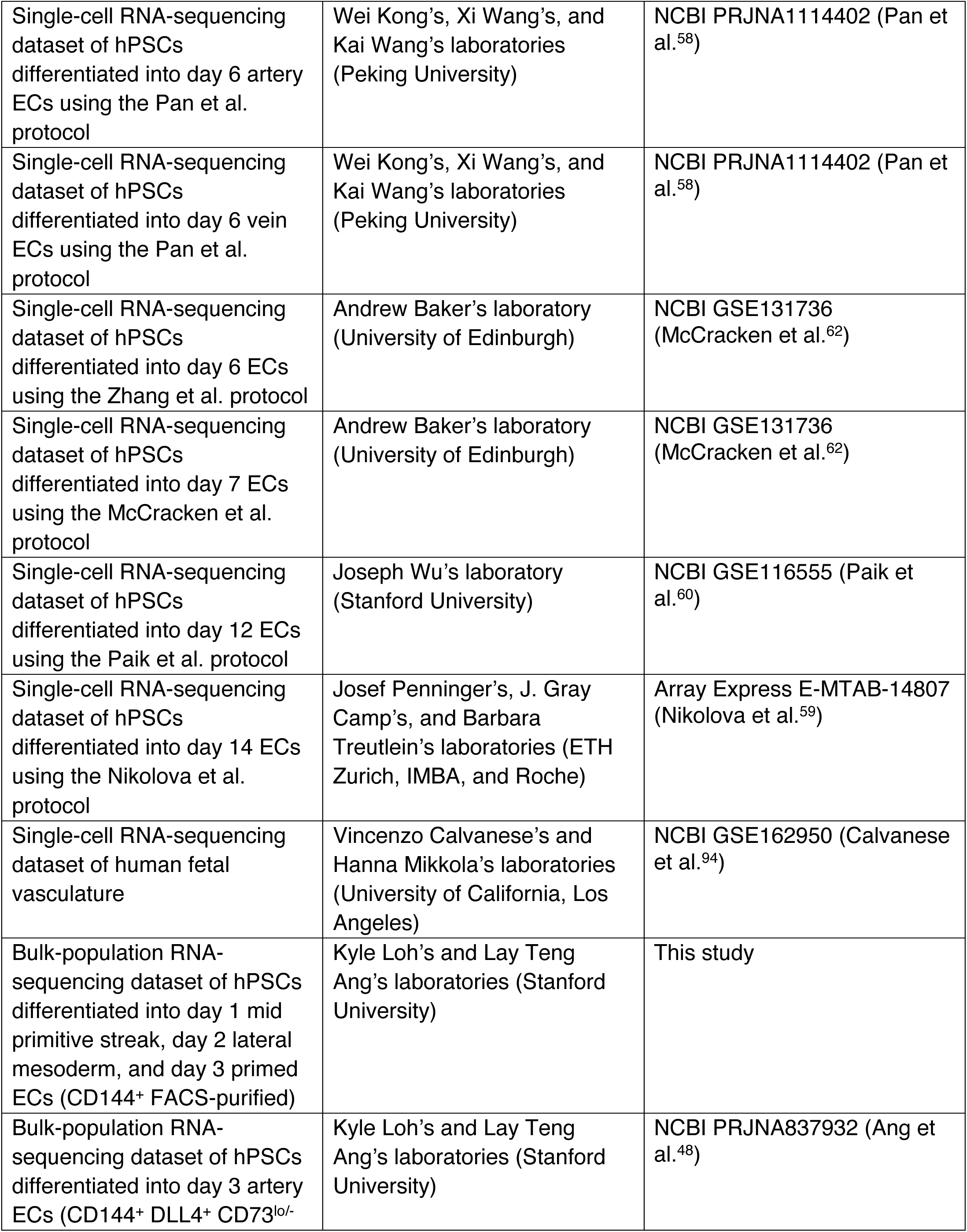
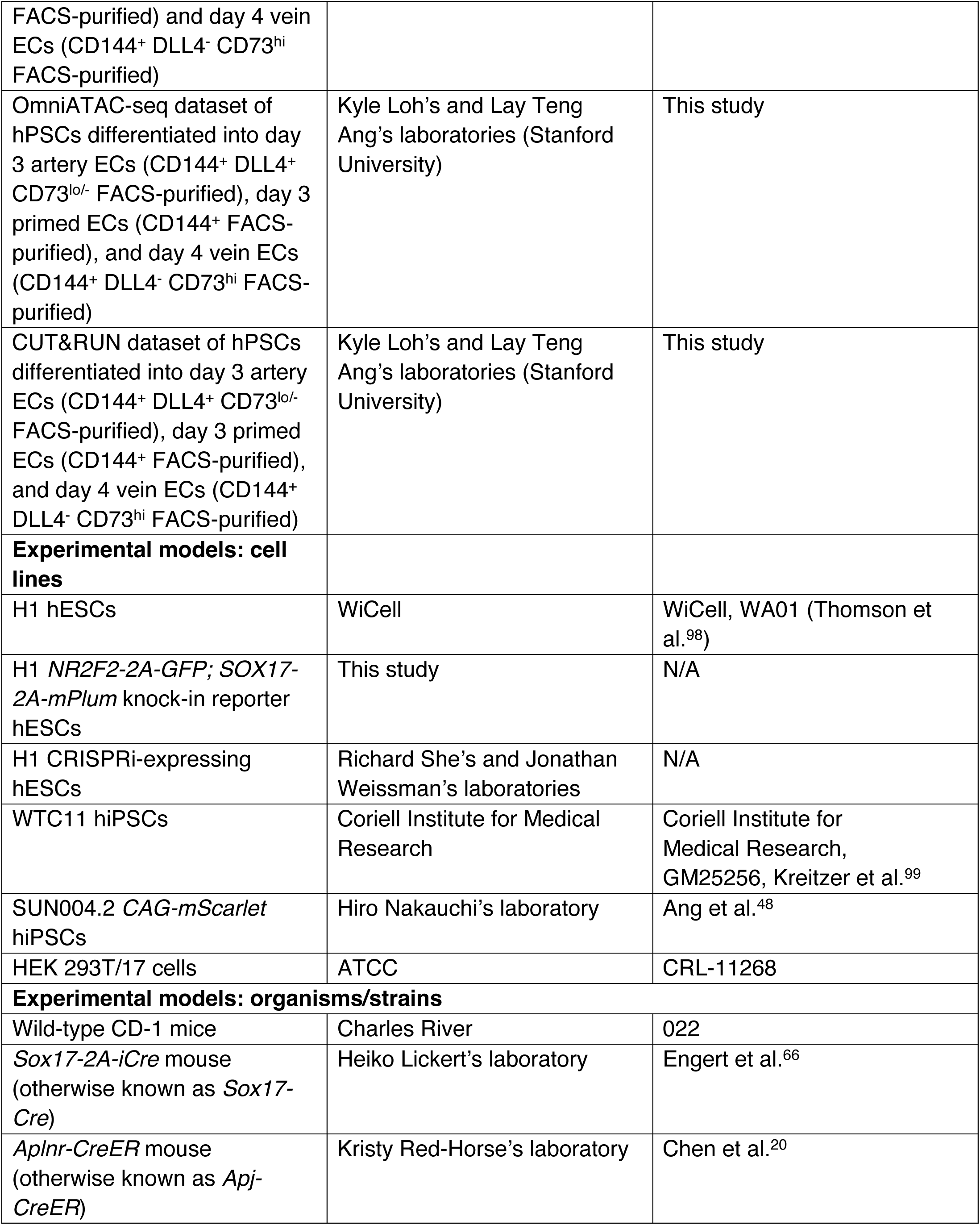
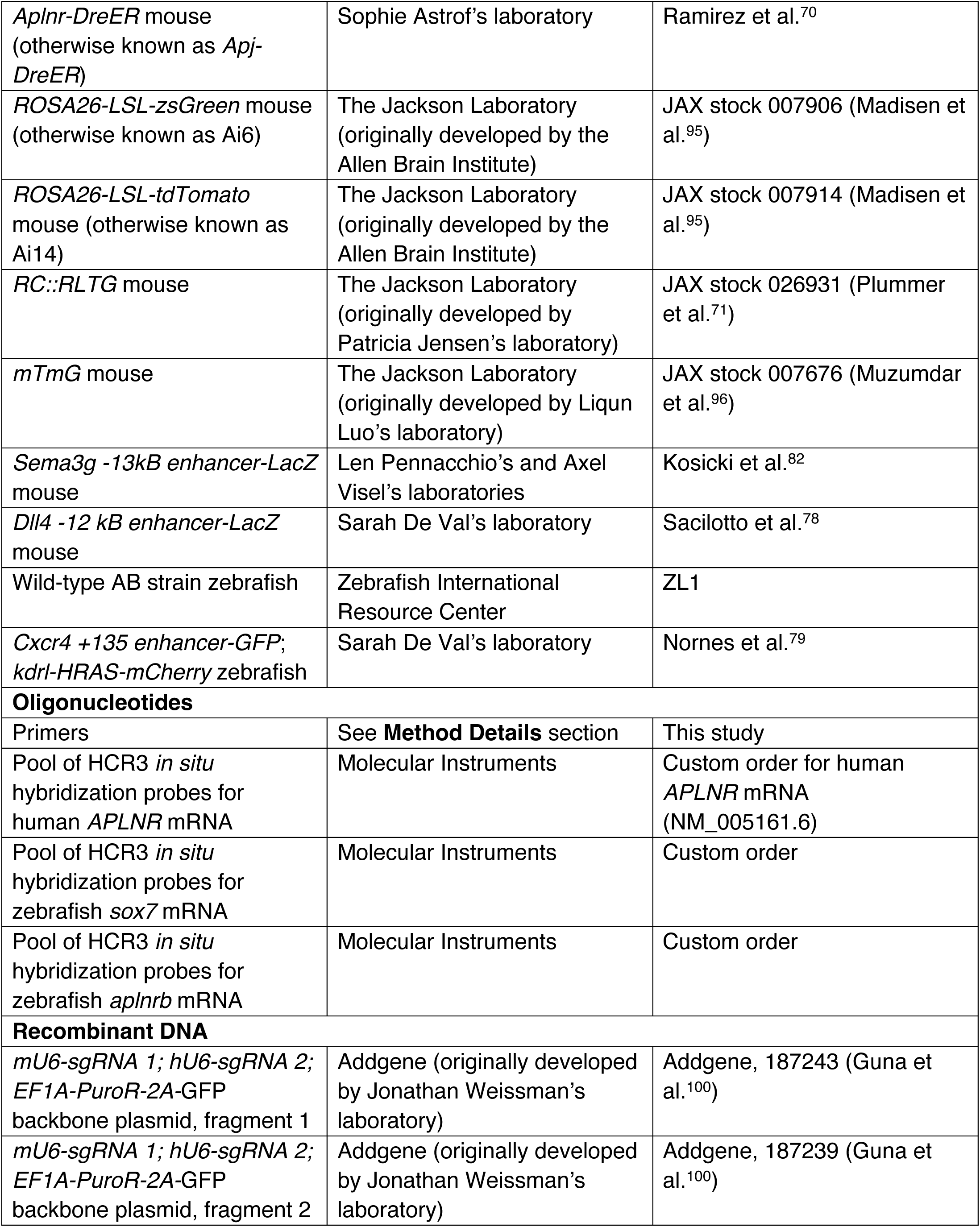
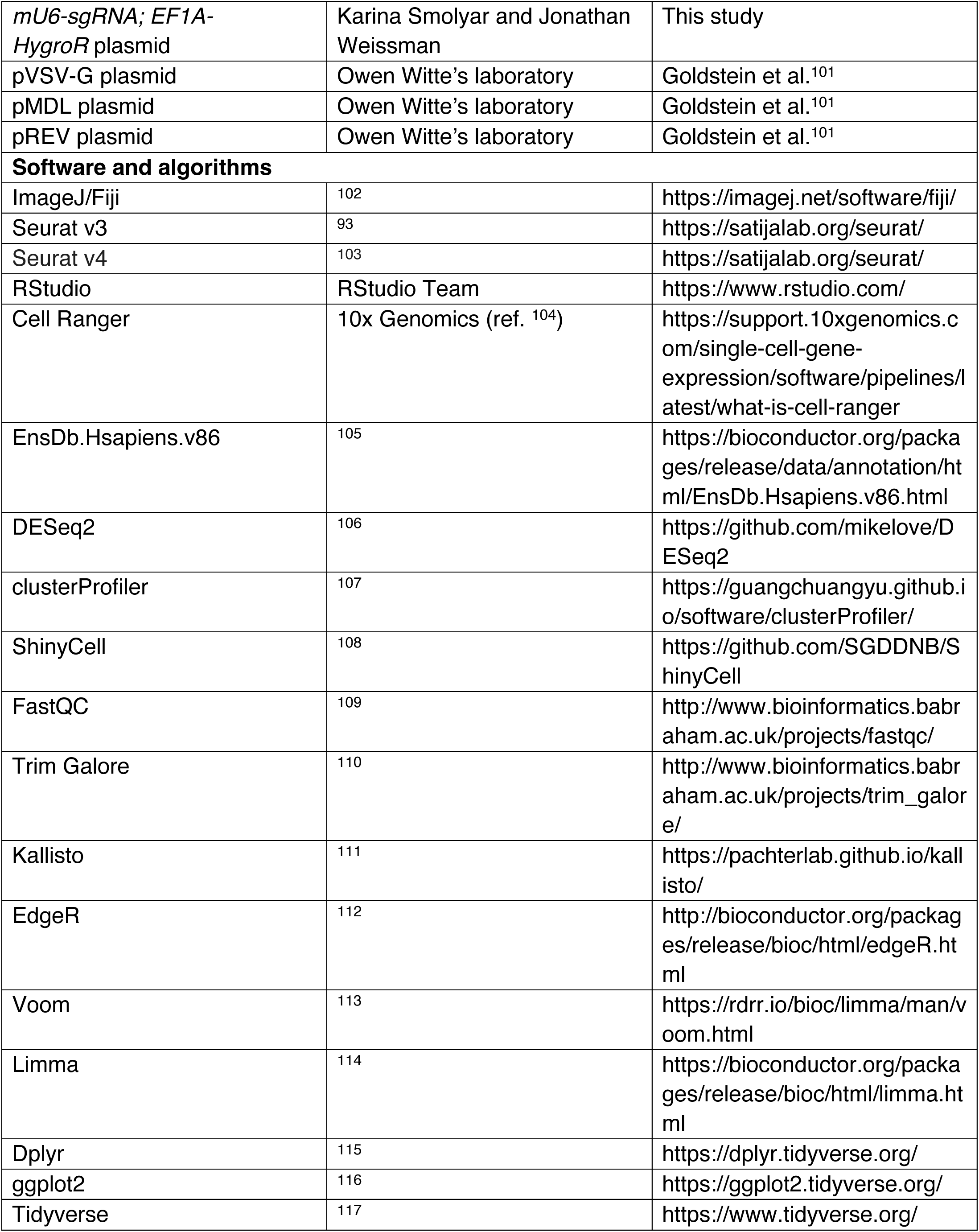
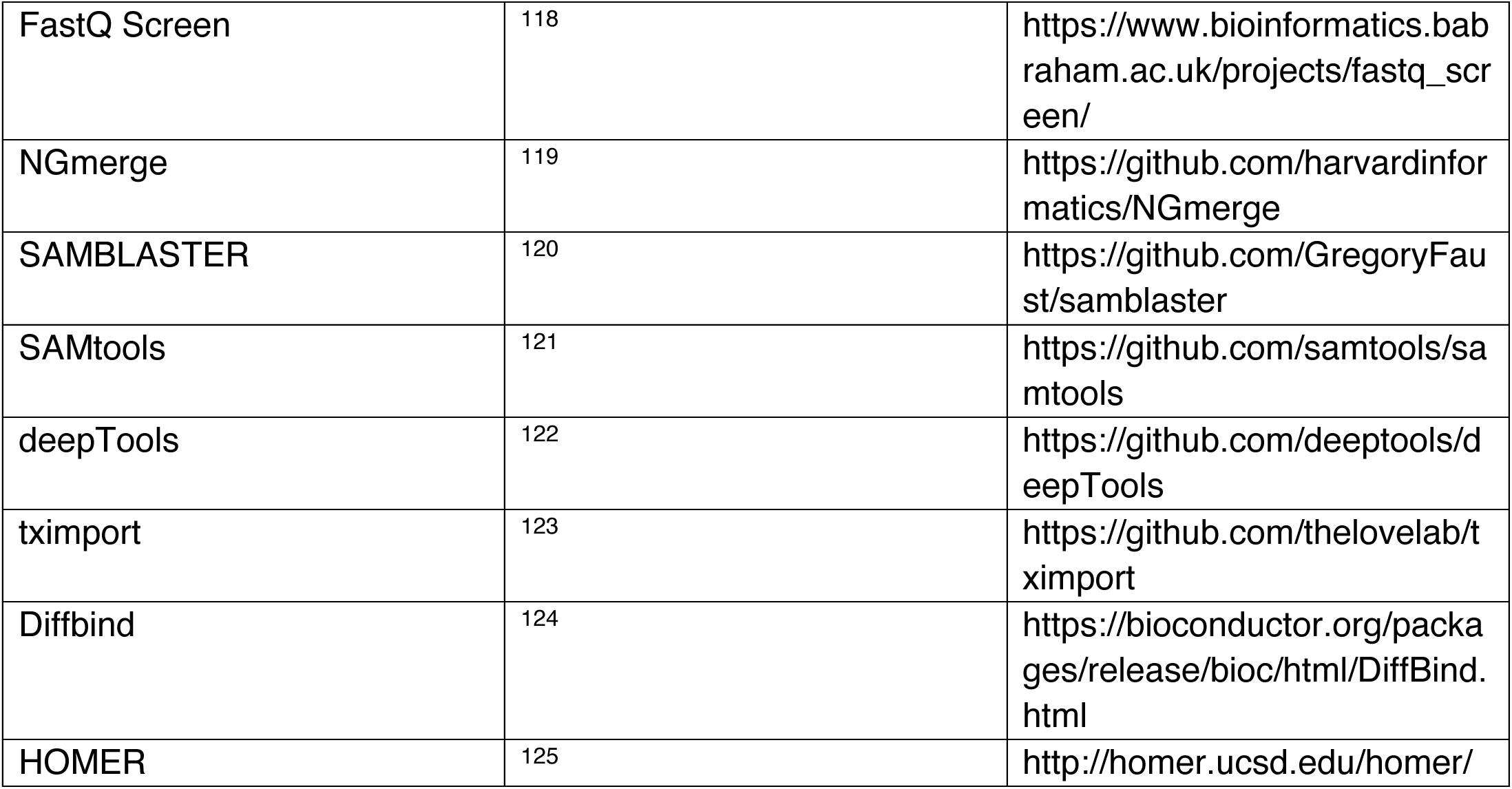

## RESOURCE AVAILABILITY

### Lead contact

Requests for further information and resources should be directed to and will be fulfilled by the lead contact, Lay Teng Ang.

### Materials availability

H1 *SOX17-2A-mPlum*; *NR2F2-2A-GFP* double reporter hESCs were generated as part of this study, and will be made freely available upon request and the completion of applicable Material Transfer Agreements.

### Data and code availability

Bulk-population RNA-seq, single-cell RNA-seq, bulk-population ATAC-seq, and bulk-population CUT&RUN datasets generated in this study will be made available upon acceptance. An interactive portal to access the single-cell RNA-seq dataset of hPSC differentiation into artery and vein ECs is available at https://anglab.shinyapps.io/artery-vein-scrna-seq/. Computational scripts used for genomics analyses conducted in this study are available at GitHub: https://github.com/LayTengAngLab/Prevein_endothelial.

## EXPERIMENTAL MODEL AND SUBJECT DETAILS

### Cell culture

All cells in this study were cultured in standard incubator conditions (20% O_2_, 5% CO_2_ and 37°C).

### Human pluripotent stem cell lines

Wild-type H1 hESCs^98^ (obtained from WiCell; XY biological sex), H1 *NR2F2-2A-GFP; SOX17-2A-mPlum* knock-in reporter hESCs (generated in this study; XY biological sex), CRISPRi-expressing H1 hESCs (generated in this study; XY biological sex), wild-type WTC11 hiPSCs^99^ (obtained from the Coriell Institute for Medical Research; XY biological sex), and SUN004.2 *CAG-mScarlet* hiPSCs^48^ (obtained from Hiro Nakauchi’s laboratory; XY biological sex) were used in this study.

All hPSC lines in this study, except for WTC11 and H1 CRISPRi hPSCs, were cultured in mTeSR Plus medium (STEMCELL Technologies). WTC11 hPSCs were cultured in Essential 8 medium^126^ (Thermo Fisher Scientific). H1 CRISPRi hPSCs were cultured in StemFlex medium (Thermo Fisher Scientific).

Methods to culture undifferentiated hPSCs have been described previously^127^. In brief: undifferentiated hPSCs were propagated in mTeSR Plus medium (STEMCELL Technologies) + 1% penicillin/streptomycin in monolayer cultures, on Geltrex basement membrane matrix-coated plates (described below). To maintain undifferentiated hPSCs, mTeSR Plus was changed either every day or every other day as per the manufacturer’s recommendations. In order to maintain cultures of undifferentiated hPSCs, when they became partially confluent, undifferentiated hPSCs were passaged by treating them for 7 minutes with Versene (Thermo Fisher; an EDTA-based dissociation buffer) at room temperature. Subsequently, Versene was removed, mTeSR Plus was added, and then hPSCs were manually scraped off the plate to generate clumps. hPSC clumps were then seeded onto new plates that had been precoated with Geltrex basement membrane matrix (described below) in mTeSR Plus medium + 1% penicillin/streptomycin. To reiterate, during Versene-based maintenance passaging of undifferentiated hPSCs as clumps, ROCK inhibitor was not added.

### Other cell lines

HEK 293T/17 cells (obtained from ATCC; isolated from tissue of XX biological sex) and Vero C1008 cells (obtained from ECACC; isolated from tissue of XX biological sex) were used in this study.

### Mouse models

Wild-type CD-1 mice (Charles River, catalog number 022), *Sox17-Cre* mice (provided by Heiko Lickert’s laboratory^66^), *Aplnr-CreER* mice (provided by Kristy Red-Horse’s laboratory^20^), *Aplnr-DreER* mice (provided by Sophie Astrof’s laboratory^70^), *ROSA26-LSL-zsGreen* mice (Ai6; JAX stock 007906, and developed by the Allen Brain Institute^95^), *ROSA26-LSL-tdTomato* mice (Ai14; JAX stock 007914, and developed by the Allen Brain Institute^95^), *RC::RLTG* mice (JAX stock 026931, and developed by Patricia Jensen’s laboratory^71^), *Sema3g -13kB enhancer-LacZ* mice (developed by Len Pennacchio’s and Axel Visel’s laboratories^82^), and *Dll4 -12 kB enhancer-LacZ* mice (developed by Sarah De Val’s laboratory^78^), were used in this study.

### Zebrafish models

Wild-type AB zebrafish and *Cxcr4 +135 enhancer-GFP*; *kdrl-HRAS-mCherry* zebrafish (developed by Sarah De Val’s laboratory^79^) were used in this study.

## METHOD DETAILS

### Data processing and visualization

Standard plots were prepared with Microsoft Excel, Microsoft PowerPoint, or GraphPad Prism. Flow cytometry data were visualized with FlowJo. Microscope images were visualized with Fiji^102^. Genomics data were respectively wrangled and plotted using dplyr^115^ and ggplot2^116^ in the tidyverse^117^, in the RStudio environment. Genomics tracks were visualized with the Integrated Genomics Viewer (IGV)^128^, and evolutionary conservation was depicted using the Phastcons 20-way mammalian conservation track. Color palettes were chosen with the assistance of https://colorbrewer2.org/.

### Mouse husbandry

Adult mice of the indicated genotypes were mated to generate timed pregnancies. Females were checked each morning for a vaginal plug; noon on the day a plug was observed was defined as embryonic day 0.5 (E0.5).

### Lineage tracing

- ***Sox17-Cre;Aplnr-DreER* lineage tracing**: As described previously^129^, 25 mg of (*Z*)-4-hydroxytamoxifen (4OHT; Sigma, H7904) was dissolved in 1250 μL ethanol (Fisher Scientific, BP2818500) by vortexing and heating at 60 °C. This yielded a 20 mg/mL stock of 4OHT, which was aliquoted and stored at -20 °C. Prior to dosing mice, 50 μL aliquots (containing 1 mg of 4OHT) were heated for 10 minutes at 65 °C, and then combined with pre-warmed corn oil (250 μL, Sigma, C8267). This mixture of 4OHT and corn oil was thoroughly vortexed. Pregnant females were intraperitoneally injected at the specified labeling timepoint with 1 mg of 4OHT per mouse.
- ***Aplnr-CreER* lineage tracing**: 25 mg of (*Z*)-4-hydroxytamoxifen (4OHT; Cayman Chemical, 14854) was dissolved at 10mg/mL in ethanol (Fisher Scientific, BP2818500) by vortexing, and then was aliquoted and stored at -80 °C. Pregnant female mice were intraperitoneally injected with 28 mg 4OHT per kg of mouse weight at the specified labeling timepoint.

### *In situ* hybridization and immunostaining of whole-mount mouse embryos

Fluorescent *in situ* hybridization (FISH) of whole-mount mouse embryos was performed using hybridization chain reaction v3.0 (HCR3)^130^, and some instances, immunostaining was simultaneously performed. HCR3 was performed as per the Molecular Instruments protocol (https://www.molecularinstruments.com/hcr-v3-protocols), and is briefly summarized here.

Mouse embryos were dissected in ice-cold 4% paraformaldehyde (Fisher Scientific, 50-980-495, diluted in PBS). They were then subsequently fixed overnight in 4% paraformaldehyde, sequentially dehydrated in methanol solutions of increasing concentration (Fisher Scientific, A412-1), and then incubated in 100% methanol overnight.

Embryos were subsequently permeabilized in PBS + 0.1% Triton X-100 for 1 hour at room temperature, and then blocked in blocking buffer (PBS + 0.05% Triton X-100 + 5% BSA) overnight at 4 °C. Embryos were then stained with primary antibodies (diluted in blocking buffer) for 24-72 hours at 4 °C, washed three times with PBS, and stained with secondary antibodies (diluted in blocking buffer) for 24 hours at 4 °C. Subsequently, embryos were washed three times with PBS, and then HCR3 was performed as per the Molecular Instruments protocol. Hybridization mRNA probes, amplifiers, and buffers were obtained from Molecular Instruments.

Embryos were incubated in DAPI + SSCT (sodium chloride, sodium citrate, and Tween buffer) prior to mounting. Images were captured on an Olympus FV3000 confocal microscope.

### Immunostaining of whole-mount mouse embryos

Mouse embryos were dissected in ice-cold PBS, and then fixed in 4% paraformaldehyde (Electron Microscopy Sciences, 15714; diluted in PBS) for 1 hour at 4 °C. Subsequently, embryos were washed in PBS and stained with primary antibodies (diluted in PBS + 0.1% Triton X-100) overnight at 4 °C. Following three washes in PBS + 0.1% Triton X-100, embryos were incubated in secondary antibodies (diluted in PBS + 0.1% Triton X-100) overnight at 4 °C. Embryos were then washed in PBS + 0.1% Triton X-100, counterstained with 1 μg/ml DAPI (Sigma, D9542), and washed twice in PBS. Embryos were then equilibrated sequentially in 25%, 50%, and 75% glycerol in PBS for 30 minutes at room temperature without agitation. Embryos were then mounted in Fluoromount-G (SouthernBiotech, 0100-010) and imaged on a Zeiss LSM 980 confocal microscope.

### Immunostaining of cryosectioned mouse embryos

Mouse embryos were dissected in ice-cold PBS, and then fixed in 4% paraformaldehyde (Electron Microscopy Sciences, 15714; diluted in PBS) for 1 hour at 4 °C. Subsequently, embryos were washed in PBS, and dehydrated in 30% sucrose at 4°C overnight. Embryos were embedded in OCT medium (Fisher Scientific, 23-730-571), and then frozen overnight at -80 °C. Embryos were sectioned to a thickness of 20 μm and incubated at room temperature for 30 minutes. Sections were subsequently permeabilized in PBS + 0.1% Triton X-100 for 15 minutes at room temperature and blocked in 5% donkey serum + PBS + 0.05% Triton X-100 at room temperature. Sections were then stained with primary antibodies (diluted in blocking buffer) at 4 °C overnight, washed with PBS + 0.05% Triton X-100. They were then stained with secondary antibodies (diluted in blocking buffer) at room temperature, followed by additional washes in PBS + 0.05% Triton X-100, counterstained with 1 μg/ml DAPI (Sigma, D9542) for 10 minutes at room temperature, and then washed with PBS. Slides were then mounted with Fluoromount-G (SouthernBiotech, 0100-010) and imaged on a Zeiss LSM 980 confocal microscope.

### Quantification of mouse embryo images

Fiji^102^ was used to generate maximum-intensity projections from z-stacks of whole-mount mouse embryos or sectioned mouse embryos.

- Sox17 immunostaining: Erg+ or Sox17+ cells were manually counted in the dorsal aorta (DA) and cardinal vein (CV). The percentage of Erg+ cells that expressed Sox17 is shown.
- *Sox17-Cre;Aplnr-DreER* lineage tracing: Erg+ or GFP+ cells were manually counted in the DA and CV. The percentage of Erg+ cells that expressed GFP is shown.
- *Aplnr-CreER* lineage tracing: Erg+, Sox17+, or tdTomato+ cells were manually counted in the DA and CV. The percentage of Erg+ cells that co-expressed both Sox17 and tdTomato is shown.

### Functional testing of *Dll4* enhancer element in mouse embryos

A stable transgenic mouse line bearing a genomic integration of the *Dll4* -12 kb enhancer element driving *LacZ* reporter expression, known as *Tg(Dll4-12:lacZ)*, was generated as previously described^78^. Embryos were fixed in 2% paraformaldehyde + 0.2% glutaraldehyde + 1x PBS for 60 minutes. After fixation, embryos were rinsed in 0.1% sodium deoxycholate, 0.2% Nonidet P-40, 2 mM MgCl_2_ and 1x PBS, then stained for 2-24 hours in 1 mg/ml 5-bromo-4-chloro-3-indolyo-β-D-galactoside solution (X-gal) containing 5 mM potassium ferrocyanide, 5 mM ferricyanide, 0.1% sodium deoxycholate, 0.2% Nonidet P-40, 2 mM MgCl_2_, and 1x PBS. After staining, embryos were rinsed through a series of 1x PBS washes, then fixed overnight in 4% paraformaldehyde at 4°C. All embryos were imaged using a Leica M165C stereo microscope equipped with a ProGres CF Scan camera and CapturePro software (Jenoptik).

### Functional testing of *Sema3g* enhancer element in mouse embryos

The activity of the *Sema3g* -13 kb enhancer element was tested in mouse embryos through a transgenic reporter assay by Len Pennacchio’s and Axel Visel’s laboratories, and was reported as part of the VISTA Enhancer Browser^82,83^ (VISTA Enhancer Browser ID hs2179: https://enhancer.lbl.gov/vista/element?vistaId=hs2179&alleleId=0&backbone=hZR&stage=e11.5). In brief, a reporter construct containing the *Sema3g* -13 kb enhancer element and the *Hsp68* minimal promoter driving *LacZ* reporter expression was randomly integrated into the mouse genome; whole-mount staining of mouse embryos for *LacZ* reporter expression was then performed^82,83^.

### Fluorescent *in situ* hybridization of zebrafish embryos

FISH of whole-mount zebrafish embryos was performed using HCR3^130^.

Wild-type AB strain embryos were fixed overnight with 4% paraformaldehyde (BT-Fix) in PBS at 4 °C, dehydrated through a sequential ethanol series, and stored at -20°C. Embryos were rehydrated and washed three times with PBT (PBS + 0.2% bovine serum albumin + 0.2% Tween 20). A prehybridization step was done using hybridization buffer (30% formamide + 5x SSC + 9 mM citric acid at pH 6.0 + 0.1% Tween 20 + 50 μg/ml heparin + 1x Denhardt’s solution, + 10% dextran sulfate) for 30 mins at 37 °C.

Following this, each 2 pmol of each HCR3 probe was combined with 500 μL of hybridization buffer and zebrafish embryos were incubated overnight in this solution at 37°C, while being gyrated. Following incubation, embryos were washed four times with 30% formamide + 5x SSC + 9 mM citric acid at pH 6.0 + 0.1% Tween 20 + 50 μg/mL heparin at 37°C, for 15 minutes each. This was then followed with three washes with 5x SSCT (5x SSC + 0.1% Tween 20) at room temperature for 5 minutes each. Subsequently, 30 minutes of incubation in amplification buffer (5× SSC + 0.1% Tween 20 + 10% dextran sulfate) at room temperature was performed. Hairpin probes (30 pmol each), fluorescently labeled through snap cooling of 3 μM stock solution, were added to the embryos in amplification buffer and incubated overnight at room temperature in the dark. Subsequently, samples were washed five times with 5x SSCT. Embryos were mounted in 0.6% low-melting agarose and imaged using a Nikon A1R confocal microscope and a 20× objective. Images were processed using the denoiseAI function in the NIS Elements software (Nikon) to reduce noise. Maximum intensity images were obtained with Fiji^130^.

### Differentiation of hPSCs into artery and vein ECs

Two differentiation protocols were used throughout this study, which we refer to “Version 1” (V1, which was used for most experiments) and “Version 2” (V2, which was only used in **Extended Fig. 7A-F**). The V1 differentiation protocol has been described previously^48,127^. All RNA-seq, ATAC-seq, CUT&RUN, and LEGENDScreen profiling in this study was conducted on cells generated by the V1 differentiation protocol.

The V2 differentiation protocol was developed as part of this study and is more efficient at generating both artery and vein ECs than the V1 protocol. The V2 differentiation protocol is identical to the V1 protocol, with the exception that the V1 protocol entails 24 hours of lateral mesoderm differentiation, whereas the V2 protocol entails 40 hours of lateral mesoderm differentiation.

All differentiation was conducted in defined, serum-free CDM2 basal media^131,132^. Detailed instructions to how to prepare CDM2 are available^127^.

- **Seeding hPSCs for differentiation** (**Step 0**). In preparation for differentiation, hPSCs were dissociated into single cells using Accutase (Thermo Fisher), because sparse seeding of cells is important for differentiation. Accutase-dissociated hPSCs were plated into recipient wells in mTeSR medium supplemented with the ROCK inhibitor thiazovivin (1 μM, Tocris; to enhance hPSC survival after passaging) onto plates precoated with Geltrex basement membrane matrix. hPSCs were seeded at a density of 25,000-50,000 cells/cm^2^ were seeded (i.e., 95,000-190,000 hPSCs/well of a 12-well plate)^48,127^. To clarify, long-term maintenance of undifferentiated hPSCs entailed passaging as clumps using Versene (an EDTA-based dissociation buffer; to maintain normal karyotype), but hPSCs were dissociated using Accutase to seed single cells for differentiation. 24 hours after seeding in mTeSR + 1 μM thiazovivin, during which the hPSCs re-formed small clumps, differentiation was initiated as described below.
- **Step 1: Mid primitive streak induction, 24 hours**. Day 0 hPSCs were briefly washed (DMEM/F12, Thermo Fisher) to remove all traces of mTeSR + thiazovivin. Then, they were differentiated towards mid primitive streak in CDM2 media supplemented with Activin A (30 ng/mL, R&D Systems), BMP4 (40 ng/mL, R&D Systems), CHIR99021 (6 μM, Tocris), FGF2 (20 ng/mL, Thermo Fisher), as previously described^48,127,132^. In both the V1 and V2 protocols, mid primitive streak induction was conducted for 24 hours.
- **Step 2: Lateral mesoderm induction, 24-40 hours**. Day 1 mid primitive streak cells were briefly washed (DMEM/F12) and then differentiated towards lateral mesoderm in CDM2 media supplemented with BMP4 (40 ng/mL), GDC-0941 (2.5 μM, Cellagen Technology), Forskolin (10 μM, Tocris), SB-505124 (2 μM, Tocris), VEGF (100 ng/mL, R&D Systems), XAV939 (1 μM, Tocris) and ascorbic acid-2-phosphate (AA2P; 200 μg/mL, Sigma), as previously described^48,127^. In the V1 protocol, lateral mesoderm induction was performed for 24 hours. In the V2 protocol, lateral mesoderm induction was performed for 40 hours. Subsequently, lateral mesoderm was subjected to either artery EC induction (Step 3A; below) or primed EC induction (Step 3A; below).
- **Step 3A: Artery EC induction, 24 hours**. Day 2 lateral mesoderm cells (V1 protocol) or day 2.67 lateral mesoderm cells (V2 protocol) were briefly washed (DMEM/F12) and then differentiated towards artery ECs in CDM2 media supplemented with Activin A (15 ng/mL), DMH1 (250 nM, Tocris), GDC-0941 (2.5 μM), VEGF (100 ng/mL), XAV939 (1 μM) and AA2P (200 μg/mL), as previously described^48,127^. In both the V1 and V2 protocols, artery EC induction was performed for 24 hours.
- **Step 3B: Primed EC induction, 24 hours**. Day 2 lateral mesoderm cells (V1 protocol) or day 2.67 lateral mesoderm cells (V2 protocol) were briefly washed (DMEM/F12) and then differentiated into primed ECs in CDM2 media supplemented with SB505124 (2 μM), DMH1 (250 nM), RO4929097 (2 μM, Cellagen Technology), VEGF (100 ng/mL), XAV939 (1 μM) and AA2P (200 μg/mL), as previously described^48,127^. In both the V1 and V2 protocols, primed EC induction was performed for 24 hours.
- **Step 4B: Vein EC induction, 24 hours**. Day 3 primed ECs (V1 protocol) or day 3.67 primed ECs (V2 protocol) were briefly washed (DMEM/F12) and then differentiated into vein ECs in CDM2 media supplemented with SB505124 (2 μM), RO4929097 (2 μM), PD0325901, (500 nM, Tocris), CHIR99021 (1 μM) and AA2P (200 mg/mL), as previously described^48,127^. In both the V1 and V2 protocols, vein EC induction was performed for 24 hours.

Detailed methods to reconstitute each differentiation-inducing small molecule and recombinant growth factor and to prepare stocks of each are available^127^. In brief, (1) all recombinant growth factors were reconstituted in PBS + 0.1% bovine albumin fraction V (both from Thermo Fisher Scientific), (2) all small molecules except for AA2P were reconstituted in DMSO (Sigma), and (3) AA2P was reconstituted in H_2_O (Thermo Fisher Scientific), as described previously^127^.

### Large-scale differentiation of hPSCs into artery and vein ECs

Large-scale differentiation of hPSCs was performed using the V2 differentiation protocol described above.

- **Artery EC differentiation**: WTC11 hPSCs were seeded at a density of 20.5K cells/cm^2^ in a Geltrex-coated 5-stack CellSTACK device (Corning), which yielded 94.5 million cells per device.
- **Vein EC differentiation**: WTC11 hPSCs were seeded at a density of 25.2K cells/cm^2^ in a Geltrex-coated 5-stack CellSTACK device, which yielded 239.6 million cells per device. Alternatively, WTC11 hPSCs were seeded at a density of 17.2K cells/cm^2^ in 60 15-cm dishes, which yielded 654 million cells altogether.

### Cryopreservation and maintenance of hPSC-derived artery and vein ECs

After hPSC differentiation into artery ECs or vein ECs as described above, they could be maintained for at least 6 additional days *in vitro* on Geltrex-coated cell culture plates. As described previously^48^, hPSC-derived artery ECs were expanded in EGM2 (Endothelial Cell Growth Medium 2, Lonza CC-3162), which was refreshed every 24 hours. By contrast, hPSC-derived vein ECs were expanded in EGM2 + SB505124 (2 μM) + RO4929097 (2 μM) + Forskolin (10 μM), which was refreshed every 24 hours.

Alternatively, as previously described^48^, hPSC-derived day 3 artery ECs and day 4 vein ECs were dissociated, cryopreserved in freezing media (90% PBS + 10% DMSO), and stored in liquid nitrogen. hPSC-derived artery and vein ECs were then thawed in their respective media (EGM2 for artery ECs and EGM2 + SB505124 + RO4929097 + Forskolin for vein ECs) and cultured for up to 6 days as described above, with Thiazovivin (1 μM) added for the first 24 hours post-thawing to improve cell survival.

### Flow cytometry

Cultured cells were dissociated by incubation in TrypLE Express (Thermo Fisher) for 5 minutes at 37 °C. Following dissociation, the cells were diluted with 5-10 times excess volume of FACS buffer (PBS + 1 mM EDTA [Thermo Fisher] + 2% v/v FBS [Atlanta Bio] + 1% v/v Penicillin/Streptomycin [Thermo Fisher]) and centrifuged at 500g for 5 minutes to pellet them. Each cell pellet was then resuspended in FACS buffer and incubated with fluorescently-conjugated primary antibodies for 15-30 minutes in the dark at 4°C. After staining, cells were washed twice with FACS buffer and resuspended in 100 μL FACS buffer containing DAPI (1 μg/mL) for live/dead discrimination. Flow cytometry was conducted on a Beckman Coulter CytoFlex analyzer in the Stanford Stem Cell Institute FACS Core Facility. Flow cytometry data analysis was performed using FlowJo software. Cells were gated based on forward and side scatter area, followed by height and width parameters for doublet discrimination. Subsequently, live cells that were negative for DAPI were gated for marker analyses and calculations of population frequency. Gates to determine the percentage of cells positive for a given marker were set based on negative controls, namely cells that were not stained with an antibody or cells that were lacking the genetically encoded fluorescent reporter of interest.

In this study, we defined hPSC-derived artery and vein ECs using the following cell-surface marker combinations:

- **Artery ECs**: CD144^+^ CXCR4^+^ DLL4^+^ (and, in some experiments, CD144^+^ DLL4^+^ CD73^lo/-^). The following antibody combination was used to define arterial identity: CD144 FITC (BD Biosciences, 560411 [1:50 concentration]), DLL4 APC (Biolegend, 346508 [1:5 concentration]), and CXCR4 PE-Cy7 (BD Biosciences, 560669 [1:50 concentration]).
- **Vein ECs**: CD144^+^ CD317^+^ CD73^+^ (and, in some experiments, CD144^+^ DLL4^-^ CD73^hi^). The following antibody combination was used to define venous identity: CD144 FITC (BD Biosciences, 560411 [1:50 concentration]), CD73 APC (BD Biosciences, 560847 [1:10 concentration]), and CD317 PE-Cy7 (BioLegend, 348416 [1:20 concentration]).

The CD73 protein is encoded by the *NT5E* gene. The CD317 protein is encoded by the *BST2* gene.

### Quantitative PCR

Methods for RNA extraction, reverse transcription, and qPCR have been described previously^48^. In brief, undifferentiated or differentiated hPSCs were first lysed in 350 μL of RLT Plus Buffer and RNA was extracted using the RNeasy Plus Mini Kit (Qiagen) according to the manufacturer’s protocol. Second, 300 ng of total RNA was reverse transcribed into cDNA using the High-Capacity cDNA Reverse Transcription Kit (Applied Biosystems) according to the manufacturer’s protocol. Third, qPCR was performed in 384-well format using the SensiFAST SYBR Lo-ROX Kit (Thomas Scientific) as previously described^48,132^, using gene-specific forward and reverse primers on a QuantStudio 5 qPCR machine (Thermo Fisher). Expression of all genes was normalized to the levels of the reference gene *YWHAZ*.

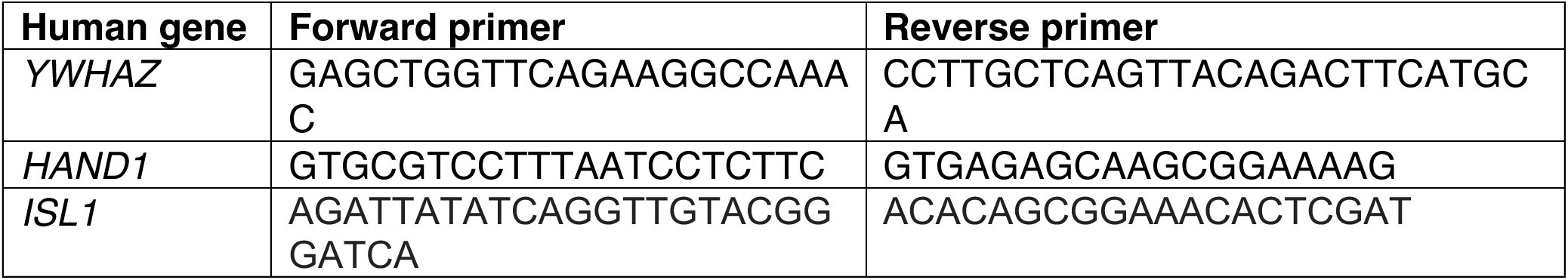
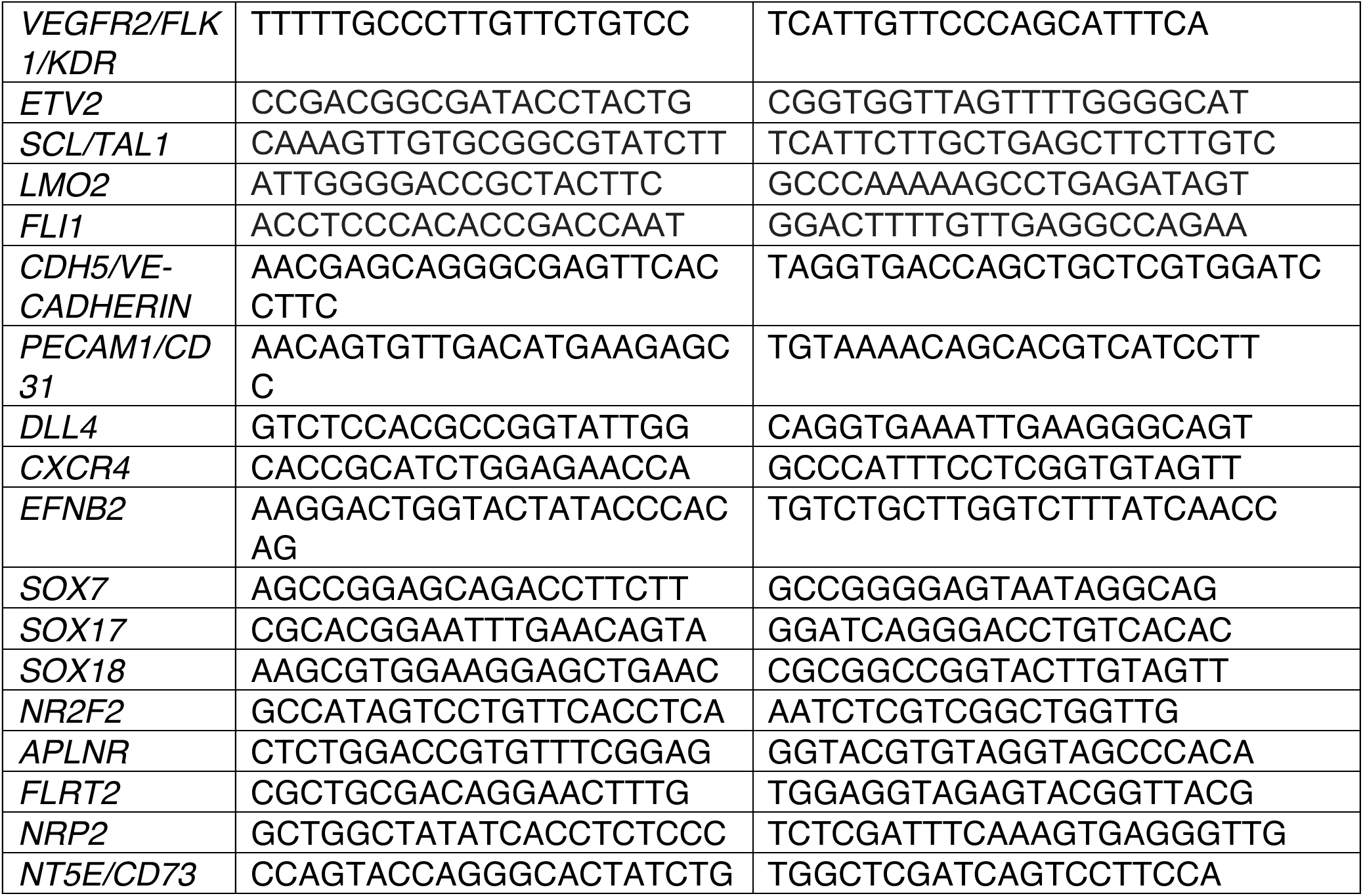

### Combined immunostaining and *in situ* hybridization of cultured cells

Combined immunostaining and *in situ* hybridization of cultured cells was performed as described by Molecular Instruments (https://files.molecularinstruments.com/MI-Protocol-2%C2%BAIF-RNAFISH-GenericSolution-Rev6.pdf). First, cultured monolayer cells were fixed, permeabilized and immunostained with primary and secondary antibodies. Next, probe hybridization, amplification, and wash steps were performed using the HCR3 protocol^130^. Imaging was conducted using an FV3000 confocal microscope (Olympus).

### Bulk-population RNA-seq

We performed bulk-population RNA-seq on the following cell populations that were generated from H1 hPSCs in the same biological experiment: *1*) day-0 hPSCs (entire cell population, unsorted); *2*) day-1 mid primitive streak (entire cell population, unsorted); *3*) day-2 lateral mesoderm (entire cell population, unsorted); *4*) day-3 artery ECs (FACS-purified for CD144^+^ DLL4^+^ CD73^lo/-^ cells; 10 μM forskolin was added during day-3 artery EC induction); *5*) day-3 primed ECs (FACS-purified for CD144^+^ cells); and *6*) day-4 vein ECs (FACS-purified for CD144^+^ cells). Bulk-population RNA-seq datasets of day-3 artery ECs and day-4 vein ECs were previously reported by Ang et al., 2022 (NCBI PRJNA837932)^48^.

Bulk-population RNA-seq was performed and computationally analyzed as described previously^48^. Cells were lysed in Zymo RNA lysis buffer, and RNA was purified using the Zymo Quick-RNA Microprep Kit (Zymo, R1051). RNA integrity was assessed by Agilent Bioanalyzer on-chip electrophoresis. High-quality RNA samples with RNA integrity number ≥ 8 underwent poly(A) enrichment, and libraries were prepared with indexed adaptors for multiplexing. RNA-seq libraries were then sequenced on the DNBSEQ-G400 sequencer by BGI Global Genomic Services to generate 150-bp paired-end reads. To limit batch effects, all libraries were pooled prior to sequencing and distributed across multiple planes.

FastQC^109^ was used to perform quality control of raw RNA-seq reads. Adapters and low-quality bases were trimmed with Trim Galore^110^; a Phred quality threshold of ≥ 33 was used, and reads shorter than 20 nucleotides after trimming were discarded. Reads from each library—which was originally sequenced on multiple lanes—were then concatenated to yield one file per library. RNA-seq reads were pseudoaligned to human reference genome hg38, and gene-level RNA-seq counts were then quantified, using Kallisto^111^. After quantification of gene-level RNA-seq counts, two separate computational workflows in the RStudio environment were used:

- **Workflow 1** (**Generation of volcano plots**): Gene counts were filtered with edgeR^112^. Counts were then transformed to log_2_ counts per million (CPM) using voom^113^. Differentially expressed genes were determined by limma^114^, and P values were adjusted for multiple hypothesis testing using the Benjamini-Hochberg method to control the false discovery rate.
- **Workflow 2** (**Generation of gene expression matrices**): Ensembl transcript IDs were mapped to Ensembl gene IDs using EnsDb.Hsapiens.v86^105^. Transcript-level abundance estimates were summarized to gene-level counts with tximport^123^, and then imported into DESeq2^106^ for differential expression analysis.

### Single-cell RNA-sequencing of hPSC-derived cell populations

The 10x Genomics Chromium platform was used to perform single-cell RNA-seq profiling every 24 hours during the differentiation of H1 hPSCs into artery and vein ECs using the V1 differentiation protocol.

In one experiment, the Chromium Single Cell 3’ GEM, Library & Gel Bead Kit v3 was used to profile the following samples: *1*) day-0 hPSCs (entire cell population, unsorted); *2*) day-1 mid primitive streak (entire cell population, unsorted); *3*) day-2 lateral mesoderm (entire cell population, unsorted); *4*) day-3 primed EC (entire cell population, unsorted); *5*) day-3 primed EC (FACS-purified for CD144^+^ cells); *6*) day-4 vein EC (entire cell population, unsorted); and *7*) day-4 vein EC (FACS-purified for CD144^+^ DLL4^-^ CD73^hi^ cells). In a second experiment, the Chromium Single Cell 3’ GEM, Library & Gel Bead Kit v3.1 was used to profile the following samples: *8*) day-3 artery EC (entire cell population, unsorted); and *9*) day-3 artery EC (FACS-purified for CD144^+^ DLL4^+^ CD73^lo/-^ cells). This scRNAseq dataset of unsorted day-4 vein ECs was previously reported by Ang et al., 2022 (NCBI PRJNA837932)^48^; all other scRNAseq datasets were generated as part of this study.

The rationale to perform scRNAseq of FACS-purified ECs was to rigorously test whether cell-surface marker combinations would enable the isolation of transcriptionally-homogeneous cell populations.

As per the manufacturer, no batch effects have been detected between the Chromium v3 and v3.1 chemistries (https://kb.10xgenomics.com/hc/en-us/articles/360047373071-Does-Cell-Ranger-distinguish-between-v3-and-v3-1-chemistry).

Sequencing libraries were prepared using Chromium Single Cell 3’ GEM Gene Expression v3 or v3.1 kits as per the manufacturer’s guidelines. Sequencing libraries were diluted in Buffer EB. Libraries were prepared with 10-nucleotide indices compatible with Illumina sequencers.

### Computational analysis of single-cell RNA-sequencing data from hPSC-derived cell populations

scRNAseq libraries were sequenced across multiple lanes on an Illumina HiSeq 4000 sequencer by Novogene. The first and last 8 nucleotides of the i7 indices were unique and thus used for demultiplexing. FASTQ sequencing files were input into Cell Ranger^104^, which was used to align reads to the hg38 reference genome (version GRCh38-2024-A), followed by filtering, barcode counting, and unique molecular identifier (UMI) counting.

Subsequent analyses were performed using Seurat v3^93^ in the RStudio environment. Cell matrix files generated from Cell Ranger were imported into R using the Seurat function “Read10x_h5”. For each individual scRNAseq dataset, we performed quality control by excluding dying/dead cells that *1*) exhibited low numbers of expressed genes, *2*) displayed anomalously low or high mitochondrial counts, or *3*) did not express reference genes *ACTB* or *YWHAZ*; additionally, we also *4*) computationally excluded likely doublets by removing cells that had significantly higher counts of expressed genes. High-quality single-cell transcriptomes that passed these quality control metrics were used for subsequent analyses.

Seurat objects from all scRNAseq datasets were merged using the “Merge” function in Seurat. scRNAseq data were then normalized using the “LogNormalized” function and scaled using a linear transformation in Seurat. Marker gene expression was depicted on UMAP plots^133^ by coloring each single cell (i.e., a dot) according to the levels of marker gene expression. Dots were randomly ordered, without visually superimposing dots that were positive for a given marker gene, thereby avoiding visual stacking bias.

For scRNAseq analysis of control vs. *SOXF*-deficient CRISPRi hPSCs that were subject to EC differentiation, first we computationally identified ECs and excluded non-ECs from the scRNAseq dataset. This was performed via Louvain clustering, which clearly distinguished *PECAM1*+ ECs vs. *PECAM1*- non-EC clusters (**Fig. S9D**). EC clusters were used for subsequent analyses, using the aforementioned scRNAseq analysis workflows. Gene set enrichment analysis of gene ontology terms was performed on genes that were differentially expressed between control vs. *SOXF*-CRISPRi ECs using clusterProfiler^107^. Selected gene ontology terms were shown.

Data wrangling and plotting were respectively performed using dplyr^115^ and ggplot2^116^ in the tidyverse^117^. An interactive web browser to explore scRNAseq data was constructed using ShinyCell^108^.

### Comparing different endothelial differentiation protocols using single-cell RNA-seq

We analyzed scRNAseq datasets of hPSCs that were subjected to 8 different EC differentiation protocols. The 8 differentiated cell populations analyzed were:

- **Ang et al., day 3 artery ECs**: hPSCs were differentiated into artery ECs in 3 days using the V1 differentiation protocol described by Ang et al., 2022^48^. This new scRNAseq dataset was generated as part of this study.
- **Ang et al., day 4 vein ECs**: hPSCs were differentiated into vein ECs in 4 days using the V1 differentiation protocol described by Ang et al., 2022^48^. This scRNAseq dataset was deposited by the Ang et al., 2022 study^48^ and is publicly available from NCBI PRJNA837932.
- **Pan et al., day 6 artery ECs**: hPSCs were differentiated into artery ECs in 6 days using the Pan et al., 2024 protocol^58^. This scRNAseq dataset was deposited by the Pan et al., 2024 study^58^, and is publicly available from NCBI PRJNA1114402.
- **Pan et al., day 6 vein ECs**: hPSCs were differentiated into vein ECs in 6 days using the Pan et al., 2024 protocol^58^. This scRNAseq dataset was deposited by the Pan et al., 2024 study ^58^, and is publicly available from NCBI PRJNA1114402.
- **Zhang et al., day 6 ECs**: hPSCs were differentiated into ECs in 6 days using the Zhang et al., 2017 differentiation protocol^61^. This scRNAseq dataset was deposited by the McCracken et al., 2019 study^62^, and is publicly available from NCBI GSE131736.
- **McCracken et al., day 7 ECs**: hPSCs were differentiated into ECs in 7 days using the McCracken et al., 2019 protocol^62^. This scRNAseq dataset was deposited by the McCracken et al., 2019 study^62^, and is publicly available from NCBI GSE131736. Data from 3 experimental replicates were deposited to the Gene Expression Omnibus, and our preliminary analysis revealed batch effects among these three experimental replicates. To reduce batch effects, we selected replicate 3 for analysis, as it was the replicate that contained the largest number of cells. Of note, the original McCracken et al. study refers to cells being harvested on day 8 of differentiation^62^. However, because the first day of differentiation in their procedure entails cell seeding in hPSC medium^62^, to be consistent with the nomenclature used here to describe other differentiation protocols, here we refer to these cells being differentiated for 7 days.
- **Paik et al., day 12 ECs**: hPSCs were differentiated into ECs in 12 days using the Paik et al., 2018 protocol^60^. This scRNAseq dataset was deposited by the Paik et al., 2018 study^60^, and is publicly available from NCBI GSE116555.
- **Nikolova et al., day 14 ECs**: hPSCs were differentiated into vascular organoids in 14 days using the Nikolova et al., 2025 protocol^59^, which in turn was modified from the Wimmer et al., 2019 protocol^134^ to generate vascular organoids. This scRNAseq dataset was deposited by the Nikolova et al., 2025 study^59^, and is publicly available from ArrayExpress E-MTAB-14807.

scRNAseq datasets were analyzed using Seurat v3^93^. For all scRNAseq datasets, first we selected single-cell transcriptomes that passed well-established quality control metrics, by excluding dying/dead cells that *1*) exhibited low numbers of expressed genes, *2*) displayed anomalously low or high mitochondrial counts, or *3*) did not express reference genes *ACTB* or *YWHAZ*; additionally, we also *4*) computationally excluded likely doublets by removing cells that had significantly higher counts of expressed genes. For initial analyses, we assessed all single-cell transcriptomes that passed these quality control criteria; we did not pre-select a given subpopulation of cells based on marker gene expression that might bias further analyses.

Seurat objects from all scRNAseq datasets were then merged using the “Merge” function in Seurat v3^93^. To quantify the degree of cellular heterogeneity generated by each differentiation protocol, Louvain clustering was applied at the same resolution (0.1) across all datasets, thus decomposing each scRNAseq dataset into multiple constituent cell-types. To assign the identity of each “cell-type” within each dataset, we analyzed genes that were differentially expressed between each cell-type within a given dataset, which we annotated based on the expression of known marker genes:

- **Ang et al., day 3 artery ECs** (3 constituent clusters): artery EC (*EDN1*+ *MKI67*-), dividing artery EC (*EDN1*+ *MKI67+*), mesenchyme (*ACTC1*+ *TPM1*+)
- **Ang et al., day 4 vein ECs** (4 constituent clusters): vein EC *(FLRT2*+ *NEFH*+), dividing vein EC (*FLRT2*+ *DGKB*+), other EC (*LNCAROD*+), mesenchyme (*ACTC1*+ *TPM1*+)
- **Pan et al., day 6 artery ECs** (4 constituent clusters): EC (*EGFL7*+ *ASPM*-), dividing EC (*EGFL7*+ *ASPM*+), mesenchyme (*IGFBP3*+ *TOP2A-*), dividing mesenchyme (*IGFBP3*+ *TOP2A*+)
- **Pan et al., day 6 vein ECs** (3 constituent clusters): EC (*KDR*+ *TOP2A*-), dividing EC (*KDR*+ *TOP2A*+), mesenchyme (*ACTC1*+)
- **Day 6 Zhang et al.** (5 constituent clusters): EC (*CLDN5*+ *UBE2C*-), dividing EC (*CLDN5*+ *UBE2C*+), mesenchyme (*MEST*+ *ACTC1*+), blood-like (*SPI1/PU.1*+), heart/kidney (*NKX2.5*+ *LHX1*+)
- **McCracken et al., day 7 ECs** (3 constituent clusters): EC (*PLVAP*+ *MKI67*-), dividing EC (*PLVAP*+ *MKI67*+), mesenchyme (*MEST*+ *TAGLN*+)
- **Paik et al., day 12 ECs** (5 constituent clusters): EC (*ECSCR*+), mesenchyme (*LUM*+ *CENPF*-), proliferating mesenchyme (*LUM*+ *CENPF*+), liver-like (*FGB+ APOA2*+ *TTR*+ *AFP*+), unknown (co-expression of epithelial marker *EPCAM* and mesenchymal marker *LUM*)
- **Nikolova et al., day 14 ECs** (3 constituent clusters): EC (*CLDN5*+), mesenchyme (*LUM*+ *TOP2A*-), dividing mesenchyme (*LUM*+ *TOP2A*+)

### Assessing arteriovenous identity of cells generated from each hPSC differentiation protocol, using single-cell RNA-seq

We quantified the arteriovenous identity of hPSC-derived ECs that were profiled by aforementioned scRNAseq studies. Each dataset comprises a mixture of ECs and non-ECs at varying proportions. First, we computationally selected the EC cluster generated by each differentiation protocol, as per the above cluster annotations. (For differentiation protocols that generated “EC” and “dividing EC” clusters, all EC clusters were combined.) Then, in these EC populations obtained from distinct differentiation protocols, we analyzed the expression of arterial vs. venous marker gene modules, which are referred to as the “artery signature” or “vein signature” in this study. The Hou et al., 2022 study^24^ previously demonstrated that the expression of these arteriovenous gene modules is evolutionarily conserved across ECs obtained from both human and mouse embryos:

- **Arterial gene module**: GJA4, UNC5B, DLL4, MECOM, HEY1, EFNB2, EPAS1, CXCR4, IGFBP3
- **Venous gene module**: NR2F2, NRP2, APLNR, FLRT2

We implemented the AddModuleScore function of Seurat v3^93^ to calculate the average expression of these arterial module genes and venous module genes in hPSC-derived ECs generated from each differentiation protocol.

### Single-cell RNA-seq analysis of endothelial heterogeneity in human and mouse embryos

We downloaded scRNAseq datasets of ECs isolated from the Carnegie Stage 17 human embryo (generated by Calvanese et al., 2022^94^; NCBI GSE162950, sample GSM4968834). Computational analysis was performed as described above, with the exception that Seurat v4^103^ was used.

### OmniATAC-seq library construction

We performed OmniATAC-seq profiling of hPSC-derived day 3 artery ECs (FACS-purified for CD144^+^ DLL4^+^ CD73^lo/-^ cells) and day 4 vein ECs (FACS-purified for CD144^+^ DLL4^-^ CD73^hi^ cells). A slightly modified version (https://www.med.upenn.edu/kaestnerlab/assets/user-content/documents/ATAC-seq-Protocol-(Omni)-Kaestner-Lab.pdf) of the original OmniATAC-seq protocol^49^ was employed here, and is briefly summarized below.

First, we prepared resuspension buffer (10 mM Tris-HCl, pH 7.5 + 10 mM NaCl + 3 mM MgCl_2_ + nuclease-free H_2_O), cold lysis buffer (resuspension buffer + 0.1% v/v NP-40 + 0.1% v/v Tween-20 + 0.01% v/v digitonin), and wash buffer (99.9% resuspension buffer + 0.1% v/v Tween-20).

50,000 cells from each hPSC-derived cell-type were pelleted and washed with 500 μL of cold PBS, before lysis in 100 μL cold lysis buffer for 3 minutes on ice. To the cell lysate, 1 mL of cold wash buffer was added. The mixture was centrifuged at 500g for 10 minutes at 4 °C. Then, the supernatant (cytoplasm) was discarded and the pellet (nuclei) was retained. Transposition reaction mix from the Nextera DNA library prep kit (1x Tagment DNA [TD] Buffer + 1x PBS + 0.1% v/v Tween-20 + 0.01% v/v Digitonin + Tn5 Transposase [Tagment DNA Enzyme 1] + nuclease-free H_2_O) was added to the pellet to resuspend nuclei. The transposition reaction was incubated at 37 °C for 30 minutes on a thermal mixer, with shaking at 1,000 rpm.

DNA was purified using the Qiagen MinElute Reaction Cleanup Kit, and then PCR-amplified using Illumina i5 and i7 index primers on a thermal cycler. Then, the libraries were purified using AMPure XP beads. All OmniATAC-seq libraries were multiplexed such that they could be sequenced as a pool on a single lane. After quality control was performed on pooled libraries, deep sequencing was performed on an Illumina HiSeq sequencer (∼350 million reads/lane) and an Illumina NovaSeq S4 sequencer (∼2500 million reads/lane). According to general guidelines, a minimum of 50 million reads are needed to identify accessible chromatin elements and 200 million reads are needed to identify enriched transcription factor motifs (https://www.illumina.com/techniques/popular-applications/epigenetics/atac-seq-chromatin-accessibility.html) by ATAC-seq. In this study, OmniATAC-seq libraries were sequenced at a depth of 292-1908 million raw reads per library.

### OmniATAC-seq computational analysis

OmniATAC-seq data were computationally processed using the standardized ENCODE ATAC-seq analysis pipeline (https://www.encodeproject.org/atac-seq/). First, reads were aligned to human reference genome hg38 using Bowtie2^135^. Then, MACS2^136^ was used to call peaks for each library. A unified peak list for each cell-type was generated by selecting only peaks that were reproducible between the two replicates. This was achieved through an irreproducible discovery rate (IDR) analysis at the threshold of 0.05 described by the ENCODE Consortium^137^.

Finally, peaks that overlapped with a “black list” of artifactual regions in hg38 (https://sites.google.com/site/anshulkundaje/projects/blacklists) were removed.

Diffbind^124^ was used to identify chromatin regions that exhibited >8-fold differential accessibility between artery and vein ECs. HOMER^125^ was used to discover DNA motifs overrepresented in these artery- or vein-accessible elements. HOMER analyses were run on repeat-masked hg38 sequences extracted in 200-nucleotide windows centered on peak summits, using the “findMotifsGenome.pl” function.

### CUT&RUN library construction

CUT&RUN profiling of H3K4me1, H3K4me3, H3K27me3, and SOX17 was conducted on H1 hPSCs differentiated into day 3 artery ECs, day 3 primed ECs, and day 4 vein ECs. For most CUT&RUN experiments, CD144+ ECs were purified by FACS to exclude any contaminating mesenchymal cells, with the exception for SOX17 CUT&RUN, as SOX17 is not expressed in mesenchymal cells. As a negative control, an isotype IgG control antibody was also separately included.

CUT&RUN profiling and library construction was performed as previously described^55^. CUT&RUN libraries were sequenced on an Illumina NovaSeq X Plus by Novogene.

### CUT&RUN computational analysis

FastQC^109^ and FastQ Screen^118^ were used to perform quality control of raw CUT&RUN reads. Paired-end reads were merged using NGmerge^119^ and aligned to human reference genome hg38 with Bowtie2^64^, using the --very-sensitive mode and fragment length parameters -I 10 -X 2000. Resulting SAM files were processed with SAMBLASTER^120^ to remove PCR duplicates. SAMtools^121^ was then used to convert files into the sorted BAM format. Bigwig signal tracks were generated using the “bamCoverage” function from deepTools^122^, with 5-nucleotide bin size and values displayed in reads per kilobase million (RPKM).

### High-throughput surface marker screen by flow cytometry

The expression of 332 cell-surface markers was assessed across undifferentiated hPSCs (day 0), primitive streak (day 1), lateral mesoderm (day 2), artery ECs (day 3) and vein ECs (day 4) through the use of high-throughput flow cytometry as described previously^132^. In brief, hPSCs or their differentiated mesoderm progeny were dissociated using TrypLE Express. They were then plated into individual wells of four 96-well LEGENDScreen PE-Conjugated Human Antibody Plates (Biolegend, 700001). Each well containing a distinct antibody against a human cell-surface antigen, altogether totaling 332 unique cell-surface markers across four 96-well plates. High-throughput cell-surface marker staining was largely done as per the manufacturer’s recommendations, and cells were stained with a viability dye (DAPI) prior to robotically-enabled plate-based analysis on an BD FACSCanto II (Stanford Stem Cell Institute FACS Core). Stained cells were not fixed prior to FACS analysis. LEGENDScreen data for undifferentiated H7 hPSCs (day 0) and H7-derived anterior primitive streak (day 1) were published previously^132^. LEGENDScreen data for H1-derived lateral mesoderm (day 2), H1-derived artery ECs (day 3) and H1-derived vein ECs (day 4) was generated in this study. Day 3 artery ECs and day 4 vein ECs were both co-stained with an anti-CD144 Alexa Fluor 647 antibody (BD Biosciences, 561567) to identify CD144+ ECs, and surface-marker expression was evaluated specifically in the CD144+ population.

### Assembling CRISPRi constructs

sgRNAs targeting the human *SOX7*, *SOX17*, or *SOX18* genes were selected from genome-wide libraries of CRISPRi sgRNAs^138^.

- *SOX7* CRISPRi sgRNA 1: TCGCCTCGCTTCGCCTGGCG
- *SOX7* CRISPRi sgRNA 2: GAAGCGAGGCGACCCGCGTG
- *SOX17* CRISPRi sgRNA: GCGACAGGCCAGAACACGGG
- *SOX18* CRISPRi sgRNA: GCGGATGGCGGTGGGGACGG

To prepare sgRNA inserts, we synthesized the following oligonucleotides in preparation for introduction into the single sgRNA plasmid (harboring BstXI and Bmtl restriction sites):

- 5’ ttg + top strand sgRNA + gtttaagagc 3’
- 5’ ttagctcttaaac + bottom strand sgRNA + caacaag 3’

To prepare sgRNA inserts, we synthesized the following oligonucleotides in preparation for introduction into the dual sgRNA plasmid (harboring BstXI, BsmBI, and BlpI restriction sites):

- 5’ ttg + position A top strand sgRNA + tctca 3’
- 5’ ctcttgaga + position A bottom strand sgRNA + caacaag 3’
- 5’ gaaaggag + position B top strand sgRNA + gtttaagagc 3’
- 5’ ttagctcttaaac + position B top strand sgRNA + ctcc 3’

These oligonucleotides were annealed using 20 μL of each oligonucleotide at 100 μM concentration + 10 μL of 10x annealing buffer (100 μM Tris HCl (pH 7.5), 500 mM NaCl, 10 mM EDTA (pH 8.0)) + 50 μL water. Annealing was performed at 99 °C for 5 minutes and then brought to 25 °C for 5 minutes in a thermal cycler.

To individually knock down each of these genes, we cloned each individual sgRNA into a separate *mU6-sgRNA; EF1A-HygroR* plasmid using the Quick Ligation Kit (New England Biolabs, M2200), generally using the manufacturer’s protocol but with modified reaction volumes: 0.33 μL digested plasmid at 25 ng/μL concentration, 1.67 μL 2x quick ligase buffer, 0.166 μL quick ligase, and 1.33 μL annealed sgRNA at 20 nM concentration per reaction.

In experiments where we sought to simultaneously knockdown all three genes, we assembled a dual sgRNA plasmid using a previously-described strategy^100^. A dual sgRNA plasmid was assembled from two backbone fragments (Addgene, 187243 and 187239, respectively). We assembled a *mU6-SOX7-sgRNA 1; hU6-SOX17-sgRNA; EF1A-PuroR-2A-*GFP plasmid using the Quick Ligation Kit (New England Biolabs, M2200). The manufacturer’s protocol was generally followed, but modified reaction volumes were used: 0.33 μL of backbone plasmid 1 at 25-33 ng/μL concentration, 0.33 μL of backbone plasmid 2 at 25-33 ng/μL concentration, 1.67 μL 2x quick ligase buffer, 0.166 μl quick ligase, 0.5 μL of annealed sgRNA 1 at 200 nM concentration per reaction, and 0.5 μL of annealed sgRNA 2 at 200 nM concentration per reaction. As described below, to achieve triple *SOXF* knockdown, we transduced CRISPRi hPSCs with a *mU6-SOX18-sgRNA*; *EF1A-HygroR* plasmid, and subsequently transduced them with this dual *SOX7/SOX17* sgRNA construct.

All plasmids were transformed into Mix & Go! *E. coli* Competent Cells (Zymo Research) and purified using the Wizard Plus SV Minipreps DNA Purification System (Promega).

### CRISPRi knockdown in hPSCs

CRISPRi plasmids were packaged into VSV-G pseudotyped lentiviruses in HEK293T/17 cells using a 3^rd^ generation lentiviral packaging system as described previously^48^. 24 hours prior to transfection, HEK293T/17 cells were seeded at a density of 105,000 cells/cm^2^ in 6-well plates coated with 0.01% poly-L-Lysine. Each well was transfected with 1.39 μg pMDL plasmid + 0.78 μg VSV-G plasmid + 0.53 μg pREV plasmid + 11.3 μL FuGENE HD transfection reagent + 2.1 μg sgRNA plasmid in Opti-MEM medium. 18 hours post-transfection, media was changed to DMEM with 10% FBS. Supernatant was collected 42 hours post-transfection, filtered with 0.45 μM polyethersulfone filter, and stored at -80 °C until used for transduction.

In parallel, H1 hPSCs were engineered to constitutively express CRISPR interference (CRISPRi) machinery, namely nuclease-dead Cas9 (dCas9) fused to the transcriptional repressor ZIM3 KRAB^72^. Validation of these CRISPRi-expressing H1 hPSCs will be described in a forthcoming manuscript.

To transduce them with lentiviruses carrying single sgRNAs, H1 CRISPRi hPSCs were dissociated with Accutase, and then re-seeded at 150,000 cells/well in 6-well plates in StemFlex (Thermo Fisher, A3349401) supplemented with 1% penicillin/streptomycin and 1 μM thiazovivin and lentivirus-containing supernatant (15 μL for *SOX17*-sgRNA, 10 μL for *SOX7*-sgRNA, and 50 μL for *SOX18*-sgRNA lentiviruses, respectively). After reaching 80% confluency after 2-3 days, cells were re-plated using Accutase and cultured in StemFlex with Hygromycin B (Thermo Fisher, 10687010) at 50-100 μg/ml concentration for at least 3 weeks. 1 μM thiazovivin was supplemented for the first day of selection.

To generate the triple *SOXF* knockdown line, first we transduced H1 CRISPRi hPSCs with the *mU6-SOX18-sgRNA*; *EF1A-HygroR* lentivirus and performed hygromycin selection for at least 3 weeks, as described above. These H1 CRISPRi *SOX18-*sgRNA hPSCs were then dissociated with Accutase, re-seeded at 200,000 cells/well in 12-well plates in StemFlex supplemented with 1% penicillin/streptomycin, 50 μg/ml Hygromycin B and 1 μM thiazovivin and transduced with 50 μl of lentivirus-containing supernatant. The following day, media was replaced with StemFlex supplemented with 1% penicillin/streptomycin and 50 μg/ml Hygromycin. After 3 days, cells were re-plated using Accutase and selected with 1 μg/ml Puromycin (Thermo Fisher, A1113803) and 50 μg/mL Hygromycin B for at least 3 days. 1 μM thiazovivin was supplemented for the first day of antibiotic selection.

H1 CRISPRi hPSC lines were then further differentiated into artery and vein ECs using the V1 protocol described above. qPCR and scRNAseq was performed on control and *SOXF* knockdown artery ECs, primed ECs, and vein ECs as described above.

### Construction of *SOX17-2A-mPlum; NR2F2-2A-GFP* reporter hPSC line

The Cas9/AAV6 knock-in strategy^139^ was used to generate a double *SOX17-2A-mPlum*; *NR2F2-2A-GFP* knock-in reporter hPSC line. We previously generated single *SOX17-2A-mPlum* or *NR2F2-2A-GFP* reporter hPSC lines^48^. In this study, starting from a *NR2F2-2A-GFP* reporter line (clone 10), we used our previously-described approach to knock-in a *2A-mPlum* cassette downstream of the endogenous *SOX17* gene such that the *SOX17* stop codon was removed^48^.

### Regulatory and ethical oversight

All animal procedures at Stanford University were approved by Stanford’s Administrative Panel on Laboratory Animal Care (APLAC). All animal procedures at Oxford University were approved by a local ethical review committee at Oxford University and licensed by the UK Home Office. All hPSC experiments at Stanford University were approved by Stanford’s Stem Cell Research Oversight (SCRO) committee.

## QUANTIFICATION AND STATISTICAL ANALYSES

Statistical tests are specified in the figure legend accompanying each experimental result. Unpaired t-tests were used to test for statistical significance in qPCR and ELISA data, and P values were reported. Wilcoxon rank sum tests were performed to test for statistical significance in module score differences quantified by scRNAseq, and P values were reported. The following convention was used in figures: not significant (n.s., P>0.05), *P<0.05, and **P<0.01.

Q-values were reported for differentially-expressed genes detected by bulk-population RNA-seq, differentially-accessible genomic loci detected by Omni-ATAC-seq, and transcription factor motif enrichments detected by Omni-ATAC-seq. The q-value represents the P value adjusted for multiple hypothesis testing using the Benjamini-Hochberg method to control the false discovery rate.

Quantification of mouse embryo images was performed as described in the section “Quantification of mouse embryo images”.

All *in vitro* experiments entailed two or more technical replicates, defined as two or more wells of cells subjected to the same culture conditions in the same experiment. The only exceptions were experiments that entailed single-cell RNA-seq and LEGENDScreen profiling; such profiling was performed on one technical replicate. For all qPCR experiments, two separate measurements were made of each technical replicate (i.e., for each well of cultured cells, qPCR for each given gene was performed twice in the same 384-well qPCR plate).

For mouse embryo experiments, the following numbers of embryos were analyzed per experiment, and one representative image is shown:

- **E8.5-E8.75 mouse embryos, stained for Erg and Sox17 (Fig. 4C-D**, **Fig. S8C**): N=5 embryos
- **E9.5 mouse embryos, stained for Erg and Sox17 (Fig. 4E)**: N=3 embryos
- **E9.5 mouse embryos, stained for *Aplnr*, Erg, and Sox17 (Fig. 4F)**: N=9 embryos
- **E12.5 mouse embryos, *Sox17-Cre; Aplnr-DreER* intersectional lineage tracing (Fig. 4H)**: N=3 embryos
- **E12.5 mouse embryos, *Sox17-Cre* lineage tracing** (**Fig. S8E**): N=3 embryos
- **E9.75 mouse embryos, *Aplnr-CreER* lineage tracing** (**Fig. S8F**): N=7 embryos
- **Quantification of E9.5 mouse embryos, *Aplnr-CreER* lineage tracing** (**Fig. S8G**): N=4 embryos. Images of 6 regions of interest (ROI) were quantified per embryo.

No sample size calculation, randomization, or investigator blinding were performed.

## SUPPLEMENTAL INFORMATION

**Table S1: Single-cell RNA-seq of hPSC-derived cell-types**

Differentially expressed genes that distinguish day 0 hPSCs (“H1”), day 1 mid primitive streak (“d1ps”), day 2 lateral mesoderm (“d2dlm”), day 3 artery ECs (“d3aus”), day 3 primed ECs (“d3pvus”), day 4 vein ECs (“d4vus”), and day 3-4 mesenchyme (“mes1”). Gene expression was measured by single-cell RNA-seq.

**Table S2: Bulk RNA-seq of hPSC-derived cell-types**

Count matrix of gene expression of day-0 hPSCs (“H1”), day-1 mid primitive streak (“D1_PS”), day-2 lateral mesoderm (“D2_DLM”), day 3 artery ECs, CD144+ FACS-purified (“D3_Artery”), day 3 primed ECs, CD144+ FACS-purified (“D3_Prevein”), and day 4 vein ECs, CD144+ FACS-purified (“D4_Vein”). Gene expression was measured by bulk-population RNA-seq, and is quantified in log_2_ counts per million (CPM) units.

**Table S3: Cell-surface marker screening of hPSC-derived cell-types**

For each hPSC-derived cell-type, the number indicates the percentage of cells that expressed a given cell-surface marker by high-throughput flow cytometry. Day 3 artery EC and day 4 vein EC populations were pre-gated on the CD144+ endothelial subset before quantifying the expression of other cell-surface markers. This was done to avoid being confounded by CD144-non-endothelial cells in the culture.

**Table S4: Single-cell RNA-seq of E9.5 mouse embryo ECs *in vivo***

Differentially expressed genes that distinguish 4 different types of ECs identified in the E9.5 mouse embryo. Raw single-cell RNA-sequencing data were obtained from a previous study^140^. We provisionally annotated the following clusters based on expression of characteristic marker genes: 0 (vein-like 1 ECs), 1 (other ECs), 2 (artery ECs), and 3 (vein-like 2 ECs).

**Table S5: Single-cell RNA-seq of gene expression changes upon triple *SOXF* knockdown during artery EC differentiation**

Genes that are differentially expressed between artery ECs generated from control vs. *SOX7/SOX17/SOX18*-CRISPRi hPSCs. Positive fold change indicates higher expression in control artery ECs, and log_2_ fold change is shown. Gene expression was measured by single-cell RNA-seq.

**Table S6: Single-cell RNA-seq of gene expression changes upon triple *SOXF* knockdown during vein EC differentiation**

Genes that are differentially expressed between vein ECs generated from control vs. *SOX7/SOX17/SOX18*-CRISPRi hPSCs. Positive fold change indicates higher expression in control vein ECs, and log_2_ fold change is shown. Gene expression was measured by single-cell RNA-seq.

**Table S7: Differentially accessible genomic elements in hPSC-derived artery vs. vein ECs, as assayed by OmniATAC-seq**

Differentially accessible genomic elements in hPSC-derived artery vs. vein ECs, as assayed by OmniATAC-seq. Fold change indicates accessibility in vein ECs relative to artery ECs (i.e., a positive value indicates that a genomic element is more accessible in vein ECs, whereas a negative value indicates greater chromatin accessibility in artery ECs). The gene closest to each genomic element is also indicated.

**Table S8: Transcription factors enriched in artery-specific regulatory elements, as assayed by OmniATAC-seq**

Transcription factor motifs enriched in artery-specific regulatory elements, as determined by HOMER. Motifs are named in accordance with the most closely matching motif in the HOMER dataset. P value indicates the statistical significance of motif enrichment in artery-specific regulatory elements, relative to vein-specific regulatory elements.

**Table S9: Transcription factors enriched in vein-specific regulatory elements, as assayed by OmniATAC-seq**

Transcription factor motifs enriched in vein-specific regulatory elements, as determined by HOMER. Motifs are named in accordance with the most closely matching motif in the HOMER dataset. P value indicates the statistical significance of motif enrichment in vein-specific regulatory elements, relative to artery-specific regulatory elements.

## Notes

### Summary of Updates

Edits to main text, figures, and authorship.

